# Individualized AI-driven Neuromodulation selectively enhances and reorganizes human tongue sensorimotor networks

**DOI:** 10.64898/2026.11.04.764393

**Authors:** T.D. Papageorgiou, J. Webb, A. Allam, S. Reddy, C. Noorbakhsh, D. Huynh, R. Hekmati, E.M. Rohren, E.M. Sturgis, S.G. Hilsenbeck, K.A. Hutcheson, S.R. Heilbronner, A.M. Thompson, C.R. Neblett, E. Froudarakis

## Abstract

Precise modulation of brain networks for tongue motor and sensory control (TMSC) is critical for neruorehabilitation. Existing neuromodulation approaches use fixed targets, ignoring individual network architecture. We present an AI-driven, individualized fMRI neuromodulation (iNM) platform adaptively targeting subject-specific TMSC networks in real-time. Using a neurofunctional twin built from each participant’s anatomy and function, iNM reinforces targeted activity and suppresses competing states. Unsupervised machine learning and hidden Markov modeling showed iNM increased TMSC-state occupancy from 82% to 97% and suppressed swallow-state interference by 83%. The inferior parietal lobule emerged as a sensorimotor integration hub, with increased magnitude (d′ 90.4 right, 84 left) and spatial expansion (134.8% right, 81.8% left). Cortico-subcortical connectivity strengthened 53% to 74% across motor, sensory, and attention networks and persisted into non-iNM blocks. Thus, iNM may offer precision neurorehabilitation for CNS or PNS disorders, such as amyotrophic lateral sclerosis, and treatment-induced cranial neuropathy in head-and-neck-cancer survivors, respectively.

## Introduction

Interoception and proprioception allow the nervous system to monitor internal physiological states and coordinate movement. Both are critical for tongue motor and sensory control (TMSC)^1–5^, which supports swallowing, speech, and eating through coordinated activity of cranial nerves IX, X, and XII, and their cortical representation in primary sensorimotor cortices. Disruption of TMSC, particularly due to damage to these nerves, is a hallmark of chronic cranial neuropathy (CCN), commonly resulting from surgery or radiotherapy for head and neck cancers^6–15^. CCN impairs speech, swallowing, and mastication through progressive atrophy and fibrosis, substantially diminishing quality of life^10,14,16^. Despite increasing survivorship, approximately 10% of head and neck cancer survivors develop CCN^17–19^, for which pharmacologic treatments are poorly tolerated and offer no practical benefit^20,21^. TMSC dysfunction also presents early in neurodegenerative conditions, such as amyotrophic lateral sclerosis (ALS), where up to 80% of patients develop dysarthria and dysphagia, culminating in respiratory failure and premature death^22,23^. In both CCN and ALS, tongue weakness is accompanied by cortical disorganization reflecting degraded network precision and excitability.^24–26^ This manifests as sensory–motor “smudging,” a marker of de-differentiated topographic organization within primary sensorimotor areas that degrades the precise representations required for selective tongue control.^27–29^

Two rationales motivate cortical targeting as a strategy for restoring TMSC. First, peripheral nerve damage in CCN compromises tongue elevation, closure, and base retraction, which places increased demands on cortical compensation. Second, motor cortex stimulation can modulate sensorimotor function, albeit with limitations^30–36^. Non-invasive brain stimulation approaches, such as transcranial magnetic stimulation and transcranial electrical stimulation are constrained by poor spatial focality, limited depth of penetration that precludes access to deep and subcortical TMSC nodes, anatomically fixed targeting that ignores individual functional topography, and open-loop (fixed) delivery ^37,38^. These limitations underscore the need for neuromodulatory strategies capable of selectively engaging circuit-specific sensorimotor networks with high spatial and temporal precision and safety^39–44^.

Recent advances in precision neurotechnology have integrated high-resolution neural interfaces, specifically noninvasive holographic imaging^45^ and invasive AI-driven digital twin models in animal models^46,47^ to enable patient-specific neural intervention in real time. These approaches capture neural activity with high spatial and temporal resolution and can thus guide targeted intervention. This coupling of precision imaging and adaptive control bridges neuroscience and clinical practice, offering a pathway from trial-and-error approaches toward spatiotemporally precise, individualized neurotherapies.

Realizing this potential, requires a system capable of targeting each individual’s unique neural architecture with sufficient precision to guide network-specific recruitment, while remaining sensitive enough to capture effects that generalize across individuals. In conceptual resemblance to digital twin models of the brain, our real-time, fMRI-based, AI-driven closed-loop neuromodulation system establishes a high-fidelity, individualized representation of a brain network (the neurofunctional twin), whose activation is continuously updated in real-time using neurophysiological data to reflect each participant’s dynamic brain-level responses to a specific function. Unlike prior real-time fMRI neurofeedback approaches that either feedback single- or dual-ROI signals from anatomically- or atlas-defined regions ^48,49^, or group-derived network parcellations into individual space ^50,51^, iNM delineates each participant’s TMSC network from the intersection of their own functional activation and anatomy, and modulates this distributed cortical–subcortical network as a dynamically updated closed-loop target (the neurofunctional twin). This neurofunctional twin is derived from each individual’s combined anatomical and functional networks, whose spatially precise and temporally resolved activity is used to dynamically guide iNM-based modulation and optimize performance.^52^ **In contrast to the invasive animal digital-twin models noted above, the neurofunctional twin is non-invasive, built from each human participant’s own anatomy and function, and operates in a closed loop at 2 s resolution.**

The iNM platform integrates several capabilities that collectively distinguish it from existing neuromodulation approaches. It targets each participant’s unique cortical and subcortical networks involved in TMSC with a native voxel size of 3.125 × 3.125 × 3.0 mm, enabling simultaneous engagement of distributed structures, including primary sensorimotor cortices, basal ganglia, thalamus, cerebellum, and insula, that cannot be directly accessed by surface-based neuromodulation tools (Fig. 2). The platform is individualized at both anatomical and functional levels, tailored to each participant’s neural organization and physiology. It operates in a spatiotemporally guided, closed-loop fashion, identifying time-resolved signal changes in real time and reinforcing specific neural patterns, for example enhancing motor network activity during tongue movement or suppressing competing swallow-related interference, to optimize the physiological response under investigation. The system is adaptive, continuously tracking each participant’s evolving TMSC brain network dynamics and promoting balanced bilateral activation (Fig. 1). As the individualized network is repeatedly engaged during the TMSC physiological response, the system progressively adjusts activation thresholds, promoting more precise and efficient recruitment within the targeted network, a mechanism reflected in the increased signal-to-noise ratio and reduced BOLD variance observed under iNM (see Results). Despite individualized targeting, consistent patterns of TMSC network reorganization emerged across participants, revealing a common architecture of sensorimotor control. This positions iNM at the intersection of precision neurorehabilitation and mechanistic neuroscience. Notably, modulation efficacy is not defined by BOLD amplitude alone, but also by the adaptive adjustment and spatial expansion of the individualized network’s functional response. The platform thus serves as a modulation tool during real-time closed-loop operation and, through offline machine learning analysis, a decoding platform that characterizes each participant’s unique neural architecture during TMSC.

**Fig. 1.**
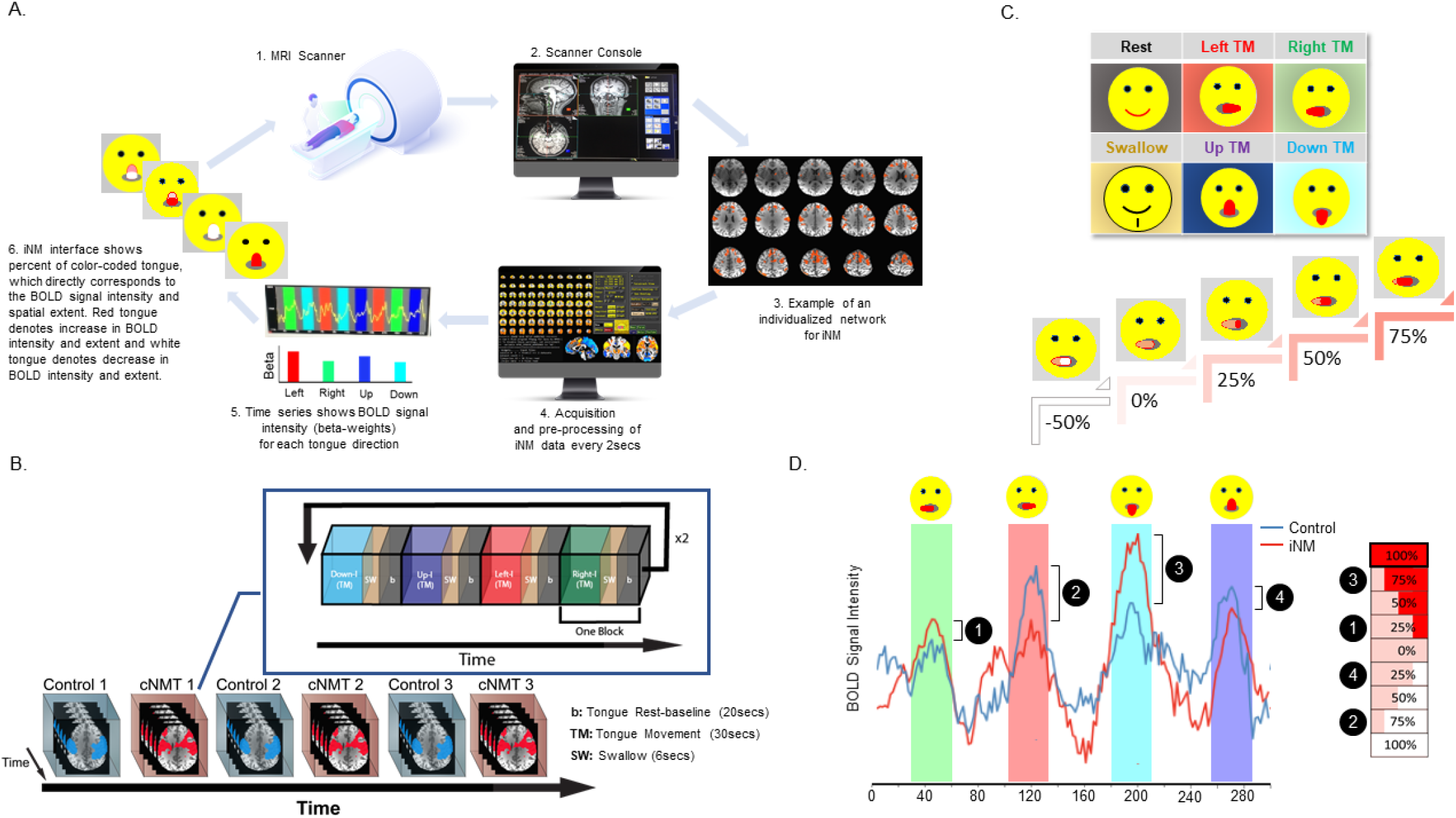
Individualized AI-driven neuromodulation (iNM) **A. 1. & 2**. High-resolution anatomical images were acquired via MRI and registered directly to the Siemens console system. **3**. For each participant, TMSC networks were delineated by mapping direction-specific BOLD responses (left, right, up, down), defined by increased signal intensity and spatial extent associated with tongue movement.**4**. In real-time (Time repetition - TR = 2 s), iNM data from each participant’s individualized network yellow trace represents the time series, with the baseline (tongue at rest) shown in grey, and direction-selective cortical responses for left, right, up, and down shown in red, green, blue, and cyan, respectively. **5**. BOLD signal intensity and spatial extent for each direction were computed using a general linear model (GLM); beta-weights were iteratively updated to define each participant’s individualized TMSC network (see Methods). **6**. The iNM interface visualizes the dynamic modulation of TMSC networks every 2 s, with red fill indicating percent upregulation, pink indicating baseline, and white fill indicating downregulation of BOLD signal intensity and spatial extent, mapped onto a tongue schematic. **B**. Scans were acquired in an interleaved sequence (Control Scan 1, iNM Scan 1, Control2, iNM2, Control3, iNM3). The temporal sequence of physiological tasks was identical across iNM and control (No-iNM) scans: each TMSC direction block lasted 30 seconds, interleaved with 6-second swallow blocks, and 20-second baseline (tongue-at-rest) intervals. Tongue movement directions were counterbalanced across runs, and same direction blocks were never presented consecutively. **C**. iNM interface. During control scans, the entire tongue was filled red to encourage maximal upregulation of the target network for each tongue direction. During iNM scans, the emoji tongue adapted in real time (2 secs) to reflect each participant’s individualized TMSC-selective network activity. Percent BOLD magnitude and spatial extent determined the fill color: red (25–100%) indicated upregulation, pink (0%) represented baseline, and white (−25% to −100%) denoted downregulation. This visualization directly reflected each participant’s dynamic neural engagement. **D**. Computation of the iNM signal. Percent BOLD signal change (PSC) during TMSC-selective periods was computed at each TR (2 s) relative to the baseline-tongue-at-rest block preceding each TMSC direction within each control scan. The current percent BOLD PSC was compared against a graded scale of 25% increments (−100%, −75%, −50%, −25%, 0, 25%, 50%, 75%, 100%). To increase task demand during iNM relative to control, this scale was shifted upward by 10%, such that a BOLD level that filled the tongue red during control produced a lesser red fill during iNM, requiring participants to generate stronger network upregulation to reach the same fill. Positive values (25% to 100%) produced a proportional red fill (upregulation), 0% produced a pink fill (baseline), and negative values (−25% to −100%) produced a proportional white fill (downregulation), as defined in panel C.

**Fig. 2.**
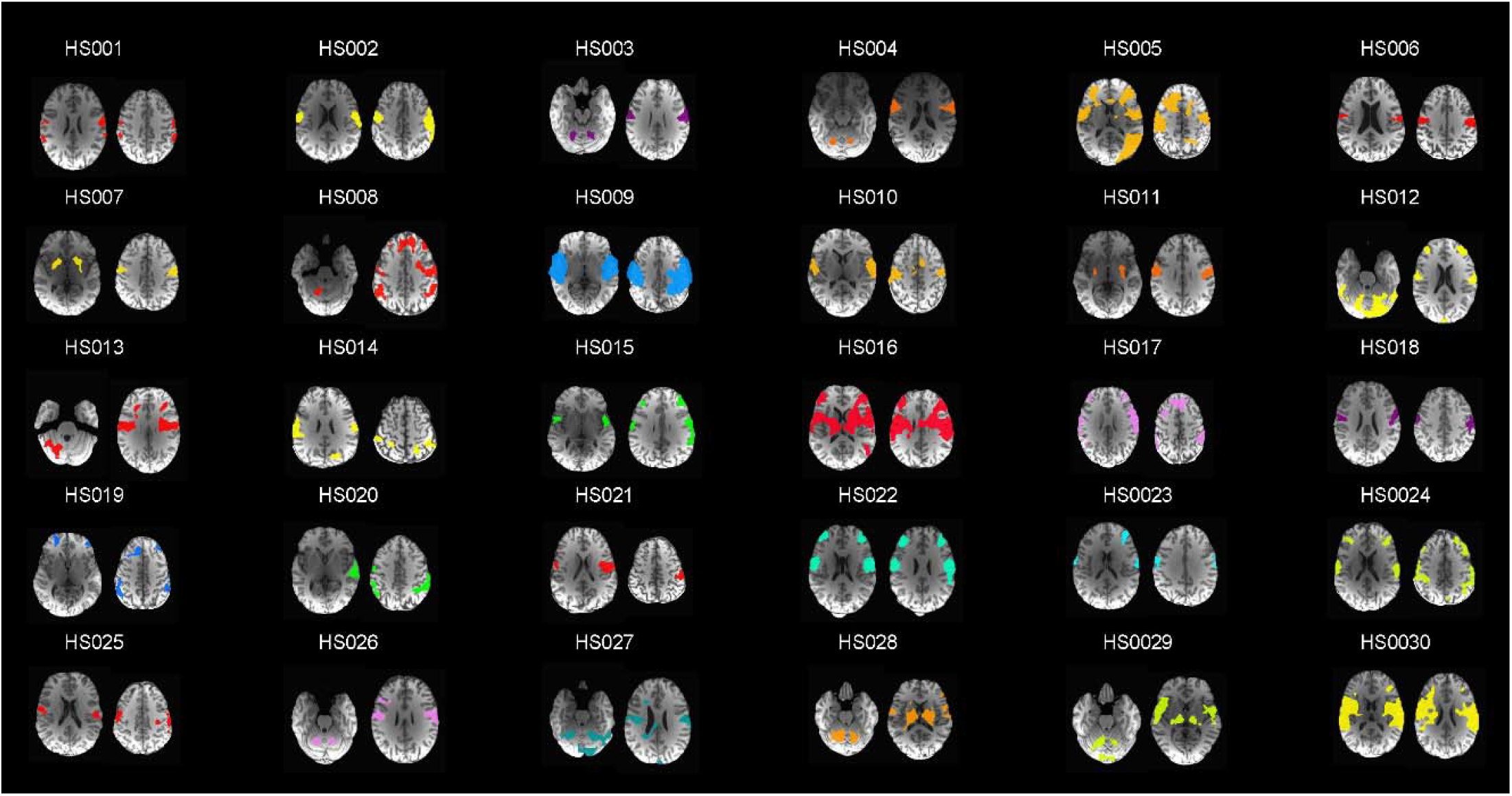
Individualized networks for iNM (n=30). Optimization and efficacy of iNM rely on precisely delineating the intensity and extent of the cortical network based on each participant’s unique anatomy during tongue movement. TMSC localization was performed by identifying motor and sensory networks exhibiting maximal upregulation across directions using a general linear model (GLM), followed by computing the intersection of these functional activations with anatomical regions. Each individualized network was then optimized by excluding noise-related signals — for example, BOLD activity extending into cerebrospinal fluid, limited to a single slice, or lacking spatial continuity. Each resulting individualized network constitutes the participant’s neurofunctional twin, which served as the target for iNM.

To establish the feasibility and mechanistic specificity of individualized neuromodulation, we applied iNM to the TMSC network, which engages well-defined, bilateral motor and sensory circuits at the cortical and subcortical levels with direct clinical relevance to CNS and PNS disorders, making it ideally suited for this proof-of-concept of adaptive, circuit-level intervention. We demonstrate that iNM selectively enhances motor, sensory, and attention networks underlying tongue control by increasing signal reliability, spatial extent of recruitment, and temporal stability while suppressing competing physiological states, such as swallowing-related interference. By integrating real-time fMRI, machine learning, and temporal causal modeling via the autoregressive hidden Markov model, we show that targeted, individualized modulation can reconfigure sensorimotor network dynamics persistently beyond the neuromodulation window. These findings provide a mechanistic foundation for precision-guided neurorehabilitation strategies and targeting TMSC impairments across the neuroaxis. By engaging CNS circuits in neurodegenerative disease and PNS pathways in cranial neuropathy, iNM addresses conditions for which effective treatments remain limited.

## Results

### iNM enhances the decodability and directional selectivity of TMSC representations

To elucidate the mechanisms of iNM, we first quantified how well the direction of tongue movement could be predicted from fMRI activity using linear support vector machine (SVM) analysis. SVM-generated cortical maps (Fig. SI-1) demonstrate enhanced signal intensity and spatial recruitment under iNM, reflecting greater directional selectivity during tongue movement tasks (Figs. 3, SI-3). This enhancement, quantified by the area under the curve (AUC), demonstrates greater consistency and reliability of the underlying neural signal under iNM (Fig. SI-3). Across all movement directions, iNM increased both classification accuracy and AUC-ROC relative to control. The gains were modest and did not reach statistical significance for the left (iNM 97.2%, 0.994 vs control 95.9%, 0.989) and right (iNM 97.7%, 0.998 vs control 96.0%, 0.988) directions. For all tongue movements combined versus baseline, iNM demonstrated significantly higher mean classification accuracy and AUC-ROC (94.3%, 0.997) than control (89.6%, 0.985; p < 0.0001). Likewise, for the up (iNM 95.9%, 0.991 vs control 90.0%, 0.959) and down (iNM 96.5%, 0.995 vs control 90.8%, 0.964) directions versus baseline, iNM significantly outperformed control in both accuracy and AUC-ROC (p < 0.0001).

**Fig. 3.**
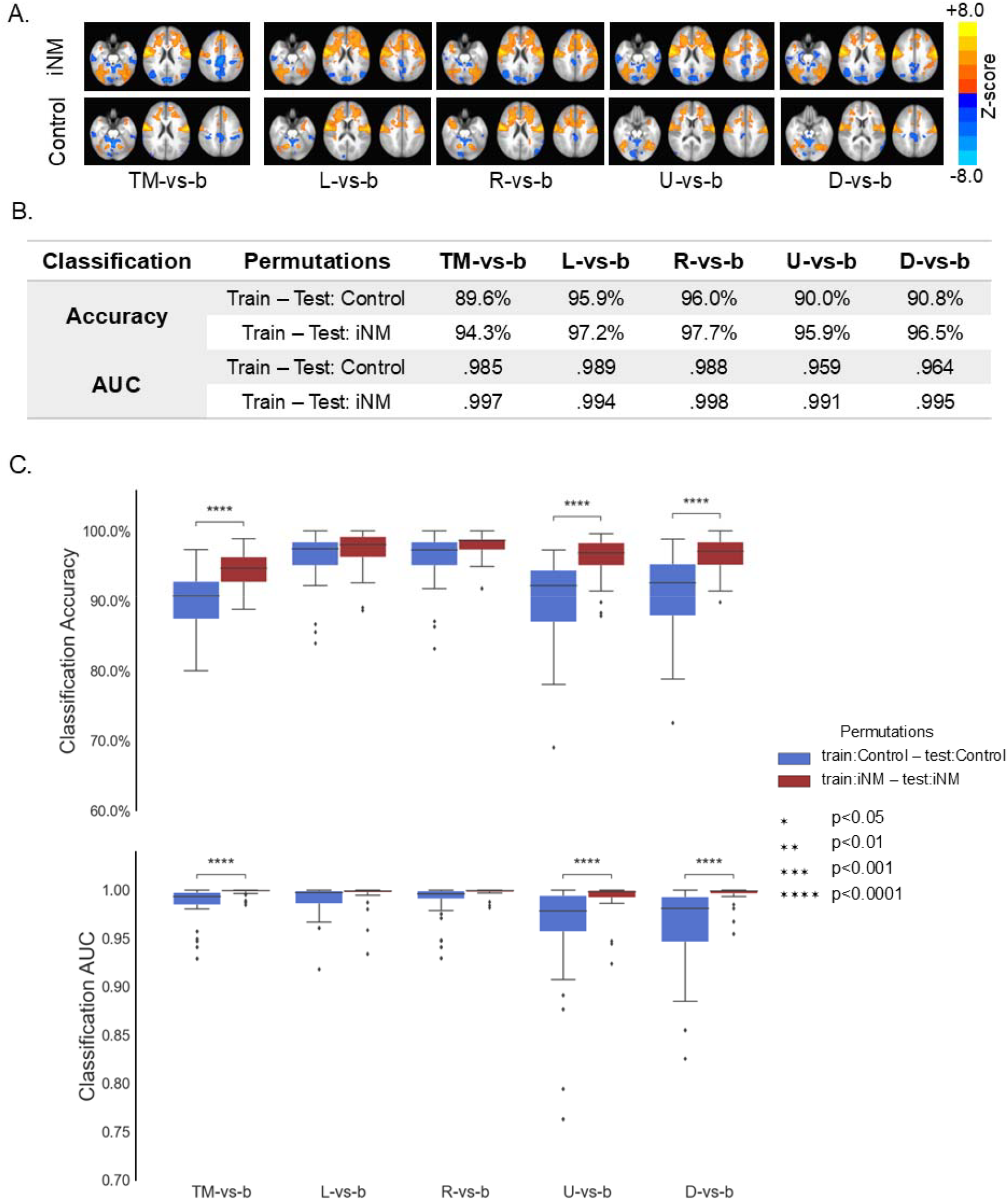
iNM increases consistency in the BOLD intensity and spatial extent compared to the control-no iNM condition, as shown by linear support vector machine (SVM) analysis. **A**. Decoding TMSC cortical networks underlying direction-specific tongue movement. SVM activation maps for each tongue movement direction under iNM (top) and control (no-iNM, bottom) conditions, thresholded voxelwise at significance levels ranging from p < 0.05 to p < 0.0001, with a cluster size > 10 voxels. **B**. SVM-derived classification accuracy and area under the receiver-operating curve (AUROC) across subjects as a function of each tongue movement and across tongue movements versus baseline classification (x-axis), with training-testing permutations indicated by color. **C**. Boxplots of SVM-generated classification accuracies and AUROC across subjects of each tongue movement and across tongue movements versus baseline classification (x-axis), with training–testing permutations indicated by color. L = Left TMSC cortical selectivity during Left tongue movement, R= Right TMSC during Right tongue movement, U = Up TMSC during Up tongue movement, D = Down TMSC during Down tongue movement, TM = TMSC selectivity during Tongue Movement across directions, b = baseline-tongue at rest. Each box displays the median (center line), 25th (Q1) and 75th (Q3) percentiles (lower and upper edges). Whiskers extend to the most extreme non-outlier data points. Black open circles denote outliers, defined as values beyond 1.5×IQR from Q1 or Q3. Black lines and asterisks indicate statistically significant differences determined by paired-sample Wilcoxon signed-rank tests.

### iNM improves BOLD signal magnitude and reduces its variance, as reflected by increased discriminability (d′) and AUC of network-specific ROIs across TMSC conditions over time

The percent change in AUC reflects the modulatory effect of iNM on signal magnitude during each TMSC interval. In the *motor* network, iNM increased mean AUC and d’ scores by 21.7% and 61.6%, respectively, with direction-specific AUC increases of 17% (left), 20% (right), 24% (up), 23% (down) (Figs. 4A, 5A, 5B, SI-6--8) and reduced the AUC in BOLD intensity variance by 9% across TMSC directions (Figs. 4C, SI-8B). In the *sensory* network, iNM increased mean AUC by 46% and d’ scores by 60% across directions with direction-specific AUC increases of 41% (left), 44.5% (right), 45% (up), 51.6% (down) (Figs. 4A, 5A, 5B, SI-6--8). iNM reduced the AUC in BOLD intensity variance by 14% across TMSC directions (Figs. 4, SI-8B). In the *attention* network, iNM increased mean AUC by 37% overall and d’ score of 59% across directions with direction-specific AUC increases of 28% (left), 40% (right), 49% (up), 40% (down) (Figs. 4, 5A, 5B, SI-6--8). iNM reduced the AUC in BOLD intensity variance by 22% across TMSC directions (Figs. 4, SI-8B). In the *reward* network, iNM increased mean AUC by 30%, and d’ score of 63.3% across directions with direction-specific AUC increases of 7.5% (left), 40.9% (right), 45.2% (up), 32.7% (down) (Figs. 4, 5A, 5B, SI-6--8), while it increased the AUC BOLD intensity variance non-significantly across TMSC directions (Figs. 4, SI-8B).

**Fig. 4.**
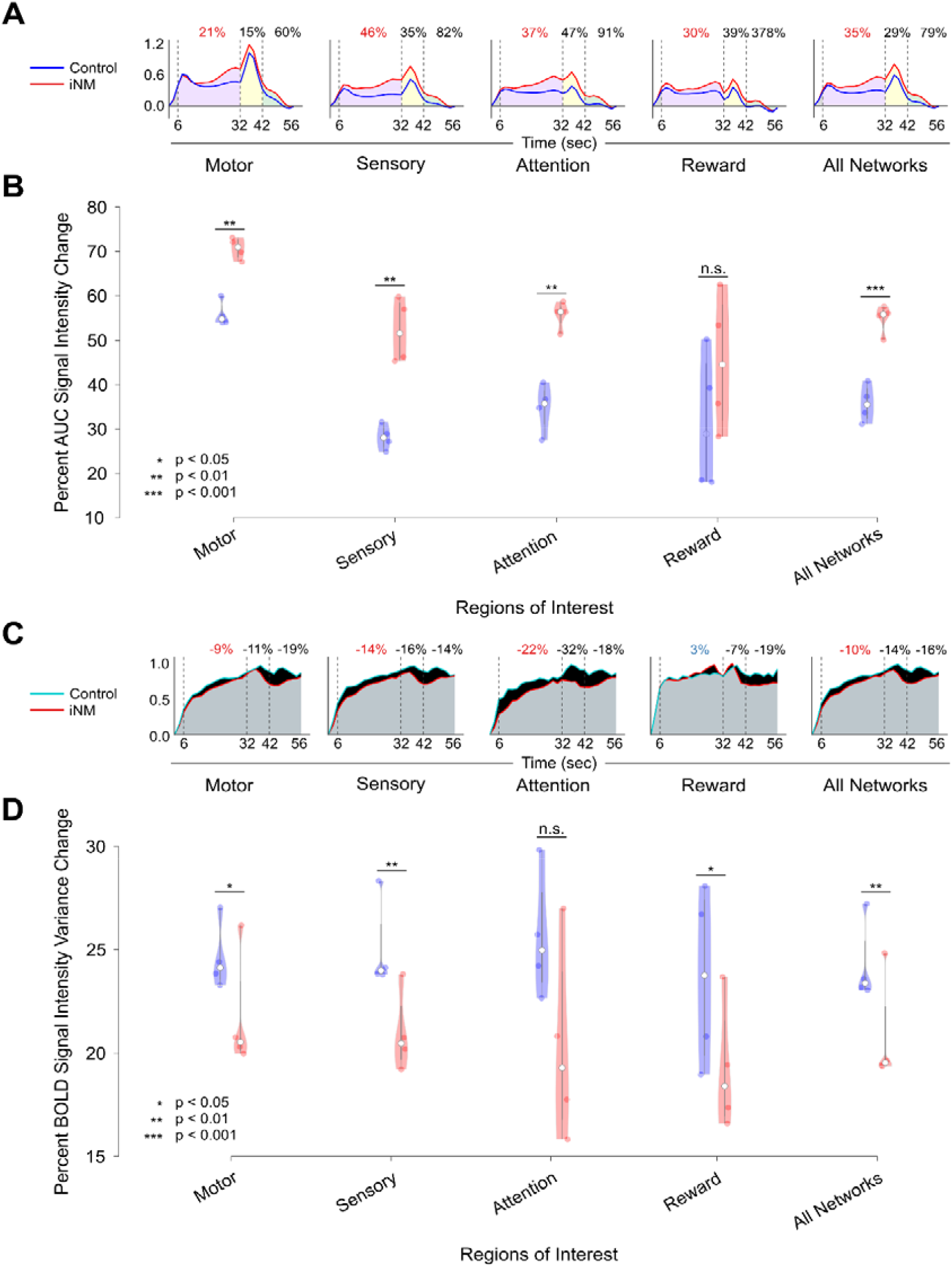
iNM (red) increases BOLD encoding consistency across networks and timepoints. **A**. iNM (red) increased the mean AUC of BOLD intensity and reduced its variance compared to control (no-iNM, blue) across cortical networks and movement directions. Task intervals are shaded: TMSC (purple; 6–32 sec), swallow (yellow; 32–42 sec), and baseline (green; 42–56 sec). Notably, iNM effects persisted into the swallow and baseline periods, when no iNM was applied. Detailed ROI-level results are shown in Figs. SI4a, SI4b, SI5a, SI5b, and SI8. **B**. Across networks and directions (dots indicate direction-specific values), iNM significantly differed from control (p < 0.01) in all networks except reward. **C**. AUCs of BOLD variance, shaded as in A, show that iNM consistently reduced variance across networks compared to control (negative values = lower variance). The reward network was the exception. ROIs that were not part of a given direction’s network were assigned a value of zero; results were consistent when replaced with Gaussian noise (mean = 0, SD = 0.05). iNM reduced the AUC BOLD intensity variance, an effect that also carried over to the non-iNM swallow and baseline blocks. **D**. Variance-based significance testing showed iNM differed from control (p<0.05) across all networks, except attention, where no significant difference was found.

**Fig. 5.**
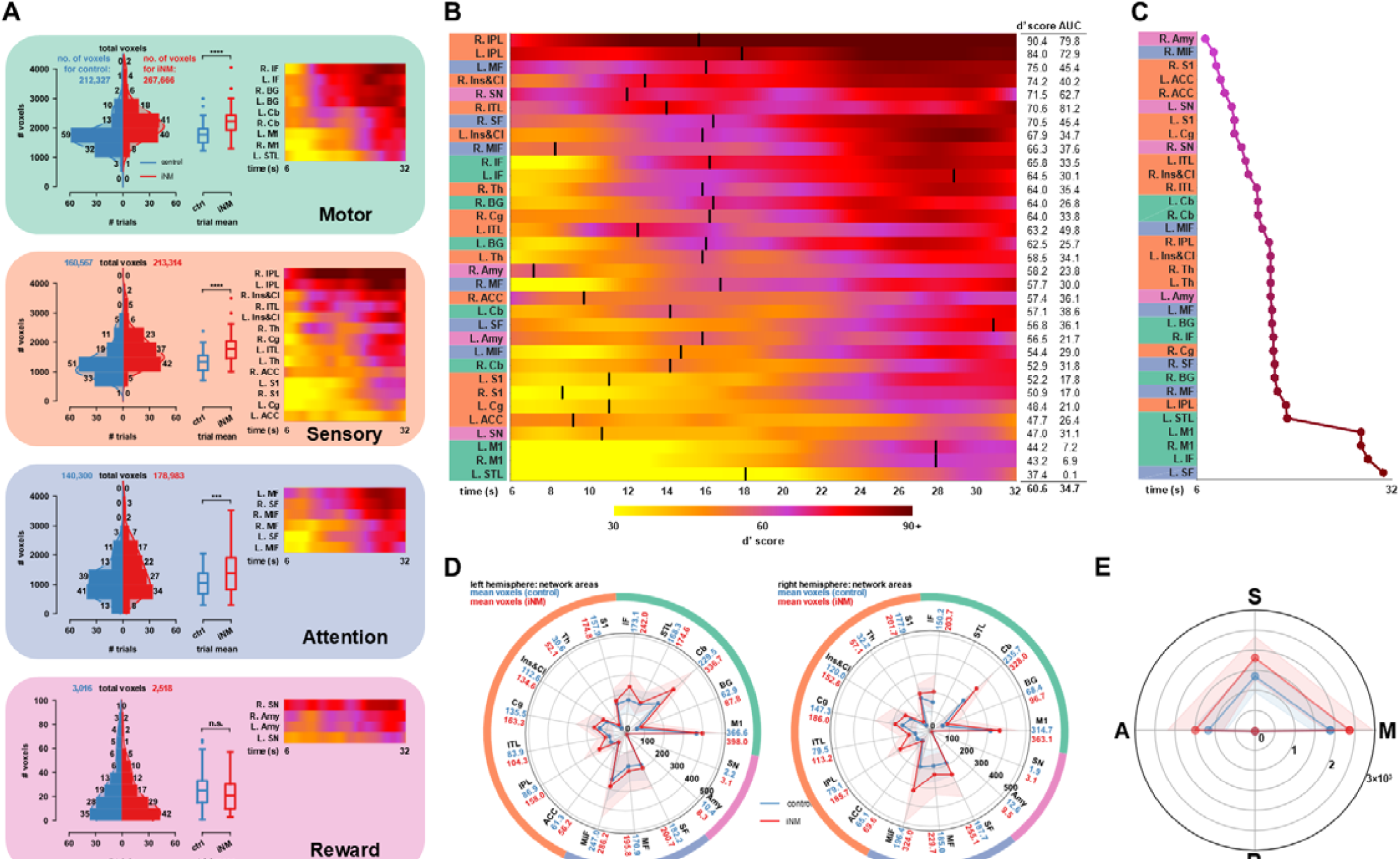
iNM increases the BOLD signal in both magnitude and spatial extent. **A**. (*left*) For each network (motor, sensory, attention, reward), the total number of voxels comprising all ROIs per block interval (by subject and direction) is shown for both control (blue) and iNM (red) conditions. Distributions are visualized via histograms and kernel density estimates (smoothed, non-parametric distribution), with corresponding summaries shown in the adjacent box plots (right; *** is p<0.001, **** is p<0.0001, n.s. not significant). Heat maps display mean□*d*′□scores and percent BOLD signal AUC change for each region, grouped by network. Red (iNM) and blue (control) indicate mean□*d*′□values across all time points and trials. **B**. ROIs from panel A are ranked by descending□*d*′□score. Each ROI is color-coded by network: **motor is green; sensory is orange; attention is purple; reward is pink**. The accompanying table reports mean□*d*′, peak□*d*′, and percent BOLD AUC change during the TMSC interval (6–32□s). The peak d′ (black line) is the earliest peak of the cubic-interpolated d′ time series (MATLAB findpeaks, default parameters; see Methods). **C**. Timing of peak□*d*′□score per ROI, using the same peaks as in panel B, ordered to depict the temporal pathway of network engagement. **D, E**. Voxel counts per ROI in motor (M), sensory (S), attention (A), and reward (R) networks increased under iNM relative to control. Lines and shading denote group mean and standard deviation (s.d.) for iNM (red) and control (blue).

When averaged across networks and directions, AUC increased by 35%. Direction-specific changes in AUC and d′ were: 27% and 60.8% (left), 35% and 62% (right), 38% and 63% (up), and 39% and 58.5% (down), respectively (Fig. SI-8A). During swallow and baseline tongue-at-rest periods—when iNM was not delivered—AUC increases were still observed across directions and networks (except for the reward network in the left direction), suggesting carry-over effects extend their impact beyond active iNM (swallow period: 32-42 secs, and baseline-tongue-at-rest period: 42-56 secs; Figs. 4A, SI-8A, SI-8B). These findings demonstrate that iNM consistently enhanced the magnitude of neural signals within the TMSC network. Increases in the variance of the magnitude under iNM were observed in the right tongue direction for the motor (6%), sensory (1%), and reward (10%) networks, with only a 1% overall increase across all ROIs and networks (Fig., SI-8B). During the swallow period when no iNM was provided (32-42 secs, Fig. 4C), there was a significant decrease in the variance of the BOLD magnitude as quantified by the AUC BOLD intensity variance compared to the control across all directions of motion, and all networks (Figs. SI-8B, 4C). Similarly, during the baseline-tongue at rest period (42-56 secs; Fig. 4C), there was a decrease in BOLD intensity variance, for each and all directions and networks. The relationship between the magnitude of the BOLD and the variance of that magnitude is critical for interpreting the effects of iNM on the tongue motor and sensory cortex (TMSC) network. A decreased variance in BOLD signal intensity, when accompanied by an increased mean signal, reflects an improved signal-to-noise ratio; i.e, the neural response becomes more consistent and reliable, with less trial-to-trial or temporal fluctuation relative to the strength of the TMSC response.^1-^

^3^ Thus, iNM can have a carry-over effect in enhancing TMSC, even when no iNM is applied (Figs. 4, SI-8A—8B). An overall 10% reduction in the AUC BOLD intensity variance indicates improved signal stability and higher SNR under iNM (Figs. 4C, 4D, SI-8B). Against this network-wide ~10% reduction in variance, the bilateral IPL was the notable exception, with BOLD variance rising by 53% relative to control (Fig. SI-8B), a dissociation examined further below.

Spatially, network expansion increased across sensory (32.1%), motor (14.9%), and attention (27.6%) networks, with the largest regional expansions in the bilateral inferior parietal lobule (134.8% in the right IPL, 81.8% in the left IPL) and thalamus (77.3% in the right, 70.3% in the left), underscoring their critical role in sensory gating during tongue movement (Figs. 5B, 5C, 5D). In contrast, the reward network exhibited an 11.4% decrease in spatial expansion.

### Temporal hierarchy of network engagement

To examine the temporal hierarchy of activation, we sorted the peak times of each region and identified the following sequence from earliest to latest (**Fig. 5C**): 1. the right amygdala (7.1 s); 2. attention-related regions, including the right medial and middle frontal gyri (8.2 and 8.4 s); 3. several sensory regions, including bilateral S1 (8.6 and 12.1 s), interleaved with the cingulate cortex (10.5 s) and bilateral inferior temporal lobules (12.7 and 14.0 s); 4. bilateral anterior cerebellar motor regions (14.0 and 14.7 s); 5. additional sensory areas, including the bilateral thalamus, right inferior parietal lobule, and left insula and claustrum (15.7–15.8 s); 6. a mixed motor–attention cluster comprising bilateral basal ganglia, bilateral superior and middle frontal gyri, right cingulate, and right inferior frontal gyrus (16.0–17.1 s); 7. late-peaking left superior temporal and left inferior parietal lobules (18.1 s); and 8. the latest-peaking motor execution regions, bilateral primary motor cortices and left inferior frontal gyrus (27.9–29.0 s), followed by the left superior frontal gyrus (31.4 s). The per-direction hierarchy of peak times is shown in Extended Data Fig. 18. This ordering - from limbic (amygdala) and attentional regions, through sensory integration zones and subcortical motor structures, culminating in primary motor cortex execution is consistent with a hierarchical recruitment sequence in which sensory and integrative tuning precedes motor output during effective TMSC.

### Consistent Strengthening Within and Across Motor, Sensory, and Attention Network Interactions by iNM Across Tongue Motor Directions

To quantify how iNM changes the stability of latent brain states (e.g., TMSC-dominant vs. swallow-interference states), we employed an unsupervised, autoregressive hidden Markov Model (ARHMM),^53^ which can infer the latent brain states and directed functional connectivity from BOLD signals during TMSC. The algorithm identified two dynamic network states (Figs. 6A, 6B, 6D): 1. Dominant state associated with TMSC (Figs. 6A, 6B, 6C); and 2. Non-Dominant state associated with swallowing. In the iNM condition, the dominant brain state that controlled TMSC was upregulated by a 15-percentage-point absolute increase (97% of iNM vs 82% of control trials, 18% relative increase; Fig. 6A), while the dominant state for swallowing during the swallow block was upregulated by a 33-percentage-point absolute increase (83% vs 50% of control trials, 66% relative increase; Fig. 6A) and by a 17-percentage-point absolute increase in the baseline-tongue-at-rest block (97% vs 80% of control trials, 21% relative increase; Fig. 6A). ARHMM analysis revealed that iNM suppressed the non-dominant state from 18% of trials under control to 3% under iNM (15-percentage-point absolute reduction, 83% relative reduction), reflecting reduced swallowing-related interoceptive interference during the TMSC interval. Kernel density estimation (KDE) applied to binned data from each iNM and control block generated a smoothed, non-parametric estimate of the probability distribution, that the largest single bin contained 646 trials under iNM and 356 under control classified as the TMSC-dominant state, with totals of 704 iNM and 587 control trials across all bins (Fig. 6B). Of 720 trials per condition, swallow-associated interference during the TMSC interval occurred in only 16 iNM trials versus 133 control trials, underscoring iNM’s stabilizing effect on TMSC brain states.

**Fig. 6.**
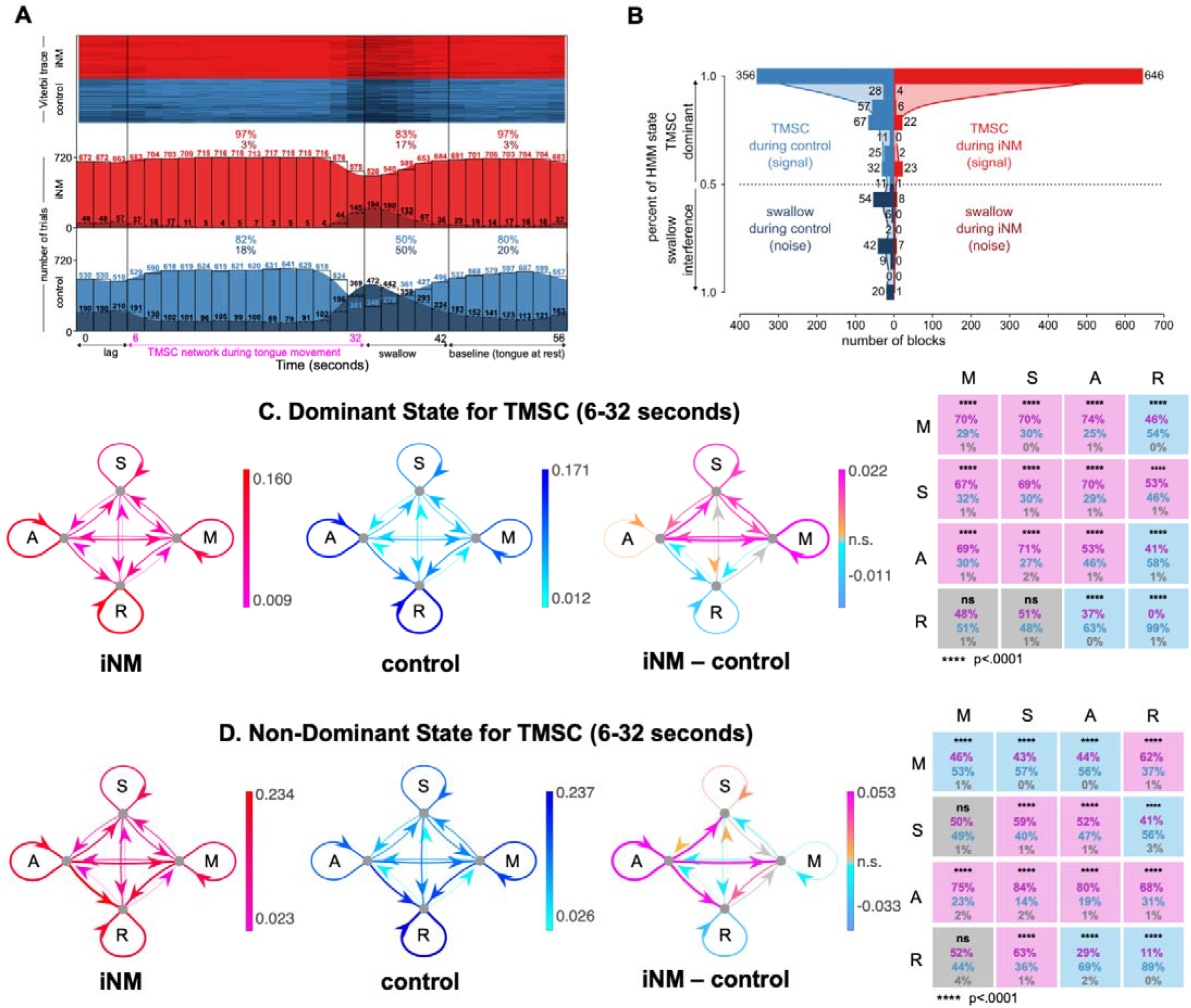
Effect of iNM on network interactions across all tongue motor control directions revealed through temporal causal modeling via unsupervised autoregressive hidden Markov modeling (ARHMM): A. We selected the timeseries from the brain areas that select for tongue movement. Our unsupervised, autoregressive hidden Markov model (ARHMM) was fit to the timeseries from the TMSC-selective regions (6-32 seconds). Model inference was then performed on the entire timeseries (0-56 seconds). Top: Viterbi trace for iNM (red) and control (blue) trials that characterize each brain state during TMSC, swallow, and baseline intervals for each condition. Bottom: Trial-wise histograms show the number of occurrences for each brain state during TMSC (light red/blue) and swallow (dark red/blue), separately for iNM and control conditions.. Percentages indicate the proportion of trials occupying each brain state during their TMSC interval, demonstrating state persistence or transition dynamics beyond the TMSC window. **B**. The histogram compares the signal (TMSC state, red/blue) and noise (swallow state, dark red/dark blue) by displaying the number of trials in each bin that correspond to each of the two states (total trials (iNM vs. control): 704 vs. 587 (TMSC state), 16 vs. 133 (swallow state). Shaded curves represent the kernel density estimate (KDE) fit to the binned data for the iNM and control conditions. The KDE was calculated by the summation of normal distributions centered at each data point, followed by normalization, to generate a smoothed estimate of the histogram bins. KDE bandwidth was estimated via Scott’s method, and boundary corrections at 0 and 1 employed reflection of the data across the boundaries. **C-D**. For the dominant (TMSC) or non-dominant state, the strength of ROI-to-ROI connections within networks (M = Motor network, S = Sensory network, A = Attention network, R = Reward network) is quantified as the mean absolute value of the autoregressive coefficients during the iNM (fuchsia/red) or control (light/dark blue) condition, or the difference in absolute values between conditions (orange/fuchsia and light/dark blue indicate increased/decreased during iNM, respectively). At far right, significant increases (fuchsia), decreases (light blue), or insignificant differences (gray) in these network-to-network interactions are shown for iNM compared to control. Percentages indicate the fraction of region-to-region interactions within each network-to-network connection that are increased (fuchsia), decreased (light blue), or insignificant (gray) in iNM compared to control. Correction for multiple comparisons used an FDR of 0.05.

ARHMM also estimated functional connectivity between ROIs within each network for both dominant and non-dominant states. Significance testing revealed ROI connections that were either upregulated or downregulated under iNM and control conditions (Figs. 6C, 6D). During the dominant state, iNM significantly increased connectivity within and across the motor, sensory, and attention networks, ranging from 53% to 74% across all TMSC-selective regions (p<0.0001; Fig. 6C), whereas reward-network connectivity decreased by 99% under iNM, with the residual 1% not statistically different from control (Figs. 6C, 6D); during the non-dominant state, reward-network connectivity decreased by 89% under iNM (Figs. 6C, 6D). The dominant state predominated across all four directions under iNM versus control: 96% vs 88% for the left (8-percentage-point absolute, 9% relative increase; Fig. SI-9), 96% vs 79% for the right (17-percentage-point absolute, 22% relative increase; Fig. SI-10), 95% vs 78% for the up (17-percentage-point absolute, 22% relative increase; Fig. SI-11), and 97% vs 81% for the down direction (16-percentage-point absolute, 20% relative increase; Fig. SI-12); the competing swallow-associated non-dominant state was correspondingly reduced to 3–5% of iNM trials versus 12–22% under control (all p<0.0001; Figs. 6A, 6B, SI-9–12). Overall, iNM reinforced a motor, sensory, and attention network by increasing within- and across-network connectivity and stabilizing the TMSC-dominant state, thereby strengthening control over tongue movement.

## Discussion

### Individualized targeting of tongue representations

Biological systems are inherently complex, in part due to their nonlinearity, which gives rise to the variability that defines their function^54^. This inherent variability necessitates modeling each system individually to precisely capture subject-specific neural dynamics. The iNM intervention constructs subject-specific functional and anatomical profiles from fMRI data (Fig. 2). This process generates a data-driven representation of an individual’s neural network architecture: a “neurofunctional twin” that serves as a functional instantiation^55^ across the iNM scans, crucial for the regulation of each individual’s unique TMSC representation. First, iNM operates through closed-loop adaptation, using iterative modulation within the fMRI environment to update individualized targets based on real-time cortical and subcortical responses. Second, iNM is designed for prediction and optimization: the individualized functional maps predict which cortical and subcortical circuits will respond most selectively to modulation, while real-time BOLD guidance optimizes the timing, intensity, and network recruitment. This mirrors how digital twins continuously refine interventions by adaptively updating predictive models against real-world system behavior. Third, iNM operates as a hemodynamically coupled intervention, relying on the MRI’s magnetic field and BOLD signal to measure oxygenation-dependent, quantifiable parameters that directly guide modulation of brain activity. The scanner’s magnetic field interacts with the individual’s vascular and metabolic landscape, dynamically altering local magnetic susceptibility^56^. iNM guides these subject-specific anatomical and functional cortical and subcortical networks by promoting greater selectivity for TMSC precision, bilateral recruitment, and stability. A key clinical implication of this individualization is that iNM could, in principle, engage lesion-specific mechanisms according to where the pathology lies along the neuraxis. For supranuclear lesions, as in the context of neurodegenerative disease, we hypothesize that iNM may strengthen intact cortical and subcortical circuits to bypass damaged pathways and restore lost functions, which could be a mechanism consistent with the enhanced network magnitude, spatial extent, and stability observed here. For infranuclear lesions, as in the context of head and neck cancer or other neurological damage after surgery, iNM may instead augment top-down cortical drive to partially denervated targets, providing a rationale for testing whether strengthened central sensorimotor networks can support functional recovery when the peripheral substrate is compromised. Both possibilities remain hypotheses to be tested in patient cohorts; the present study establishes only that iNM can selectively reconFig. these networks with effects that persist beyond the active modulation window in healthy individuals.

### iNM enhances the consistency, specificity, and stability of neural activation across TMSC networks

While previous research has established that motor learning and neuromodulation can induce functionally specific changes in motor and somatosensory networks, these effects have typically been observed after training or in response to external cues. The findings of the current study demonstrate that iNM enhances specific and reliable neural encoding across TMSC networks indicated by high SVM classification accuracy and reduced BOLD signal magnitude variance, representing a significant advance over prior work. These results position iNM as a precision platform for enhancing functional specificity and stability in sensorimotor networks with translational relevance for neurorehabilitation^52^. This platform-intervention establishes a direct, one-to-one contingency between the performed physiological response (actual tongue movement) and the individually targeted, millimeter-precise network. This enables stable, adaptive control between afferent and efferent networks^52^. Consequently, iNM is both functionally and anatomically specific, minimizing the diffuse activation patterns and intersubject variability observed in neurofeedback approaches^57,58^ that rely on generic, atlas-based regions or whole-brain networks. With emerging consensus in the field that personalization of neural targets, precise temporal encoding-decoding of BOLD responses, and explicit performance-based cues, as opposed to mental imagery, are critical for optimizing iNM outcomes, this platform provides a technically rigorous and clinically scalable framework for next-generation, individualized neuromodulation interventions^52^.

Temporal analyses revealed that iNM effects persisted beyond the active modulation window (6–32 s), extending into the swallow (32–42 s) and baseline-tongue-at-rest (42–56 s) periods, as evidenced by sustained increases in BOLD magnitude and decreased variance of that magnitude, each quantified by AUC (Figs. 4, 6). Evaluating both mean and variance is essential: it distinguishes adaptive, directionally stable network reorganization from a nonspecific reduction in neural noise, and true SNR enhancement from transient fluctuation This dual-metric pattern indicates that iNM achieves targeted modulation with adaptive persistence across TMSC directions and networks, providing a framework for optimizing individualized protocols for clinical applications^1–5^.

### Bilateral inferior parietal lobule (IPL) engagement underlies adaptive, circuit-specific modulation during iNM

The bilateral IPL emerged as a central integration hub for iNM-induced sensorimotor adaptation. Under control, IPL responses oscillated around zero, indicating no consistent task-locked recruitment; under iNM the region became reliably engaged, showing high directional discriminability (d′) and the largest spatial expansion in the study (134.8% right, 81.8% left). Against a network-wide reduction in BOLD variance, IPL variance instead rose by 53% relative to control. Because this increase co-occurred with high d′ and the greatest spatial recruitment - rather than with degraded decoding accuracy - it is unlikely to reflect signal instability and is instead consistent with dynamic prediction-error processing. A region representing the trial-to-trial mismatch between predicted and actual sensory feedback must produce a variable signal, since an error that did not change across trials would carry no corrective information; the elevated variance therefore may encode the error term that drives real-time updating of the tongue motor plan. This interpretation aligns with prior work linking the IPL to forward-model correction, gamma-band synchronization, and precision-weighted error processing^59–61^. Temporally, the right IPL peaked before the left, consistent with right-lateralized sensory and attentional processing preceding left-lateralized motor control. Together, these features position the IPL as a multimodal integration center for the adaptive, lateralized modulation of TMSC.

### iNM engages a temporally coordinated network supporting anticipatory control, sensory integration, and motor refinement

Early peak activation in the right amygdala, followed by medial inferior frontal (MIF) and bilateral S1 activation, suggests that affective-salience regulation by the amygdala initiates anticipatory control and proprioceptive tuning, followed by motor execution^62^. Amygdala-MIF interactions reflect involvement in task salience and internal state monitoring, while early S1 activation (right: 8.6 s; left: 12 s) supports real-time proprioceptive and interoceptive modulation. Bilateral M1 engagement occurred later (~28 s), reinforcing a hierarchical sequence where sensory tuning precedes motor output. Concurrently, the sequential activation of cingulate cortices and inferior temporal lobes indicates layered cognitive control, imagery, and motor planning^63,64^. Finally, peak engagement of the cerebellum (left: 14 s; right: 14.7 s) and basal ganglia (left: 16 s; right: 16.4 s) underscores their role in adaptive, time-locked regulation of tongue movement^65^. The clustered peak activation of the cerebellum (14–14.7 s), basal ganglia (16 s), and right IPL (~16 s) identifies a critical integrative window where subcortical error-correction and parietal spatial updating converge. This mid-latency processing likely serves as a functional gate through which the updated internal model guides primary motor cortex execution at 28 s, near the close of the 30 s TMSC window, reflecting the convergence of sensorimotor integration onto motor output.

### Temporal causal modeling via unsupervised autoregressive hidden Markov modeling (ARHMM)

Unsupervised ARHMM revealed that iNM consistently stabilized the TMSC-dominant brain state, suppressing swallow-related interference, while it enhanced TMSC state persistence not only during tongue movement, but also during swallowing and rest intervals. During TMSC blocks, 97% of iNM trials were classified as TMSC compared to 82% in the control, with similar dominance observed during swallow and baseline periods, indicating sustained TMSC network engagement. This was accompanied by reduced BOLD signal variance, reflecting improved and consistent cortical responsivity and minimized noise, hallmarks of precise and stable engagement of physiologically relevant networks, time-locked to the instructed motor and sensory control output. These findings are indicative of experience-dependent plasticity, the biological mechanism that strengthens the TMSC network.

Furthermore, iNM significantly enhanced connectivity within and across motor, sensory, and attention networks (53–74% increase; Fig. 6, Extended Data Figs. 9-12), while simultaneously suppressing reward-network interactions. These findings reinforce iNM’s rehabilitative specificity and translational potential. Specifically, iNM promotes both functional specificity and adaptive persistence by optimizing TMSC stability without disrupting flexibility. iNM achieves robust coupling between cortical and physiological responses while minimizing stochastic noise, preserving the brain’s ability to transition between functional states. This stabilization of task-relevant states across time windows supports a mechanistic foundation for neurorehabilitation, where lasting gains depend on generalization beyond trained tasks^66–68^.

iNM produced the strongest neuromodulatory effects in motor and sensory regions, reflecting the high specificity of tongue representations in these areas and the brain’s innate ability to modulate these systems through adaptive control. Motor pathways are highly organized, densely myelinated, and evolutionarily tuned for rapid sensorimotor integration, especially responsive to adaptive control and error correction^69–71^. Adaptive control of motor and sensory networks enhances the continuous refinement of movement and the sensory system’s ability to estimate the state of the body, in this case, the tongue and its environment, the oral cavity and surrounding tissues. This precise targeting of functionally relevant networks supports strong and consistent neuromodulatory effects. The reward network illustrates a dissociation between local signal magnitude and network-level recruitment. Its AUC and d′ score gains reflect passive hemodynamic propagation: because the reward regions share draining vasculature and arousal-related global signal with the strongly driven motor, sensory, and attention nodes, any large increase in local metabolic demand raises their BOLD magnitude without implying task-specific engagement. Connectivity and spatial extent, by contrast, reflect whether a region is actively integrated into the TMSC network. On these measures the reward network moved in the opposite direction, losing connectivity in both the dominant (−99%) and non-dominant (−89%) states and contracting in spatial extent (−11.4%). The coexistence of rising magnitude with falling connectivity and shrinking spatial extent indicates that iNM did not recruit reward circuitry into the TMSC network; rather, it functionally pruned this network while sparing the incidental hemodynamic spillover. This dissociation confirms that iNM’s sensorimotor specificity operates at two levels: the precision of its guided targeting and the selectivity of its network-wide effects^52^. This dual-level specificity suggests that iNM enhances neural efficiency by engaging only physiologically relevant circuits, unlike the diffuse or compensatory activations observed in traditional neurofeedback paradigms^72^.

### Future Directions: additional iNM applications for motor and sensory control

Functionally, iNM is a real-time, adaptive brain-computer platform that modulates circuit-level dynamics at the individualized level, mirroring key features of a neurofunctional twin: continuous tracking, personalized targeting, and real-time adjustment of brain networks. As an adaptive closed-loop system, iNM reconFigs. TMSC circuits precisely and adaptively, with effects that persist beyond the neuromodulation window within the same session, addressing the patient-specific damage patterns characteristic of CCN or neurodegenerative disease. By accessing each subject’s full brain and adapting to individual connectivity patterns, iNM offers a pathway toward high-precision neuromodulation for functional recovery across diverse neurological conditions.

Although the present findings derive from healthy participants, the network-level mechanisms iNM engages are relevant to conditions in which TMSC circuitry is disrupted. Beyond its application in radiation-induced cranial neuropathy (RICN), iNM holds promise for other forms of cranial and peripheral neuropathies. CCN and tongue motor deficits can also occur after anterior cervical discectomy and fusion (ACDF), one of the most common cervical spine procedures. About 34–38% of patients experience dysphagia^73^, with 25.5% of those developing chronic symptoms that can progress to tongue atrophy^74–83^ rate^84,85^, it carries risks such as esophageal perforation, laryngeal nerve palsy, and dysphagia^74,75,86–93^. About 34–38% of patients experience dysphagia^73^, with 25.5% of those developing chronic symptoms that can lead to tongue atrophy^74–83^. As with CCN in head and neck cancer survivors, effective treatments remain limited^94–104^; iNM may therefore offer a precision neurorehabilitation strategy post-ACDF surgery.

Beyond CCN, iNM can be used for peripheral neuropathies experienced after breast cancer (BCA) surgery or radiation, applying the same principle of strengthening motor and sensory networks that control arm movement and sensation^105^. Despite high survival rates, 20–60% of BCA patients develop post-mastectomy pain syndrome (PMPS), a debilitating neuropathy following lymph node dissection^106–111^. As in cranial neuropathy, PMPS is fundamentally a sensory disorder, and the resulting sensory disruption impedes movement: the pain interferes with inspiration, expiration, and trunk movement, accompanied by severe depression; 54% of affected survivors report disruptions to employment and access to healthcare.^112–121^ As with CCN, effective treatments are lacking^122–125^. iNM offers a targeted, circuit-based approach that could be extended to sensorimotor pathways involved in arm strength, proprioception, and the sensory processing that constrains movement. Therefore, its potential as a precision intervention extends beyond orofacial applications^105^.

The functional persistence of iNM effects beyond the neuromodulation window has direct implications for clinical scalability. Because iNM precisely targets the individual’s network from the outset, it produces measurable within-session network reorganization that persists beyond the active modulation window, suggesting that fewer sessions may be required to achieve clinically meaningful outcomes compared to non-individualized approaches. This efficiency, derived from precision targeting rather than repeated, generalized or region-specific static feedback, positions iNM as a scalable intervention for populations where treatment burden is a critical barrier, such as head and neck cancer survivors and early-stage neurodegenerative disease patients.

Together, these applications demonstrate that iNM’s precision-guided architecture is translatable across a spectrum of neurological, post-surgical, and post-radiation conditions, with the potential to transform neurorehabilitation. By integrating adaptive control, real-time neuroimaging, and millimeter-precise targeting, iNM serves as a neurofunctional model (individualized and predictive) that updates the brain’s TMSC networks, rather than a fixed therapeutic tool. Specifically, its ability to access each subject’s entire brain is a key factor to successfully manage unique symptoms to each patient, based on the afferent or efferent nerves lesioned, and thus, brain networks involved. In doing so, iNM not only advances the field of brain-computer interfacing but also lays the foundation for a new generation of non-invasive, individualized neuromodulation therapies.

## Acknowledgements

This study is funded by grants to T.D. Papageorgiou: McNair Medical Institute; Robert and Janice McNair Foundation; Mission Connect, IP Award - The Institute of Rehabilitation Research; Center for Alzheimer’s and Neurodegenerative Disease; Jan and Dan Duncan Neurological Research Institute; Naman Basic Science Faculty Award; Fight for Sight; NIH-T32, NEI; NIH-T32, NINDS, Breast SPORE – Development Research Project Award.

TDP wishes to thank Dr. Sameer A. Sheth, Professor – Cullen Foundation Endowed Chair of Neurosurgery and Vice Chair of Research Neurosurgery at Baylor College of Medicine for his support of this study.

TDP gratefully acknowledges the legal guidance provided by the intellectual property team Kevin Keeling and James P. Cargas, whose expertise led to the patent of the iNM platform (Patent No. 19 753 851.5, WIPO Publication No. WO 2019/160754).

## Ethics, Consent to Participate, and Consent to Publish declarations

Informed consent was obtained from all human research participants

## Author Contributions

A. Conceptualization, development of iNM: T.D.P.

Research Design and Methodology: T.D.P.

Software & Analytical Tools: T.D.P., J.W., A.A., S.R., C.N., D.H., R.H., E.F.

Validation & Formal Analysis: T.D.P., J.W., A.A., S.R., C.N., D.H., R.H.

Investigation & Experiments: T.D.P., D.H.

Funding Acquisition: T.D.P.

Supervision & Project Administration: T.D.P.

Writing – Original Draft: T.D.P.

Writing – Review & Editing: T.D.P., E.F., J.W., A.A., K.H., E.M.R., E.M.S., A.M.T., S.G.H., R.H., C.N.

