## Supplementary Figures 1-18 and Supplementary Tables 1-4 for "Individualized AI-driven Neuromodulation selectively enhances and reorganizes human tongue sensorimotor networks"

**Acknowledgements:** This study is funded by grants to T.D. Papageorgiou: McNair Medical Institute; Robert and Janice McNair Foundation; Mission Connect, IP Award - The Institute of Rehabilitation Research; Center for Alzheimer’s and Neurodegenerative Disease; Jan and Dan Duncan Neurological Research Institute; Naman Basic Science Faculty Award; Fight for Sight; NIH-T32, NEI; NIH-T32, NINDS, Breast SPORE – Development Research Project Award.

TDP wishes to thank Dr. Sameer A. Sheth, Professor – Cullen Foundation Endowed Chair of Neurosurgery and Vice Chair of Research Neurosurgery at Baylor College of Medicine for his support of this study.

TDP gratefully acknowledges the legal guidance provided by the intellectual property team Kevin Keeling and James P. Cargas, whose expertise led to the patent of the iNM platform (Patent No. 19 753 851.5, WIPO Publication No. WO 2019/160754).

**Ethics, Consent to Participate, and Consent to Publish declarations**: Informed consent was obtained from all human research participants

**Competing Interest Declaration --** Patent granted to Dr. T. Dorina Papageorgiou & Dr. E. Froudarakis: No. 19 753 851.5, WIPO Publication No. WO 2019/160754

**Author Contributions:**

- Conceptualization, development of iNM: T.D.P.
- Research Design and Methodology: T.D.P.
- Software & Analytical Tools: T.D.P., J.W., A.A., S.R., C.N., D.H., R.H., E.F.
- Validation & Formal Analysis: T.D.P., J.W., A.A., S.R., C.N., D.H., R.H.
- Investigation & Experiments: T.D.P., D.H.
- Funding Acquisition: T.D.P.
- Supervision & Project Administration: T.D.P.
- Writing – Original Draft: T.D.P.
- Writing – Review & Editing: T.D.P., E.F., J.W., A.A., S.R., C.N., H.K.A., E.M.R., E.M.S., A.M.T., S.G.H., S.R.H., C.R.N.

**Materials and Methods**

**Subjects**

Thirty healthy, right-handed volunteers (19 male, 11 female, mean age = 25.37±5.56) were recruited in this two-day study Written informed consent was obtained from all participants in accordance with the Declaration of Helsinki, under a protocol (H-33628) approved by the Institutional Review Board of Baylor College of Medicine. Exclusion criteria included any current or prior medical or psychiatric diagnosis, use of prescription medications (over-the-counter use was permitted), and standard contraindications to MRI. Participants had no history of head, neck, or brain radiation or surgery, no neurological deficits, and normal or corrected-to-normal vision (with MR-compatible glasses where needed). Participants were compensated at the end of each session.

**MRI and fMRI Pulse Sequence Parameters**

Structural and functional imaging was performed at the Core for Advanced Magnetic Resonance Imaging (CAMRI), Baylor College of Medicine, Houston, Texas, on a 3.0 T Siemens MAGNETOM TrioTim (syngo MR B19; Siemens, Erlangen, Germany) using a 64-channel head/neck receiver-array coil. A T1-weighted 3D magnetization-prepared rapid gradient-echo (MPRAGE) sequence acquired 192 transversal slices at 1.0 mm isotropic resolution (FOV = 245 × 245 mm²; base resolution = 256; TR = 1,200 ms; TE = 2.66 ms; TI = 600 ms; flip angle = 12°; slice resolution = 80%; bandwidth = 210 Hz/pixel), reconstructed with in-plane interpolation to a 0.479 × 0.479 × 1.0 mm voxel grid (448 × 512 × 192). Functional data comprised 32 contiguous interleaved slices (0% distance factor) acquired with a T2*-weighted gradient-echo echo-planar imaging (EPI) sequence (ep2d_bold; FOV = 200 × 200 mm²; matrix = 64 × 64; voxel size = 3.125 × 3.125 × 3.0 mm; TR = 2,000 ms; TE = 30 ms; flip angle = 90°; bandwidth = 2,442 Hz/pixel; echo spacing = 0.47 ms; EPI factor = 64; phase-encoding direction A→P; fat saturation; no parallel-imaging acceleration), with slices prescribed at a 16.7° oblique angle and 244 volumes per run.

**Real-time fMRI Neuromodulation Acquisition**

Turbo-BrainVoyager (TBV; version 2.0; Brain Innovation, Maastricht, The Netherlands)^112^ software performed the following preprocessing computations on the EPI images acquired at each repetition time (TR): (i) 3D motion correction; (ii) incremental linear detrending to remove BOLD signal drifts; (iii) computation of statistical brain maps generated from a general linear model (GLM) along with beta weights (BOLD signal intensity values) for each condition; (iv) extraction of the average BOLD signal intensity values from the individualized networks defined on Day 1 scans (see *Data Analysis*); and (v) presentation of the network’s average BOLD signal intensity via the neuromodulation interface (Fig.1D). The steps of the iNM intervention are summarized in Fig. 1. To further increase the signal-to-noise ratio (SNR), we applied an exponential moving average (EMA) algorithm to high-pass filter the ROI’s average BOLD signal and suppress low-frequency noise components, such as scanner drifts, and physiological noise from heart rate and respiration. The EMA output of was then low pass filtered with a Kalman filter to eliminate high frequency noise.^113^ Details on the EMA and Kalman filter computations can be found in Allam and colleagues (2024).^47^

**Study Structure**

Our study comprised two day-sessions, separated by 5–7 days. Day 1 consisted of four functional (EPI) scans: three functional task scans to delineate the iNM network and one iNM scan. Day 2 consisted of six EPI scans: three control-no iNM scans, which alternated with three iNM scans. Each EPI scan comprised eight continuous periods and lasted 8 minutes and 8 seconds (244 volumes × 2 s), inclusive of 4 dummy TRs (3 at the start to allow task initiation and steady-state magnetization, and 1 at the end). Within each period, subjects were cued to move their tongue in one of four directions (up, down, left, right). Each tongue-movement block lasted 30 s and was interleaved with a swallow block (6 s) and a baseline-tongue-at-rest block (20 s). Analysis windows were shifted relative to cue onset to account for the hemodynamic lag of the BOLD response, so the TMSC window began at 6 s rather than at cue onset. This ensured the measured signal reflected task-evoked activity rather than residual response from the preceding block, preventing baseline contamination and improving TMSC time-series specificity. The directions of the movement blocks were randomly counterbalanced across scans and followed three rules: (i) each TMSC direction block occurred twice during each period; (ii) a TMSC direction block was never followed by the same TMSC direction cue; and (iii) each scan consisted of a unique order of blocks (Fig. 1B).

All subjects were instructed by the principal investigator (TDP) to foster consistency in the receipt of instructions and training. These standardized instructions minimized confounding factors unrelated to tongue movement and promoted consistent TMSC, enabling reliable replication across subsequent scans. Subjects were coached to move their tongue without engaging other facial or jaw movements prior to participating in the study. After brief coaching, all 30 subjects were able to perform isolated tongue movements and were included in the study. Participants were asked to control their tongue movement in a consistent manner with respect to two parameters: (i) the force of tongue movement and (ii) the location of tongue placement in the mouth. During the 30-s tongue movement period, subjects were also coached to abstain from clenching their teeth, pursing their lips, squinting, applying pressure to their cheeks, and straining their facial muscles, such as those of the forehead between the eyebrows and at the corners of the eyes; they were also instructed to keep the lower jaw still, avoiding any backward, forward, or side-to-side movement. Subjects were instructed to move and place their tongue in the same location in the mouth by mentally assigning landmarks, such as a specific ridge on the roof of the palate, or a specific tooth for left and right movements. A time interval was dedicated to swallowing after each TMSC block. Swallowing could be detected in real time during the TMSC task, because the fMRI signal was significantly upregulated during swallowing compared with TMSC. Subjects adhered to abstinence from swallowing during TMSC task execution; therefore, swallowing did not have to be excluded from the TMSC-dedicated time series signal. During the rest block, which served as the baseline, participants stabilized their tongue between the hard and soft palate without touching either surface, because contact would be registered as tongue movement and would increase the hemodynamic response, thereby artifactually elevating the baseline time series.

**Study Design**

**Overview:** On Day 1, we acquired three control-no-iNM scans to delineate each participant’s individualized network. These networks captured each participant’s unique TMSC BOLD amplitude and spatial extent during tongue movement as determined by general linear model (GLM). During the control scans, the visual cue (tongue of the emoji; Figs. 1A, 1C) was color-coded in red to motivate subjects to execute the task at maximal performance. Each individualized network served as the targets during iNM. which was applied in real-time every two seconds during the tongue movement blocks.

**Computing Individualized Network for Neuromodulation Delivery:** We acquired functional localizer scans on Day 1 to identify each participant’s individualized TMSC network. After scanning, the anatomical and functional images were processed offline using AFNI.^114^ The anatomical data was spatially transformed to Talairach space using an average volume of 452 skull-stripped brains (TT_icbm452). The functional data were preprocessed to reduce artifacts and increase SNR. The signal pre-processing protocol included: 1. removal of outliers (head motion, physiological artifacts; 3dDespike) from the time series; 2. slice-time correction (3dTShift); 3. transformation of the oblique-acquired functional dataset to a cardinal dataset (3dWarp -oblique2card); 4. motion correction by registering each functional dataset to the first volume of the first functional scan, using a 3D-rigid-body transformation; 5. spatial smoothing with a 6 mm-FWHM Gaussian kernel filter; and 6. co-registration of the functional data with the individual T1-weighted 3D-structural data.

The three control-no-iNM scans acquired on Day 1 were then concatenated to increase the SNR prior to generating parametric brain maps across the four TMSC directions via a General Linear Model (GLM; 3dREMLfit, which adjusts for temporal correlations in each voxel’s time series, AFNI). The GLM included: 1. four regressors, one for each of the four TMSC directions (left; right; up; down cortical direction selectivity); 2. six motion covariate vectors that controlled for head motion; 3. white matter and cerebrospinal fluid means were regressed out to increase our SNR, since activity in these areas represents noise; and 4. the baseline-tongue at rest regressor (tongue held between the hard and soft palate). Each directional regressor was convolved with a gamma-variate canonical hemodynamic response function. The swallow condition was excluded from the GLM, since the number and onset of swallows were not controlled. The first 3 TRs (6 s) were removed from the baseline-tongue-at-rest condition to account for hemodynamic-response lag. Following generation of a TMSC-versus-baseline network (voxelwise p = 0.001), a cluster analysis identified highly significant regions (FWER < 0.05; Fig. 2).  This individualized network, delineated from the Day 1 localizer (Fig. 2), constituted each participant's neurofunctional twin (functional and anatomical network), which was superimposed on the participant's brain and updated in real time during iNM.

**Neuromodulation Paradigm:** These individualized networks -- each participant’s unique BOLD amplitude and spatial extent – served as the iNM targets with the goal of achieving maximum upregulation during tongue movement (Fig. 1A). The color and extent of the emoji’s tongue, which directly represented the magnitude and spatial extent of each subject’s individualized network, conveyed the degree of network upregulation or downregulation in response to tongue movement (Fig. 1C). The iNM signal was calculated by first acquiring the percent BOLD signal change (% BOLD PSC) generated during each control scan for each TMSC direction relative to the rest block that preceded it. We computed the % BOLD PSC change from each participant’s individualized areas every 2 seconds as follows:

*BOLD PSCi(j) = 100% * [ROIs BOLD during each TMSC direction_i(j) – ROIs BOLD during tongue-at-rest_i(j)]/tongue-at-rest_i(j)*

where *i* represents each TMSC direction (left, right, up, down), *j* represents the time interval (2secs) over which the BOLD PSC of tongue motor control direction *i* was computed. The iNM value presented at each TR (2 seconds) was computed by comparing the current PSC value against a graded scale of 25% BOLD increments: −100%, −75%, −50%, −25%, 0, 25%, 50%, 75%, 100% (Fig. 1D). To increase task demand during iNM relative to control, this scale was shifted upward by 10%; consequently, a BOLD level that filled the tongue red during control produced a lesser red fill during iNM, requiring participants to generate stronger network upregulation to reach the same fill. During the iNM scan that followed each control scan, if the PSC at a given time point was positive (25% to 100%), the tongue was filled red in proportion to the increment reached, representing upregulation of the BOLD signal of the targeted network that controlled a specific tongue movement (Fig. 1C). At 0%, the tongue was filled pink, representing baseline. If the PSC was negative (−25% to −100%), the tongue was filled white in proportion to the increment reached, representing downregulation of the BOLD signal of the targeted network that controlled a specific tongue movement (Fig. 1C).

**Support Vector Machine Classification for Direction of Controlled and Voluntary Tongue Movement:** Linear support vector machines (SVM) were used to decode the BOLD spatial patterns of each TMSC direction generated by the 3 iNM and 3 control scans. The 3dSVM AFNI function implements a linear SVM classification in which the weight vector specifies the signal direction for classification in the multivariate image space.^115–117^

We applied a cross-validation approach for each cortical selectivity classification: 1. concatenated all TMSC directions; 2. left direction; 3. right direction; 4. up direction; 5. down direction vs. baseline-tongue at rest. For each subject, we computed classification accuracies (CAs) for the following SVM training-testing permutations using the means from a 3-fold cross-validation method: i. training and testing with control data and ii. training and testing on iNM data (Fig. SI-1). We used soft margin SVMs to account for data points that were not linearly separable because we found no difference in performance over a wide range of the regularization parameter c (10⁻⁴ to 10⁺⁴, geometric progression with ratio 10), we applied the default c = 1 for all classifications to optimize training and testing error.

SVM brain maps were generated for each TMSC direction and condition (control or iNM) as follows: a. the weight vectors corresponding to the optimal hyperplanes obtained from training each of the 3 cross-validation datasets were mapped and mean-zero standardized; b. each subject's weight vectors were normalized; c. the mean weight vectors across the 30 subjects were entered into a t-test for group and cluster analysis (3dttest++ -ClustSim, AFNI); and d. the t-test-generated brain maps were z-scored (p < 0.001, cluster size > 10 voxels). From the activation map of each TMSC direction and condition, we identified each cluster's local maxima (3dExtrema, AFNI). When more than one local maximum occurred within a cluster, or a cluster spanned an anatomical region, only the voxels with the highest z-scores — corresponding to increased BOLD signal intensity — were included for that region.

**Percent Signal Change (PSC) Area Under the Curve:** We computed the BOLD percent signal change (PSC) for each TMSC direction and condition as a function of time. ROIs were categorized into motor, sensory, attention, and reward networks. To generate the BOLD PSC time series, we first averaged the voxel-wise time series within each ROI, then averaged across segments corresponding to blocks of the same tongue-motor-control direction from scans of the same type. Each segment began at the onset of a TMSC block and ended at the final time point of the swallow block. For each subject — 3 control and 3 iNM scans on Day 2, each scan containing 2 TM blocks per direction — the PSC time series for each direction, condition, and subject was the average over 6 BOLD signal segments. To compute the AUC, we first averaged the PSC across blocks for each time point, subject, and ROI (yielding a time × subject × ROI matrix). We then averaged these values across subjects to yield a time × ROI matrix, and then across ROIs to produce a single time series. Finally, Simpson's method was applied to this time series to estimate the average AUC for the network. The mean area under the PSC curve for each condition (control or iNM) was calculated using Simpson's Rule:

$${AUC}_{con}^{reg}={\frac{1}{3}({PSC}_{1}+4PSC}_{2}+2{PSC}_{3}+\ldots+4{PSC}_{T-1}+{PSC}_{T}$$

Where PSCt is the PSC at time t. The BOLD percent signal change was then calculated using the following formula:

$$100*({AUC}_{iNM}^{reg}-{AUC}_{ctrl}^{reg})/{AUC}_{iNM}^{reg}$$

**D’ Sensitivity Index:** To measure the strength of iNM over time and rank regions with respect to this strength, we computed the D’ sensitivity index, also known as Cohen’s d. This index quantifies the separation between the means of the control and iNM distributions, relative to their standard deviations. Assuming that the fMRI data set has $T$ time points (29 secs), $B$ blocks (6 blocks), $S$ subjects (n=30), and $N$ ROIs, the number of which ranges between 32-36 depending on direction (left direction 32 ROIs; right 36 ROIs; up direction 35; down direction 32). Each ROI is characterized by a three-dimensional matrix of dimensions $T\left( 29 \right)\times B\left( 6 \right)\times S$(30). Each subject is then characterized by a two-dimensional matrix of dimensions $T (29)\times B$ (6). For a fixed time point $t$, let the distribution of blocks for control and iNM be denoted as ${dist}_{ctrl}^{s}\left( t \right)$ and ${dist}_{nFb}^{s}(t)$, respectively. D’ for subject s at time point $t$ is:

$${d^{s}}^{'}\left( t \right)=\frac{\mu_{iNM}^{s}(t)-\mu_{ctrl}^{s}(t)}{\sqrt{\frac{1}{2}\left( {\sigma_{iNM}^{s}(t)}^{2}+{\sigma_{ctrl}^{s}(t)}^{2} \right)}}, t=1,\ldots,T$$

Where $\mu_{con}^{s}(t)=mean({dist}_{con}^{s}(t))$, $\sigma_{Con}^{s}(t)=std({dist}_{con}^{s}(t))$.

The sensitivity index between the two conditions at time point $t$ and for fixed region is the average sensitivity index across subjects:

$${{d^{'}\left( t \right)=\underset{s}{\mathrm{mean}} {{(d}^{s}}^{'}) , s=1,\ldots,S}}$$

For each subject, bootstrapped distributions of *d′* time series across all ROIs were divided into bins at the 5th, 15th, 25th, ..., 95th percentile thresholds. At each time point, the *d′* value a given ROI was assigned a discretized score from 1 to 10, reflecting the percentile bin in which it fell. The average d′ score for each network was then, determined as the mean of the d′ scores across all ROIs within that network.

Additionally, peak times were estimated by first applying cubic interpolation to each region’s *d′* score to generate a smoothed time series. The earliest peak was then identified using MATLAB’s “findpeaks” function with default parameters. These peak times are visualized in the heatmap representation (Figs. 5B, 5C).

**Network-Specific Mapping of Spatial Activation Extent:** ROIs were first identified using a general linear model (GLM) thresholded at p = 0.001*,* and FDR-corrected (q ≤ 0.01*.* Surviving clusters were then assigned to one of four non-overlapping functional networks: motor, sensory, attention, and reward. This classification was guided by anatomical criteria from the Talairach brain atlas and TMSC-specific functional activation patterns. For each subject and each region, we computed the number of activated voxels to quantify the spatial extent of activation, as shown in Figs. 5A; 5D; 5E. We then calculated the proportion of suprathreshold voxels within each anatomically defined ROI relative to that region’s total template-space volume, yielding a normalized activity index that accounts for differences in region size. These proportions were converted to z-scores across subjects, which provided standardized activation values for cross-subject and network-level comparisons. This approach enabled quantification and comparison of functional network engagement while controlling for anatomical constraints and inter-subject variability.

**Autoregressive Hidden Markov Model (ARHMM):** To elucidate the dynamic latent network states underlying TMSC during tongue movement—and to validate findings from BOLD signal quantification using the sensitivity index (d′), and AUC—we applied an unsupervised ARHMM. A step-by-step overview of ARHMM analysis is provided in Fig. SI-2. Briefly, the BOLD data were organized by stacking trials into a single dimension to enable ARHMM processing. The ARHMM was trained in an unsupervised manner on BOLD time series from TMSC-selective regions during tongue movement (6-32 seconds), learning latent state dynamics and transition probabilities directly from the data. Inference via the Viterbi trace was applied to the entire time series (0-56 seconds), characterizing brain states during TMSC, swallow, and baseline intervals for each condition (Figs. 6A, SI- 9--12).

Initially, a single ARHMM was fit to both conditions combined, after which the frequency of latent states was calculated separately for iNM and control trials. For each of the four TMSC directions under iNM and control conditions, we concatenated the six TMSC blocks across subjects to generate a dataset of dimensions *N × M × T* (ROIs × Trials × Time points), where *M = B × S* reflects the total number of blocks (*B = 6*) across subjects (*S = 30*). After collapsing across all four directions, this yielded a dataset of size *N × 4M × T*, comprised of 720 trials per condition. We performed a leave-one-subject-out analysis, applying the ARHMM to data from 29 subjects (696 trials) while holding out 24 trials from one subject in each iteration, yielding 30 models per condition. The resulting autoregressive (AR) coefficients generated average ROI-to-ROI connectivity matrices (Figs. SI-13—16), grouped by functional networks (motor, sensory, attention, reward). Network-level connectivity matrices were computed, and within each network, the summed differences in AR coefficients between control and iNM conditions quantified iNM-induced connectivity changes. Connectivity measures within and between the Motor, Sensory, Attention, and Reward networks were calculated as the AR coefficients between regions comprising each network.

**Extended Data Figures**

**Extended Data Fig. 1. Training and testing permutations for SVM classification using 3-fold cross-validation.**
SVM classification accuracy was assessed separately for the control (no-iNM) and iNM conditions using a 3-fold cross-validation approach in each of the 30 subjects. For each condition, the dataset comprised three independent fMRI scans. Classification accuracy was computed across the following permutations:

- Control Condition: Set 1: Train on Scans 1 & 2, Test on Scan 3; Set 2: Train on Scans 2 & 3, Test on Scan 1; Set 3: Train on Scans 1 & 3, Test on Scan 2
- iNM Condition: Set 1: Train on Scans 1 & 2, Test on Scan 3; Set 2: Train on Scans 2 & 3, Test on Scan 1; Set 3: Train on Scans 1 & 3, Test on Scan 2

For each subject, classification accuracy for each condition was obtained by averaging the results across all three test sets, for each TMSC direction (left, right, up, down) and across all TMSC directions. These per-subject accuracies were then averaged across the 30 subjects to yield the group-level classification accuracy reported in Fig. 3.

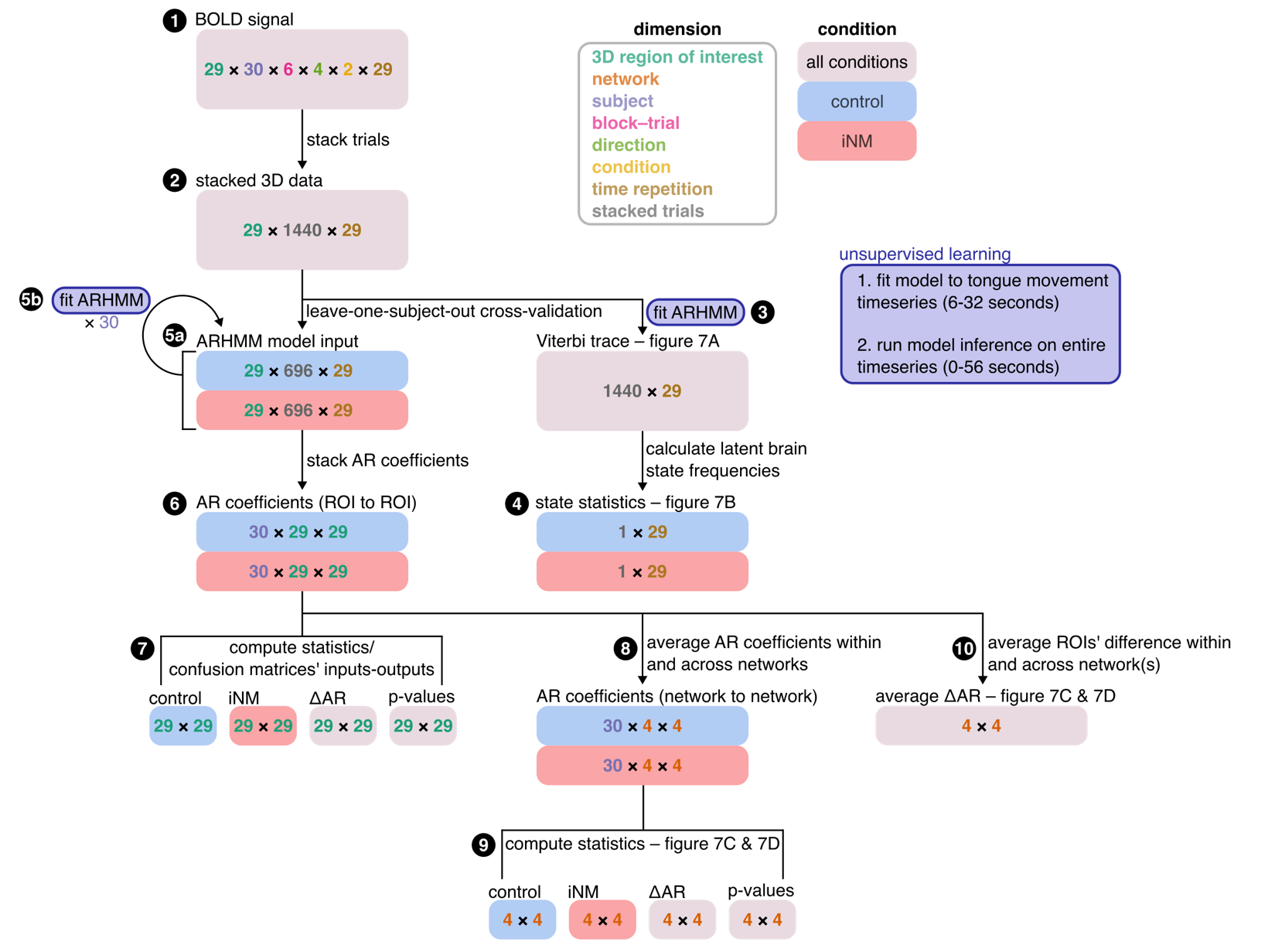

**Extended Data Fig. 2. Step-by-step overview of autoregressive hidden Markov model (ARHMM) analysis.**

1. At each TR (2 s), BOLD signal data were stacked by aggregating values from all subjects, regions, blocks, movement directions, and conditions corresponding to that time point, forming a temporally resolved matrix used as input to the AHRMM.
2. ARHMM was fit exclusively to time series corresponding to the tongue movement intervals (6-32 s).
3. Viterbi decoding was then used to infer latent state sequences across the entire time series (0-56 s) for both conditions (see Steps 3 & 5, inset in purple box).
4. Latent state frequencies were computed separately for trials from the control and iNM conditions (Step 4).
5. Next, leave-one-subject-out cross-validation was performed on the stacked dataset. All trials from one subject were held out (step **5a**), while ARHMM models were fit to data from the remaining twenty-nine subjects under each condition (step **5b**), resulting in a total of thirty ARHMM models for both the control and iNM conditions.
6. From the autoregressive (AR) coefficients for these models (step **6**), the average confusion matrices representing functional connectivity between regions-of-interest (ROIs) were calculated (step **7**).
7. AR coefficients were subsequently grouped by network (step **8**; i.e. motor, sensory, attention, reward), and confusion matrices representing network-to-network interactions were similarly computed (step **9**).
8. For each network, the cumulative difference in model coefficients between control and iNM conditions was calculated (step **10**).

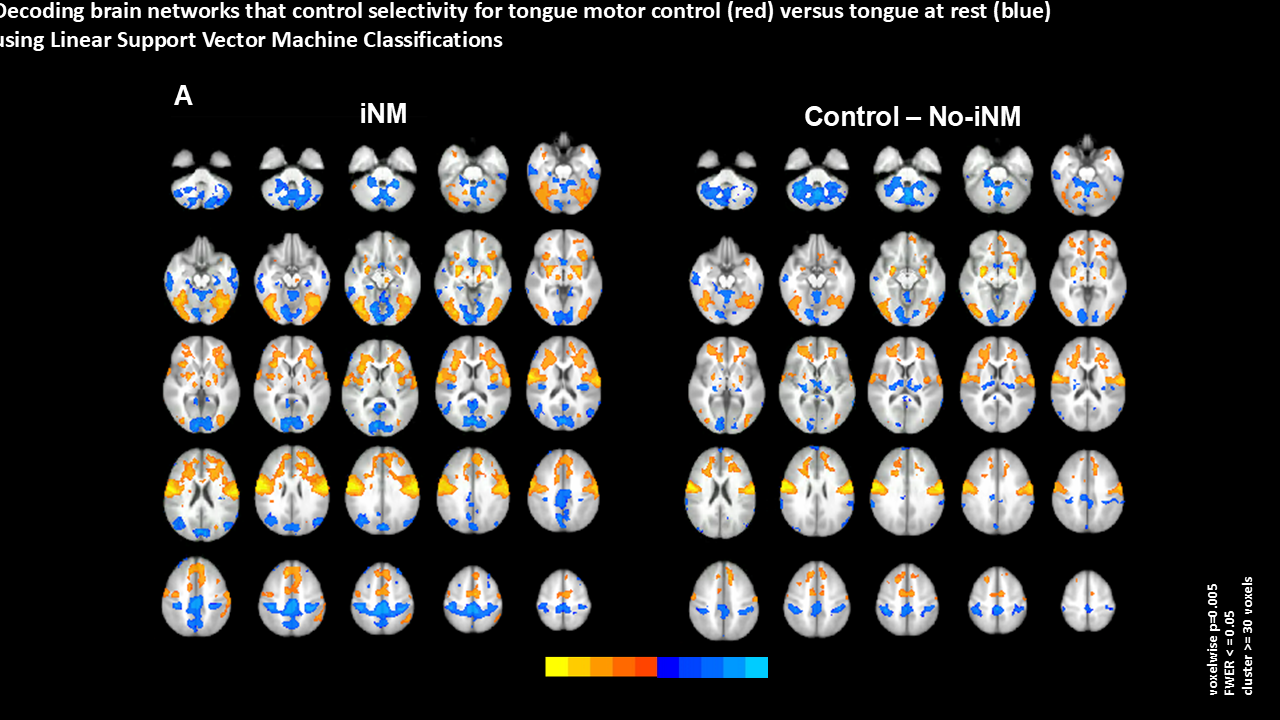

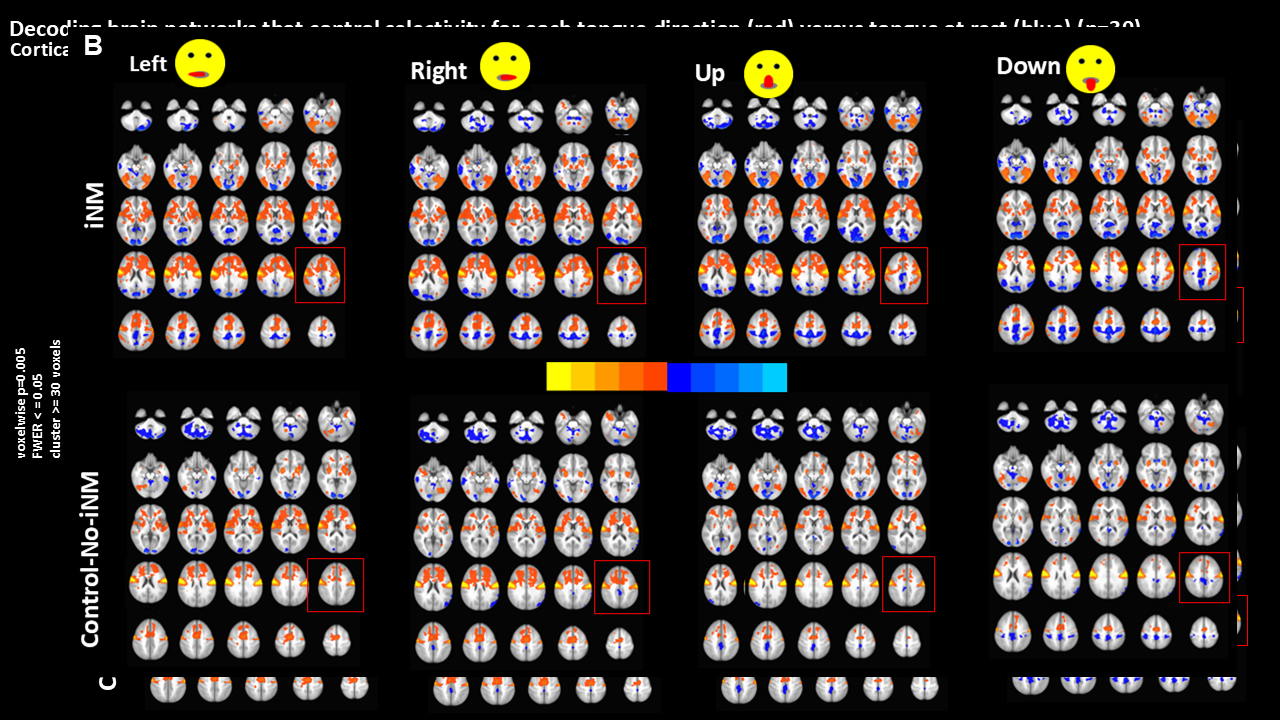

**Extended Data Fig. 3. Cortical maps show that iNM increases consistency in the BOLD signal intensity and spatial extent compared to the control-no iNM condition determined by linear support vector machine analysis.**

**A.** Concatenated activation maps across all TMSC directions for iNM and control conditions, thresholded at voxelwise p < 0.0001 (FDR-corrected, q ≤ 0.01) with a minimum cluster size >10 voxels.

**B.** Direction-specific SVM-generated activation maps under iNM (top) and control (bottom) conditions, using identical thresholding criteria.

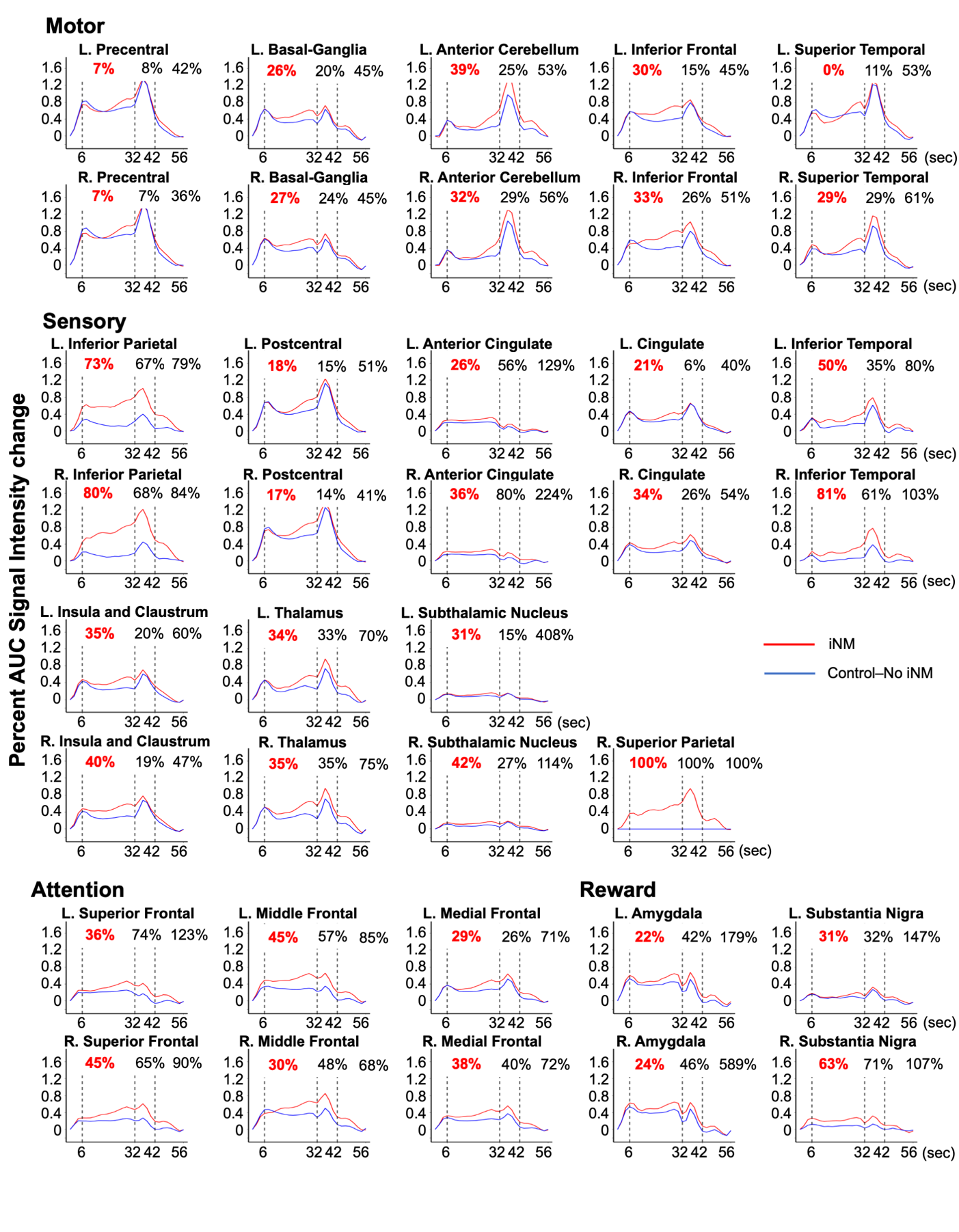

**Extended Data Fig. 4a. iNM (red) increases mean percent AUC signal intensity change compared to control-no iNM (blue) condition for each ROI within each network**. Mean percent AUC values are shown for each ROI categorized by network, across intervals: tongue movement (6–32 s), swallow (32–42 s), and baseline-tongue-at-rest (42–56 s). iNM led to enhanced AUC responses ranging from 7–39% in motor, 17–100% in sensory, 29–45% in attention, and 22–63% in reward-related regions, reflecting consistent signal amplification across networks. Notably, the iNM effects persisted into the swallow and rest blocks - periods without active modulation - indicating that iNM promotes functional specificity with adaptive carryover beyond the iNM window.

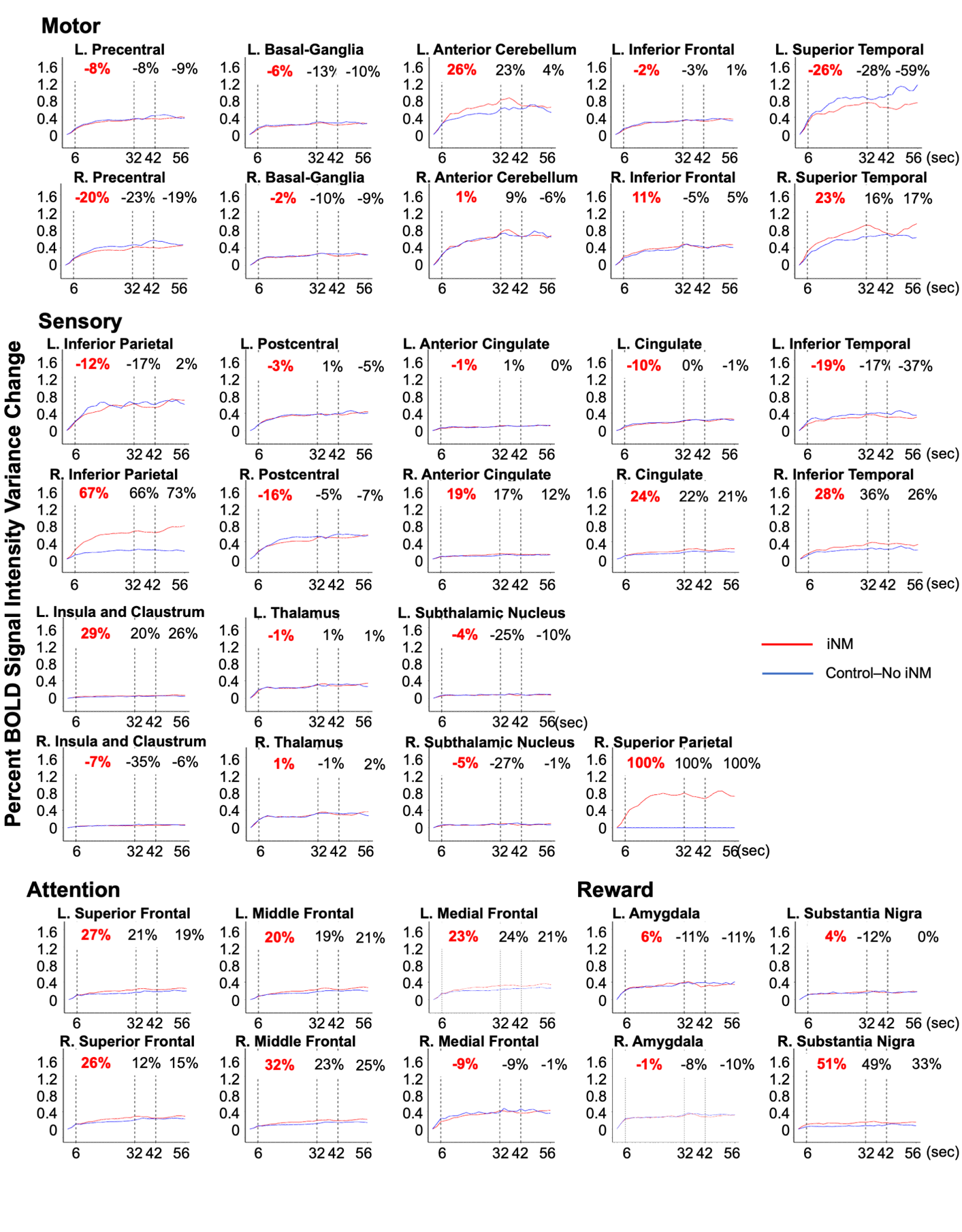

**Extended Data Fig. 4b. iNM (red) reduced the variance of mean percent AUC signal intensity change relative to the control (blue) across all ROIs, categorized by network.** AUC values are shown for tongue movement (6–32 s), swallow (32–42 s), and baseline-tongue-at-rest (42–56 s). Variance decreased consistently across motor, sensory, attention, and reward regions. Notably, these effects persisted into the swallow and rest blocks—intervals without active modulation—indicating that iNM enhances functional specificity with adaptive carryover beyond the iNM window.

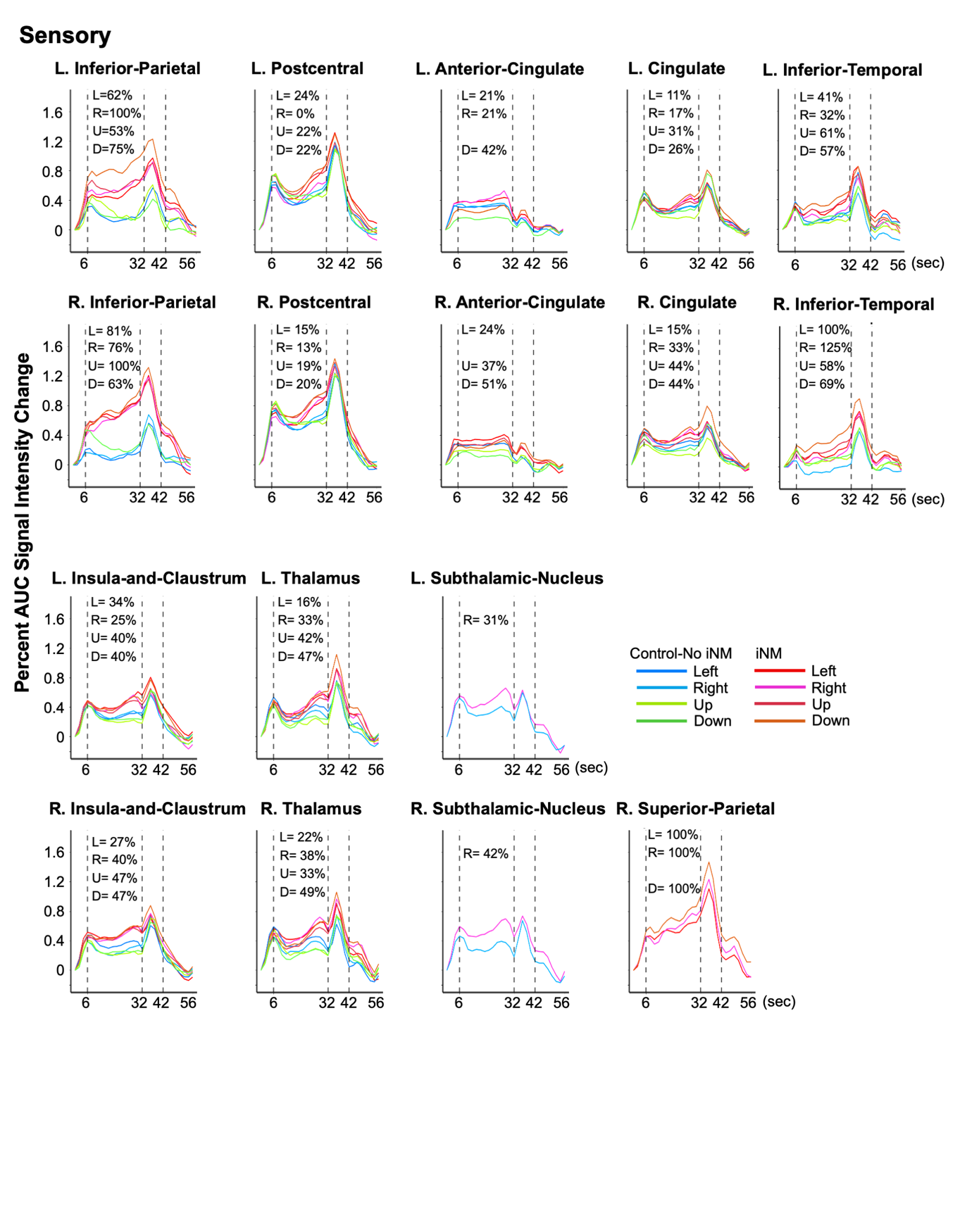

**Extended Data Fig. 5a. iNM enhances mean percent area under the curve (AUC) of the BOLD signal intensity during tongue motor and sensory control.** iNM increased the mean percent AUC during the TMSC interval (6–32 seconds) compared to the control-no-iNM condition, across cortical direction-selective regions of interest (ROIs) within the sensory network.

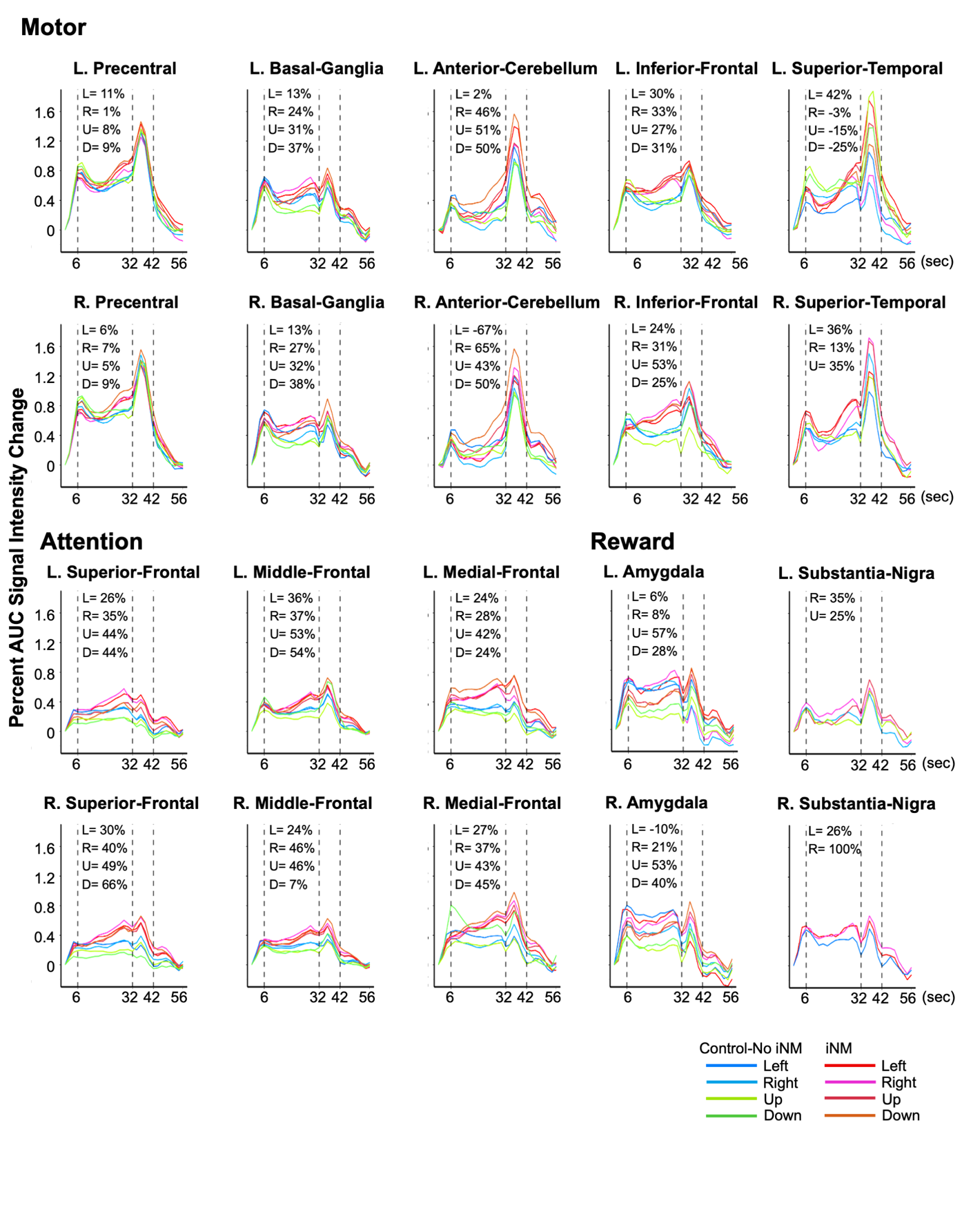

**Extended Data Fig. 5b. iNM enhances mean percent area under the curve (AUC) of the BOLD signal intensity during tongue motor and sensory control.** iNM increased the mean percent AUC during the TMSC interval (6–32 seconds) compared to the control-no-iNM condition, across cortical direction-selective regions of interest (ROIs) within the motor, attention, and reward networks.

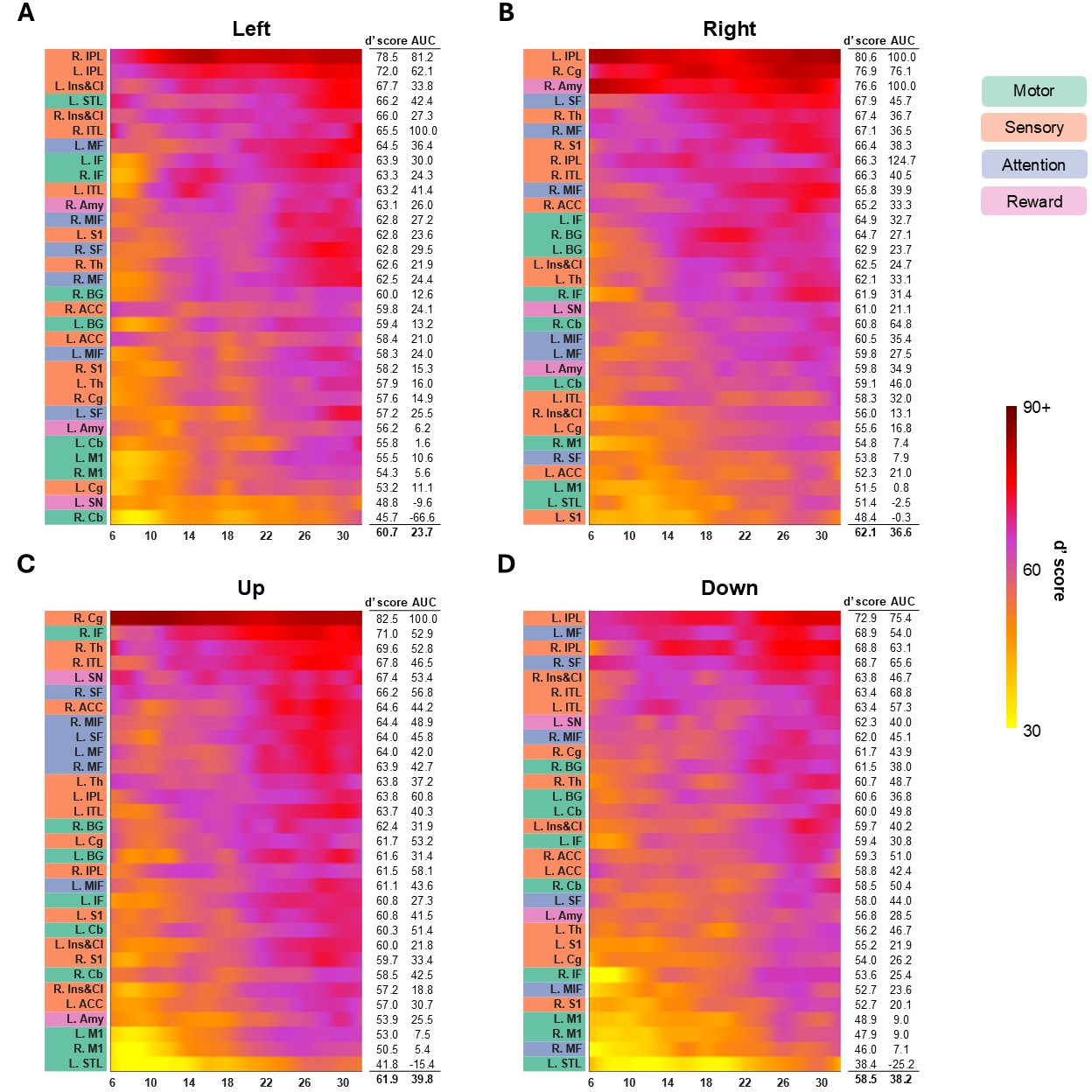

**Extended Data Fig. 6. iNM increases the sensitivity index (d’), d’ score, and AUC spatiotemporally for TMSC direction selectivity as a function of direction compared to control.** As shown in Fig. 5, the heatmaps for left (A), right (B), up (C), and down (D) tongue directions display the mean d′ score for each region, comparing the BOLD response between iNM and control conditions across time (TR = 2 s). Each heatmap presents the mean d′ score, peak d′ value, and AUC during the tongue movement interval (6–32 s), with region labels color-coded by network affiliation. The bilateral inferior parietal lobule (IPL) exhibited the highest d′ scores for left (R. IPL: 78.4; L. IPL: 71.8), right (L. IPL: 80.5; R. IPL: 70.8), and down (L. IPL: 72.9; R. IPL: 68.8) TMSC direction selectivities. For the up direction, the right IPL exhibited the highest d′ score (83.7), while the left IPL showed a moderate d′ score (62.2) relative to other regions engaged during upward movement.

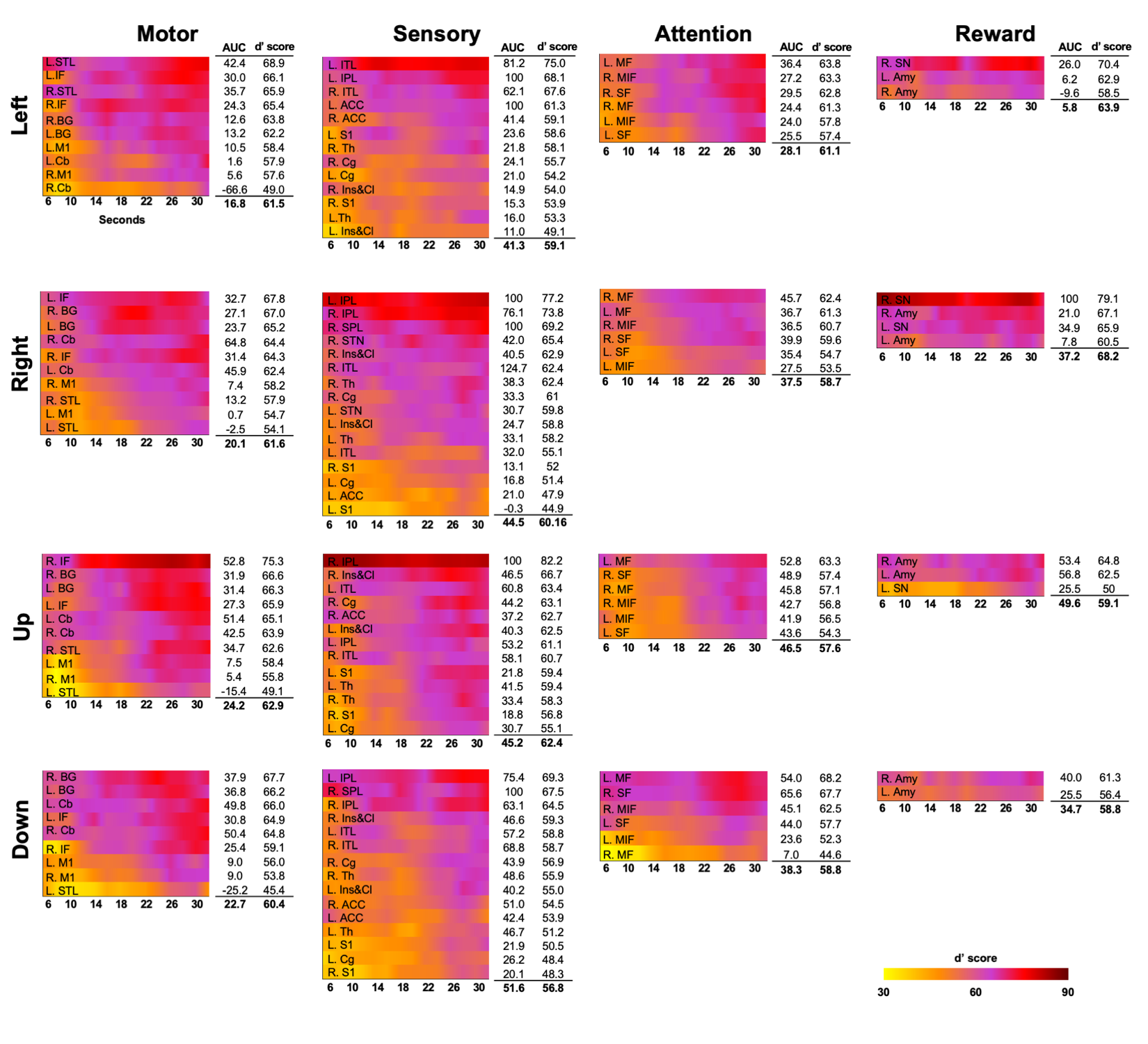

**Extended Data Fig. 7. iNM enhances the sensitivity index (d′) and AUC for TMSC direction selectivity, shown as spatiotemporal heatmaps organized by functional network and movement direction.** Heatmaps display d′ scores over time for each region, alongside mean AUC values, highlighting iNM-driven improvements in directional specificity across networks.

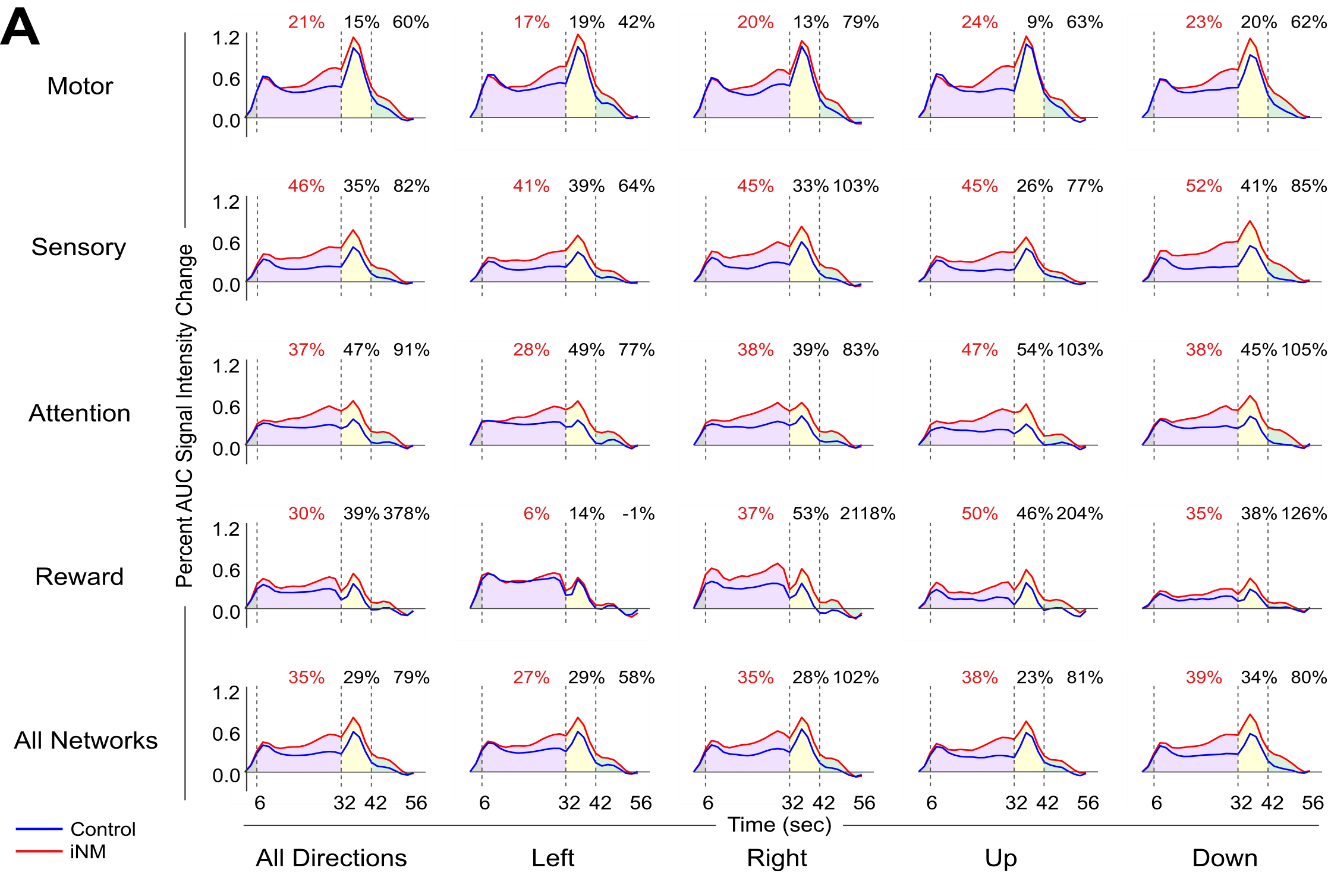

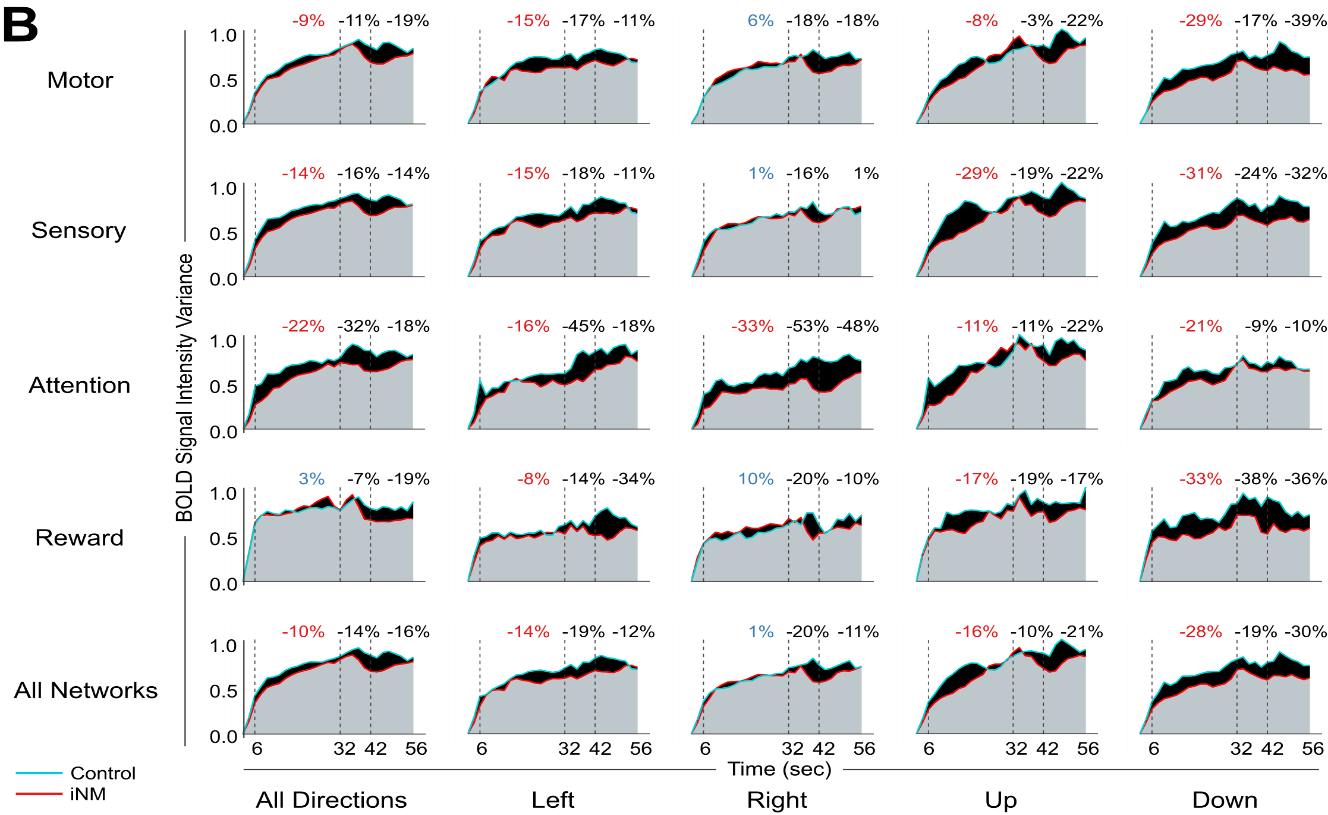

**Extended Data Fig. 8. iNM increases the BOLD magnitude and reduces its variance across TMSC directions and networks.**
iNM (red) overall decreases the mean percent AUC change of the BOLD intensity variance compared to the control (blue) condition for most tongue motor control directions and networks, as a function of time: (1) TMSC interval (6–32 s), (2) swallow (32–42 s), and (3) baseline-tongue-at-rest (42–56 s). Negative values indicate iNM-induced reductions in variance; positive values indicate increases. ROIs specific to each movement direction were analyzed. For the analysis across all directions, missing ROIs were set to zeros, though the results were unchanged when instead substituted with random numbers drawn from a normal distribution with a mean of 0 and standard deviation of 0.05. Notably, iNM effects persisted into the swallow and rest intervals, despite the absence of active modulation, indicating functional carryover.

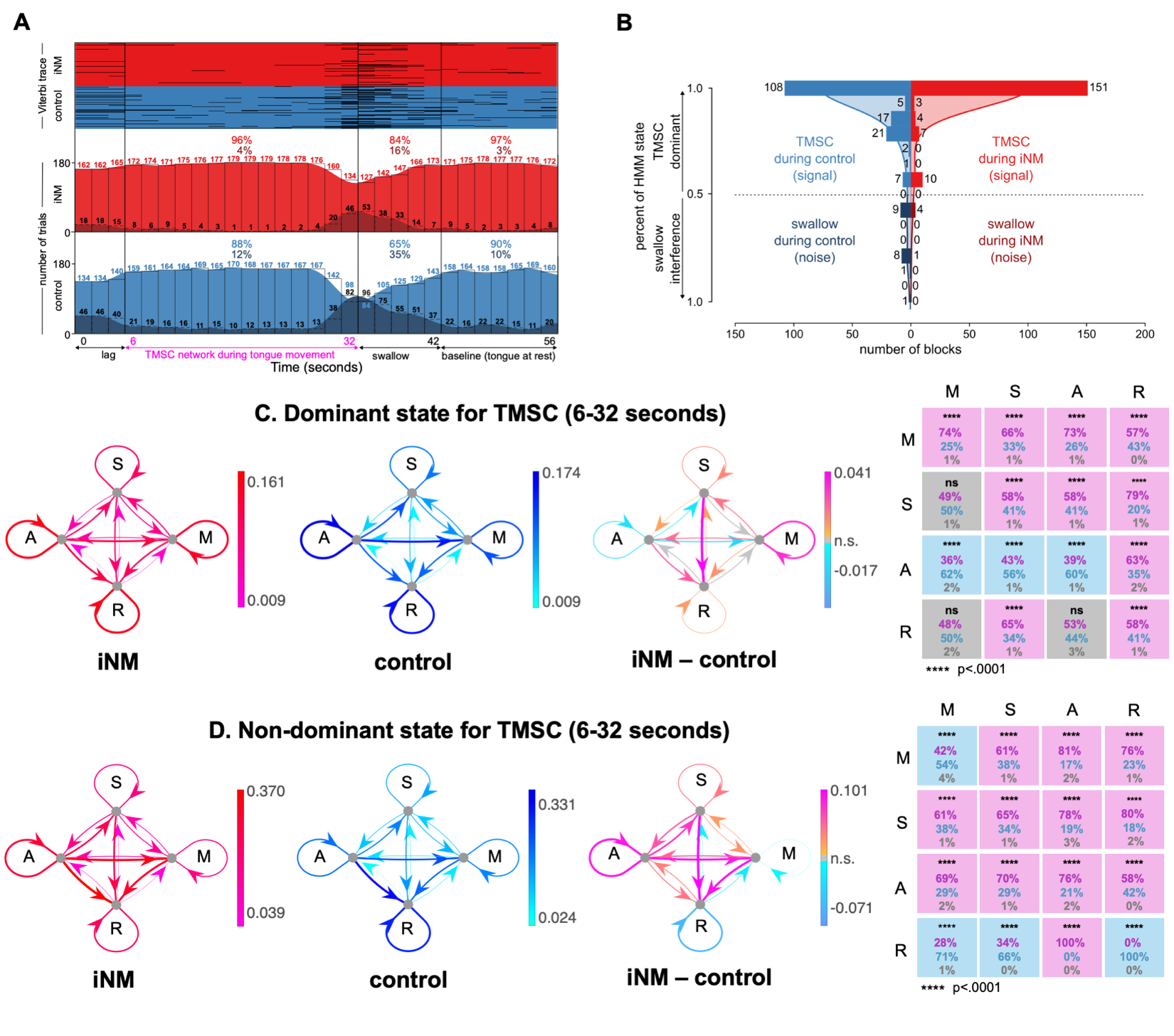

**Extended Data Fig. 9. Effect of iNM on network interactions across Left tongue motor control directions elucidated via temporal causal modeling: A.** Our unsupervised, autoregressive hidden Markov model (ARHMM) was fit to the brain areas that selected for Left tongue movement direction (6-32 seconds). Model inference was then performed on the entire timeseries (0-56 seconds). (top) Viterbi trace for iNM (red) and control (blue) trials that characterize each brain state during Left TMSC, swallow, and baseline intervals for each condition. (bottom) Histogram of the number of trials corresponding to Left TMSC (light red/blue) or swallow (dark red/blue) brain state for each time repetition, divided into the iNM (red) and control (blue) conditions. Percentages correspond to the fraction of trials within each brain state across the Left TMSC, swallow, or baseline intervals. **B.** The histogram compares the signal (Left TMSC state, red/blue) and noise (swallow state, dark red/dark blue) by displaying the number of trials in each bin that correspond to each of the two states (total trials (iNM vs. control): 151 vs. 108 (Left TMSC state), 5 vs. 19 (swallow state). Shaded curves represent the kernel density estimate (KDE) fit to the binned data for the iNM and control conditions. The KDE was calculated by the summation of normal distributions centered at each data point, followed by normalization, to generate a smoothed estimate of the histogram bins. KDE bandwidth was estimated via Scott’s method, and boundary corrections at 0 and 1 employed reflection of the data across the boundaries. **C-D.** For the dominant (Left TMSC) or non-dominant state, the strength of ROI-to-ROI connections within networks (M = Motor network, S = Sensory network, A = Attention network, R = Reward network) is quantified as the mean absolute value of the autoregressive coefficients during the iNM (fuchsia/red) or control (light/dark blue) condition, or the difference in absolute values between conditions (orange/fuchsia and light/dark blue indicate increased/decreased during iNM, respectively). At far right, significant increases (fuchsia), decreases (light blue), or insignificant differences (gray) in these network-to-network interactions are shown for iNM compared to control. Percentages indicate the fraction of region-to-region interactions within each network-to-network connection that are increased (fuchsia), decreased (light blue), or insignificant (gray) in iNM compared to control. Correction for multiple comparisons used an FDR of 0.05.

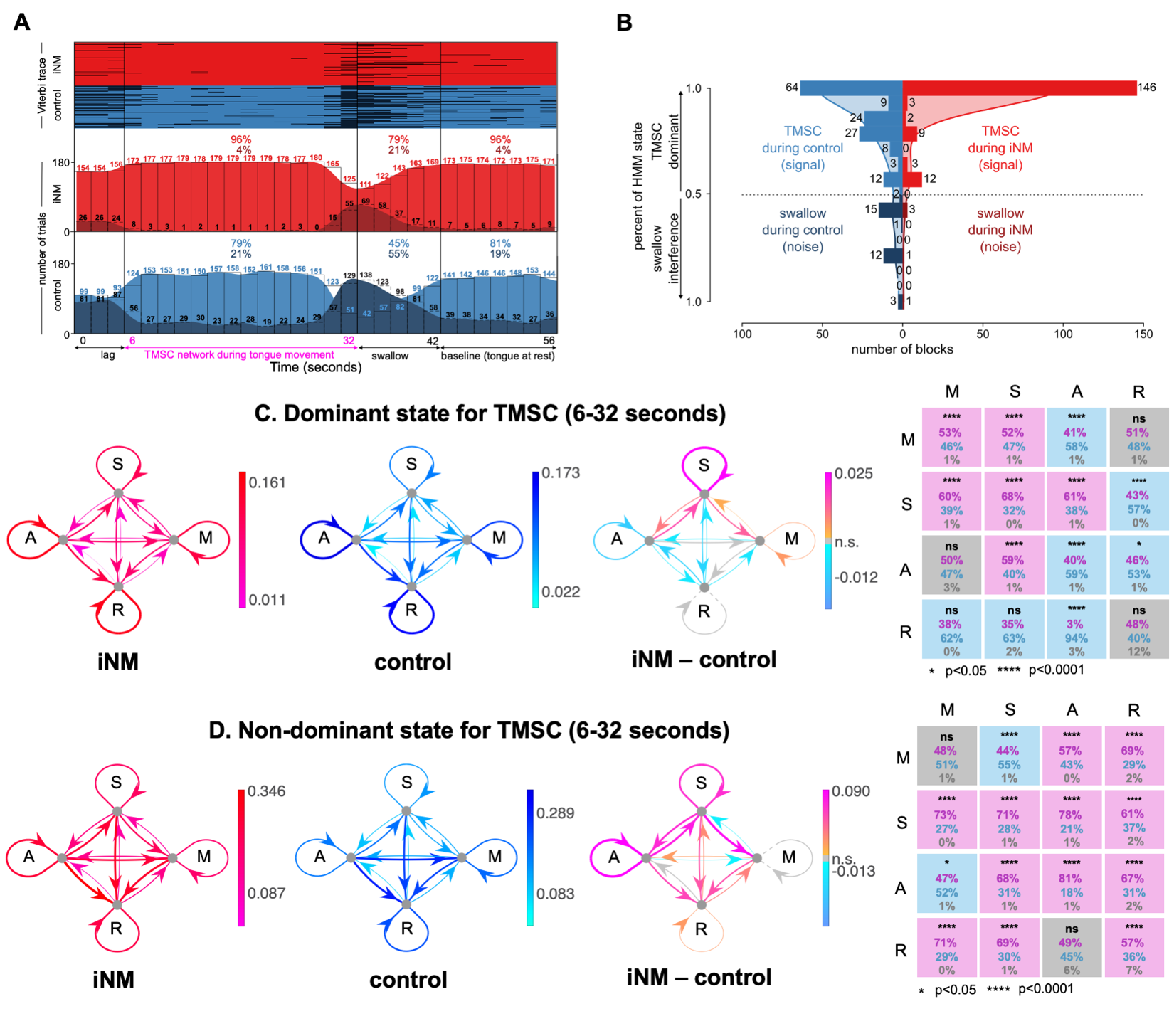

**Extended Data Fig. 10. Effect of iNM on network interactions across Right tongue motor control directions elucidated via temporal causal modeling: A.** Our unsupervised, autoregressive hidden Markov model (ARHMM) was fit to the brain areas that selected for Right tongue movement direction (6-32 seconds). Model inference was then performed on the entire timeseries (0-56 seconds). (top) Viterbi trace for iNM (red) and control (blue) trials that characterize each brain state during Right TMSC, swallow, and baseline intervals for each condition. (bottom) Histogram of the number of trials corresponding to Right TMSC (light red/blue) or swallow (dark red/blue) brain state for each time repetition, divided into the iNM (red) and control (blue) conditions. Percentages correspond to the fraction of trials within each brain state across the Right TMSC, swallow, or baseline intervals. **B.** The histogram compares the signal (Right TMSC state, red/blue) and noise (swallow state, dark red/dark blue) by displaying the number of trials in each bin that correspond to each of the two states (total trials (iNM vs. control): 146 vs. 64 (Right TMSC state), 5 vs. 33 (swallow state)). Shaded curves represent the kernel density estimate (KDE) fit to the binned data for the iNM and control conditions. The KDE was calculated by the summation of normal distributions centered at each data point, followed by normalization, to generate a smoothed estimate of the histogram bins. KDE bandwidth was estimated via Scott’s method, and boundary corrections at 0 and 1 employed reflection of the data across the boundaries. **C-D.** For the dominant (Right TMSC) or non-dominant state, the strength of ROI-to-ROI connections within networks (M = Motor network, S = Sensory network, A = Attention network, R = Reward network) is quantified as the mean absolute value of the autoregressive coefficients during the iNM (fuchsia/red) or control (light/dark blue) condition, or the difference in absolute values between conditions (orange/fuchsia and light/dark blue indicate increased/decreased during iNM, respectively). At far right, significant increases (fuchsia), decreases (light blue), or insignificant differences (gray) in these network-to-network interactions are shown for iNM compared to control. Percentages indicate the fraction of region-to-region interactions within each network-to-network connection that are increased (fuchsia), decreased (light blue), or insignificant (gray) in iNM compared to control. Correction for multiple comparisons used an FDR of 0.05.

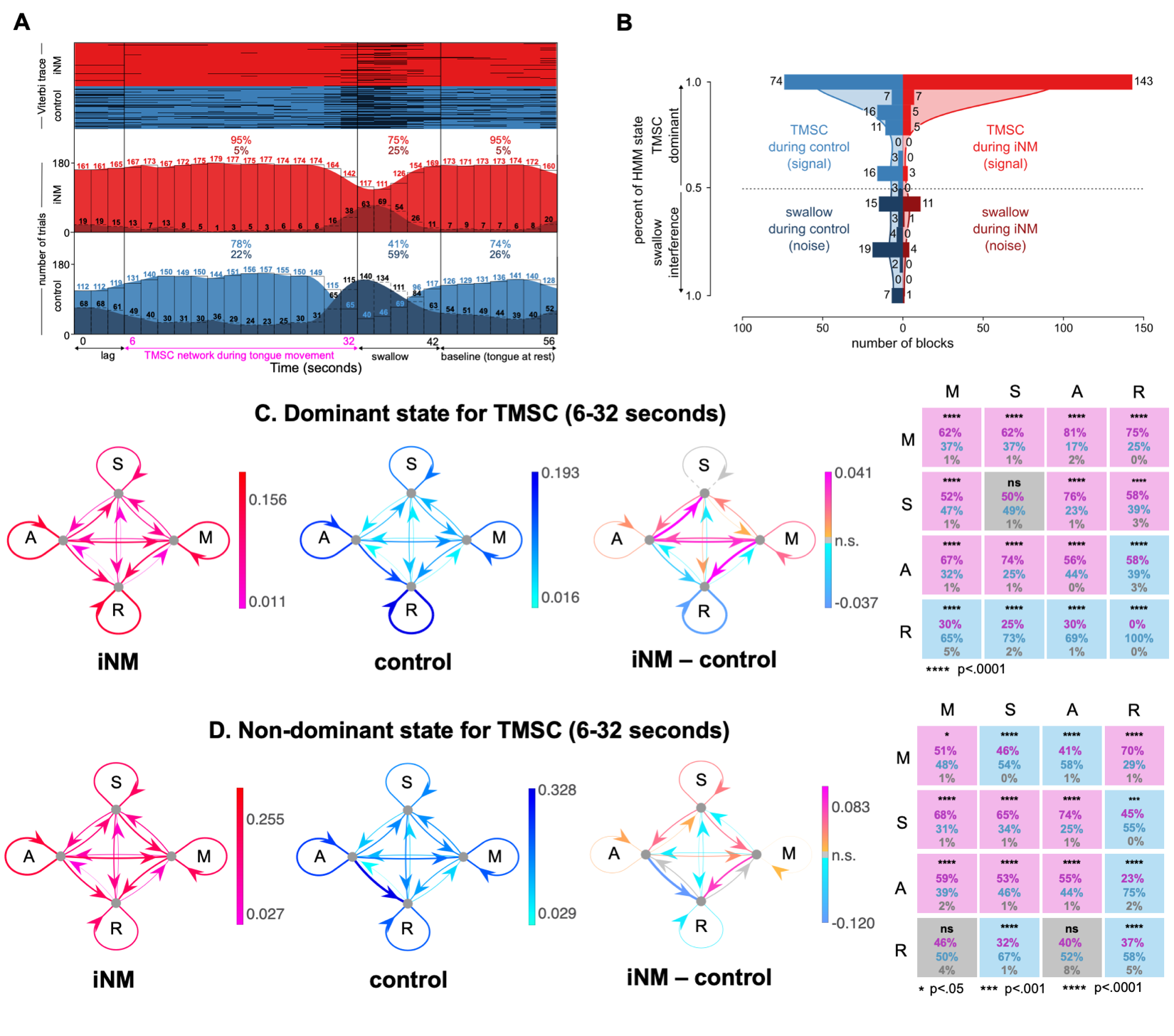

**Extended Data Fig. 11. Effect of iNM on network interactions across Up tongue motor control directions elucidated via temporal causal modeling: A.** Our unsupervised, autoregressive hidden Markov model (ARHMM) was fit to the brain areas that selected for Up tongue movement direction (6-32 seconds). Model inference was then performed on the entire timeseries (0-56 seconds). (top) Viterbi trace for iNM (red) and control (blue) trials that characterize each brain state during Up TMSC, swallow, and baseline intervals for each condition. (bottom) Histogram of the number of trials corresponding to Up TMSC (light red/blue) or swallow (dark red/blue) brain state for each time repetition, divided into the iNM (red) and control (blue) conditions. Percentages correspond to the fraction of trials within each brain state across the Up TMSC, swallow, or baseline intervals. **B.** The histogram compares the signal (Up TMSC state, red/blue) and noise (swallow state, dark red/dark blue) by displaying the number of trials in each bin that correspond to each of the two states (total trials (iNM vs. control): 143 vs. 74 (Up TMSC state), 17 vs. 53 (swallow state)). Shaded curves represent the kernel density estimate (KDE) fit to the binned data for the iNM and control conditions. The KDE was calculated by the summation of normal distributions centered at each data point, followed by normalization, to generate a smoothed estimate of the histogram bins. KDE bandwidth was estimated via Scott’s method, and boundary corrections at 0 and 1 employed reflection of the data across the boundaries. **C-D.** For the dominant (Up TMSC) or non-dominant state, the strength of ROI-to-ROI connections within networks (M = Motor network, S = Sensory network, A = Attention network, R = Reward network) is quantified as the mean absolute value of the autoregressive coefficients during the iNM (fuchsia/red) or control (light/dark blue) condition, or the difference in absolute values between conditions (orange/fuchsia and light/dark blue indicate increased/decreased during iNM, respectively). At far right, significant increases (fuchsia), decreases (light blue), or insignificant differences (gray) in these network-to-network interactions are shown for iNM compared to control. Percentages indicate the fraction of region-to-region interactions within each network-to-network connection that are increased (fuchsia), decreased (light blue), or insignificant (gray) in iNM compared to control. Correction for multiple comparisons used an FDR of 0.05.

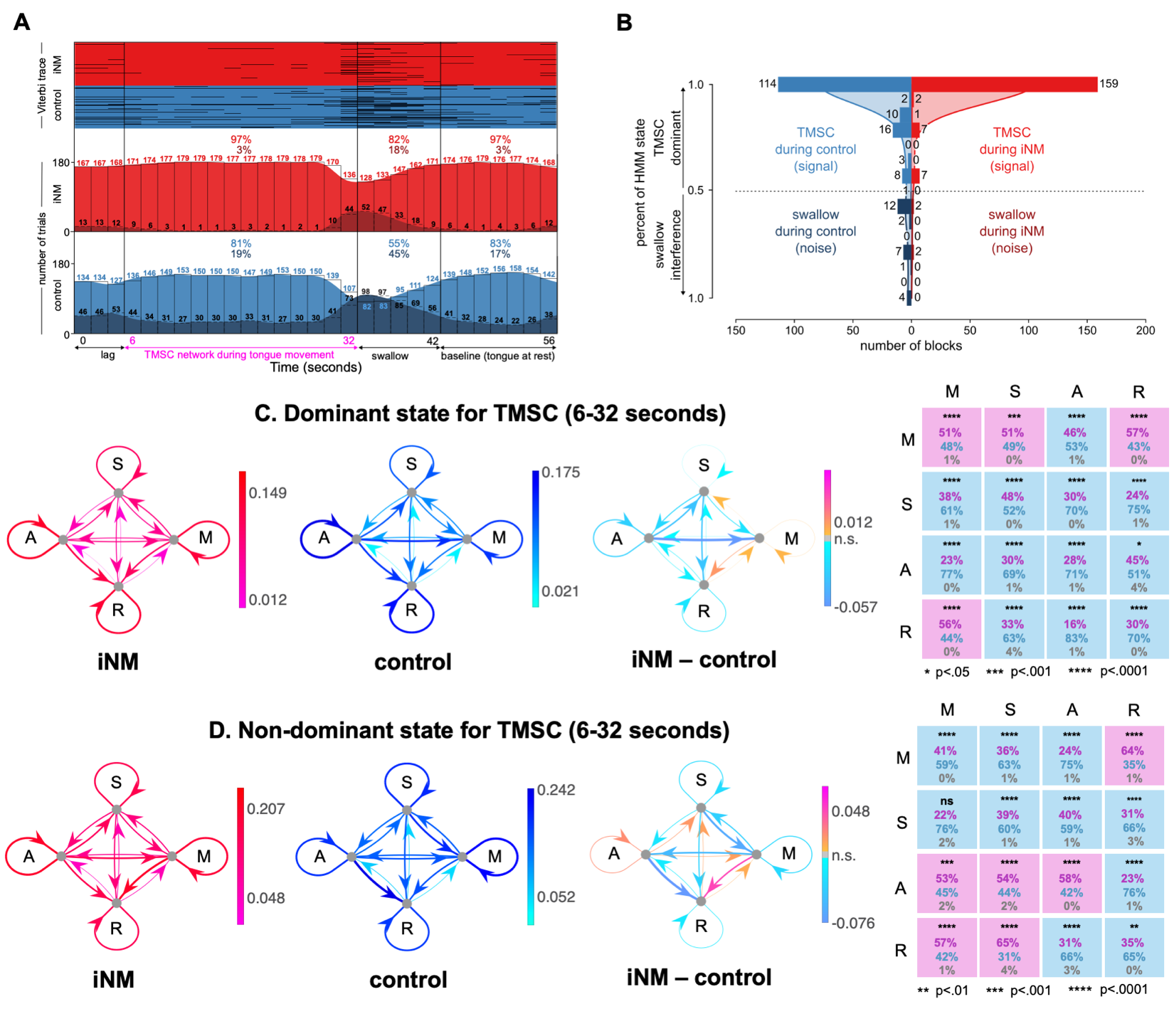

**Extended Data Fig. 12. Effect of iNM on network interactions across Down tongue motor control directions elucidated via temporal causal modeling: A.** Our unsupervised, autoregressive hidden Markov model (ARHMM) was fit to the brain areas that selected for Down tongue movement direction (6-32 seconds). Model inference was then performed on the entire timeseries (0-56 seconds). (top) Viterbi trace for iNM (red) and control (blue) trials that characterize each brain state during Down TMSC, swallow, and baseline intervals for each condition. (bottom) Histogram of the number of trials corresponding to Down TMSC (light red/blue) or swallow (dark red/blue) brain state for each time repetition, divided into the iNM (red) and control (blue) conditions. Percentages correspond to the fraction of trials within each brain state across the Down TMSC, swallow, or baseline intervals. **B.** The histogram compares the signal (Down TMSC state, red/blue) and noise (swallow state, dark red/dark blue) by displaying the number of trials in each bin that correspond to each of the two states (total trials (iNM vs. control): 159 vs. 114 (Down TMSC state), 4 vs. 26 (swallow state). Shaded curves represent the kernel density estimate (KDE) fit to the binned data for the iNM and control conditions. The KDE was calculated by the summation of normal distributions centered at each data point, followed by normalization, to generate a smoothed estimate of the histogram bins. KDE bandwidth was estimated via Scott’s method, and boundary corrections at 0 and 1 employed reflection of the data across the boundaries. **C-D.** For the dominant (Down TMSC) or non-dominant state, the strength of ROI-to-ROI connections within networks (M = Motor network, S = Sensory network, A = Attention network, R = Reward network) is quantified as the mean absolute value of the autoregressive coefficients during the iNM (fuchsia/red) or control (light/dark blue) condition, or the difference in absolute values between conditions (orange/fuchsia and light/dark blue indicate increased/decreased during iNM, respectively). At far right, significant increases (fuchsia), decreases (light blue), or insignificant differences (gray) in these network-to-network interactions are shown for iNM compared to control. Percentages indicate the fraction of region-to-region interactions within each network-to-network connection that are increased (fuchsia), decreased (light blue), or insignificant (gray) in iNM compared to control. Correction for multiple comparisons used an FDR of 0.05.

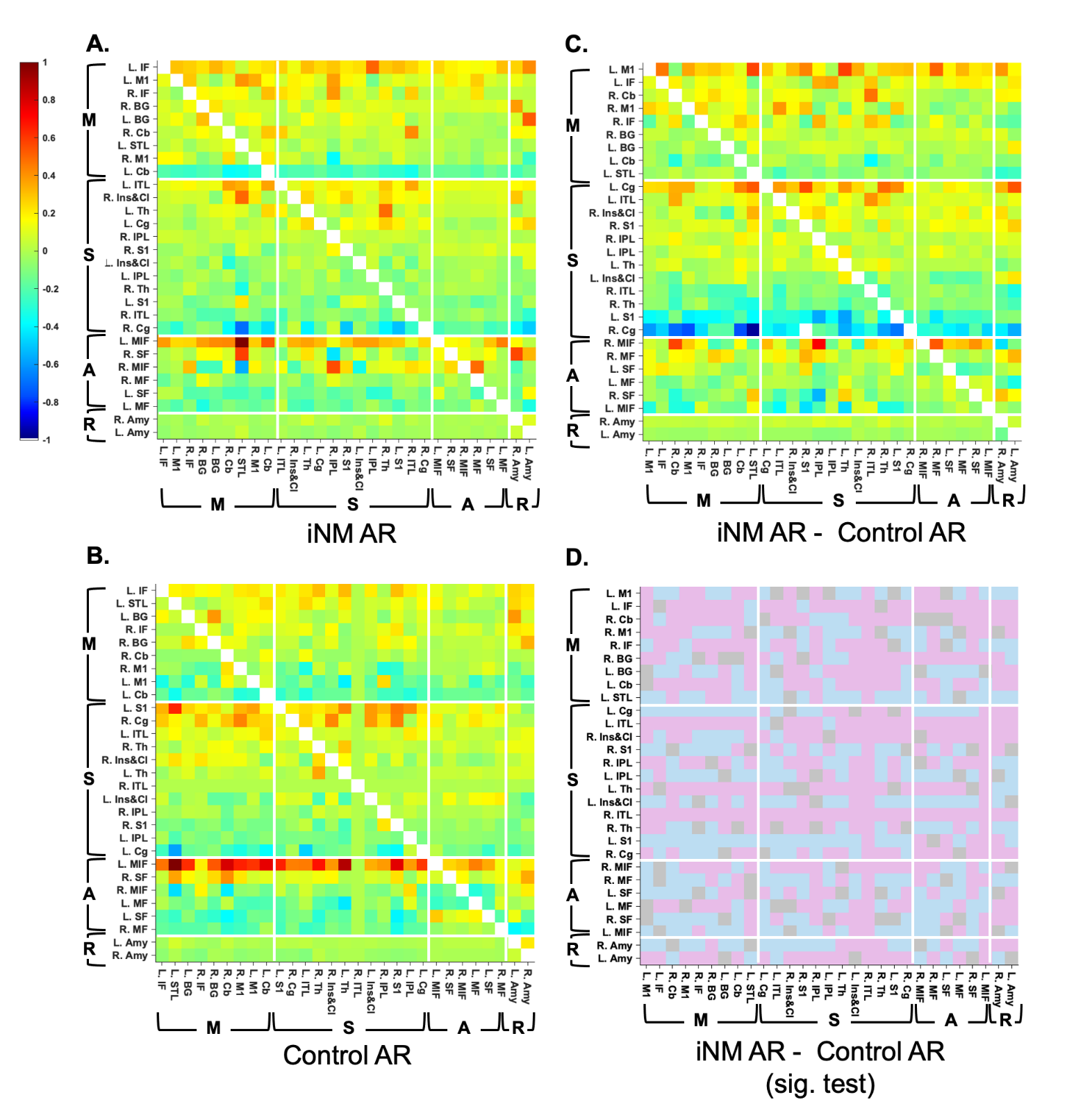

**Extended Data Fig. 13. Autoregressive Hidden Markov model (ARHMM) coefficients reflecting ROI-to-ROI connectivity during tongue movement in the left direction.** The autoregressive coefficient matrix represents directional connectivity from row to column, computed separately for the (A) iNM and (B) control conditions. Heatmaps show values normalized to the range [–1, +1], with ROIs sorted within each network, motor (M), sensory (S), attention (A), reward (R) in descending order based on row-averaged values. (C) Difference matrix showing iNM minus control coefficients. (D) Statistical significance of pairwise differences across 30 subjects (two-tailed paired t-test): pink indicates iNM > control, blue indicates iNM < control, and gray denotes non-significant changes (FDR-corrected).

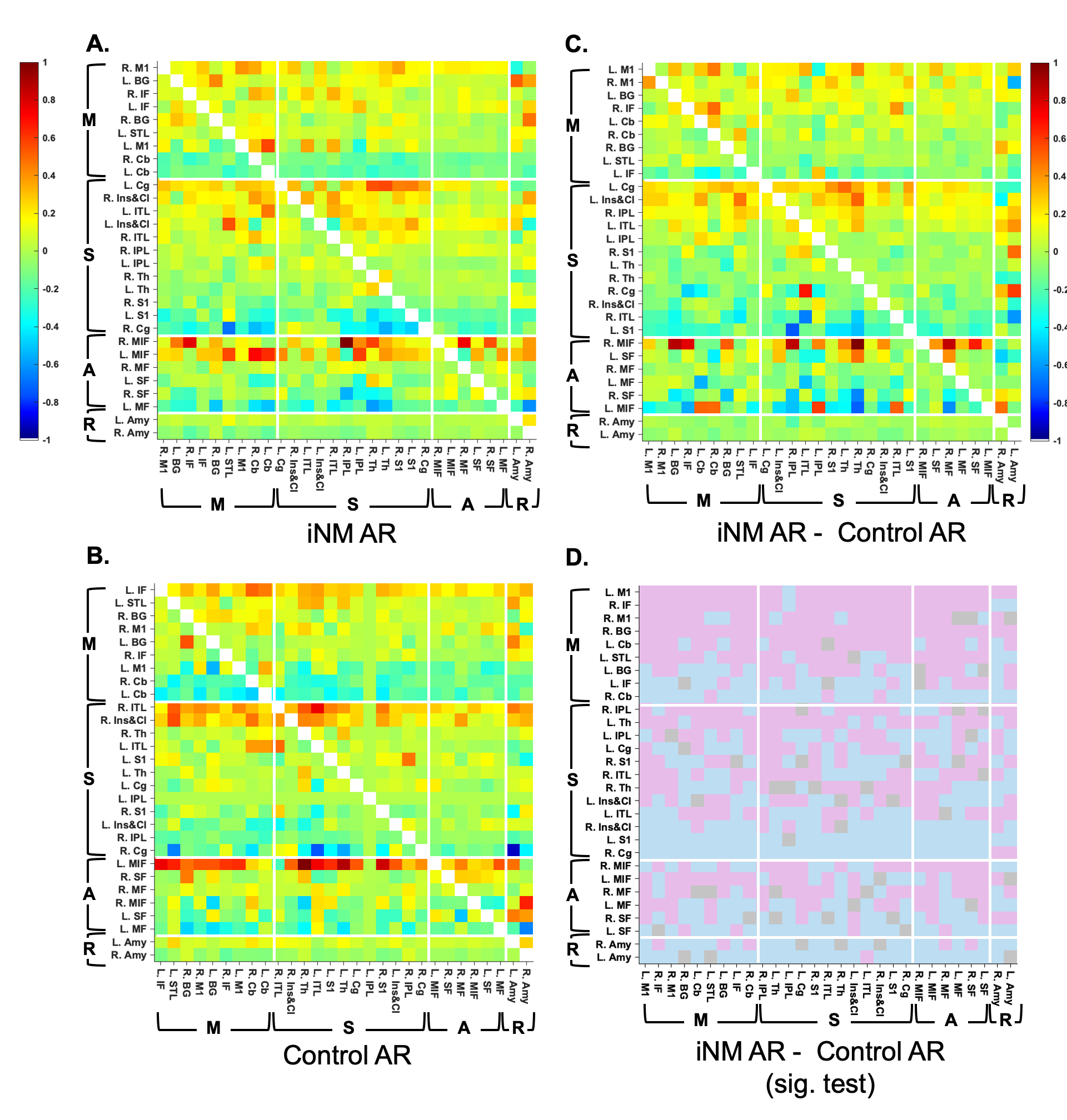
**Extended Data Fig. 14. Autoregressive hidden Markov model (ARHMM) coefficients reflecting ROI-to-ROI connectivity during tongue movement in the right direction.** The autoregressive coefficient matrix represents directional connectivity from row to column, computed separately for the (A) iNM and (B) control conditions. Heatmaps show values normalized to the range [–1, +1], with ROIs sorted within each network, motor (M), sensory (S), attention (A), reward (R) in descending order based on row-averaged values. (C) Difference matrix showing iNM minus control coefficients. (D) Statistical significance of pairwise differences across 30 subjects (two-tailed paired t-test): pink indicates iNM > control, blue indicates iNM < control, and gray denotes non-significant changes (FDR-corrected).

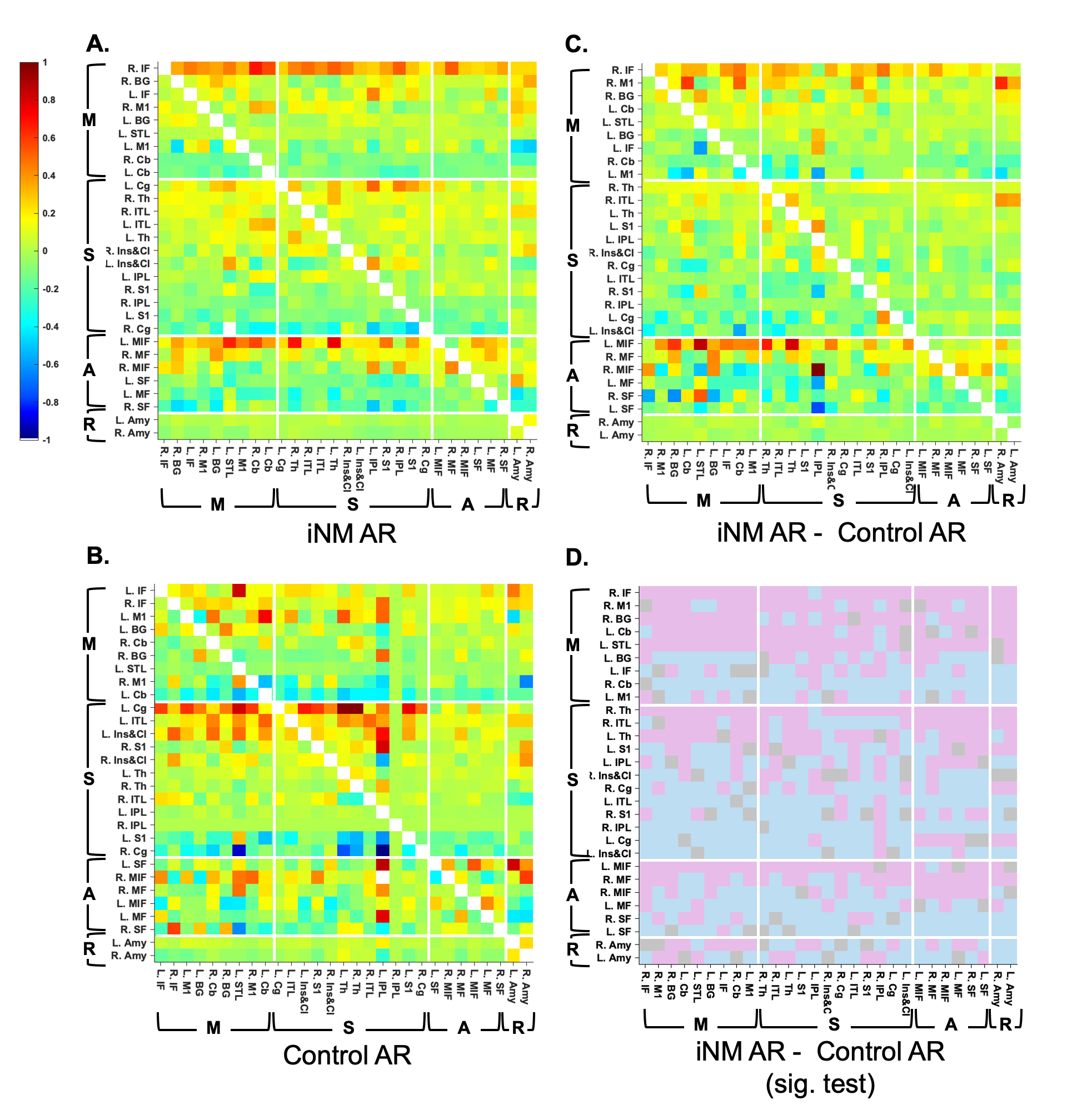

**Extended Data Fig. 15. Autoregressive hidden Markov model (ARHMM) coefficients reflecting ROI-to-ROI connectivity during tongue movement in the up direction.** The autoregressive coefficient matrix represents directional connectivity from row to column, computed separately for the (A) iNM and (B) control conditions. Heatmaps show values normalized to the range [–1, +1], with ROIs sorted within each network, motor (M), sensory (S), attention (A), reward (R) in descending order based on row-averaged values. (C) Difference matrix showing iNM minus control coefficients. (D) Statistical significance of pairwise differences across 30 subjects (two-tailed paired t-test): pink indicates iNM > control, blue indicates iNM < control, and gray denotes non-significant changes (FDR-corrected).

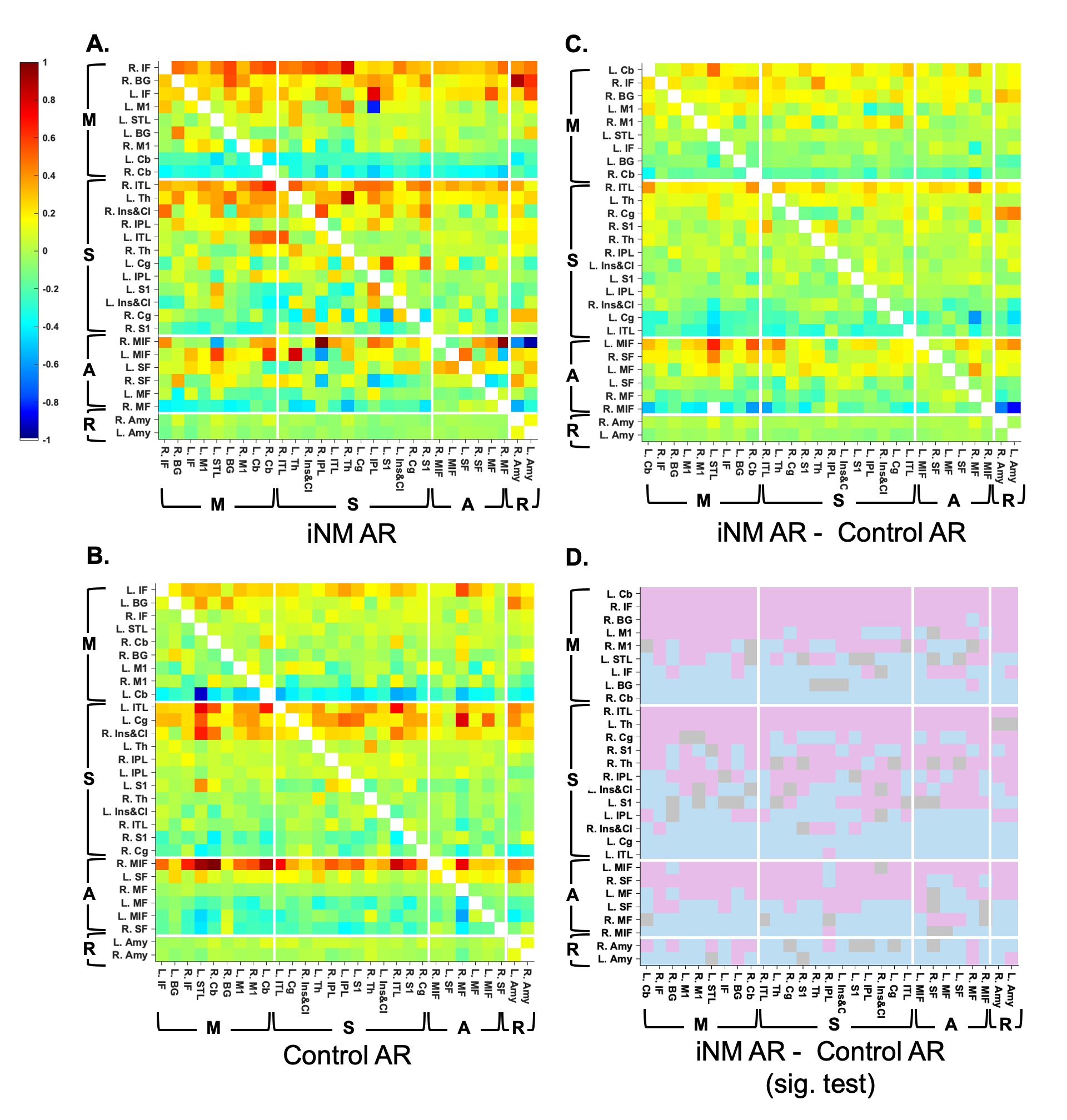

**Extended Data Fig. 16. Autoregressive hidden Markov model (ARHMM) coefficients reflecting ROI-to-ROI connectivity during tongue movement in the down direction.** The autoregressive coefficient matrix represents directional connectivity from row to column, computed separately for the (A) iNM and (B) control conditions. Heatmaps show values normalized to the range [–1, +1], with ROIs sorted within each network, motor (M), sensory (S), attention (A), reward (R) in descending order based on row-averaged values. (C) Difference matrix showing iNM minus control coefficients. (D) Statistical significance of pairwise differences across 30 subjects (two-tailed paired t-test): pink indicates iNM > control, blue indicates iNM < control, and gray denotes non-significant changes (FDR-corrected).

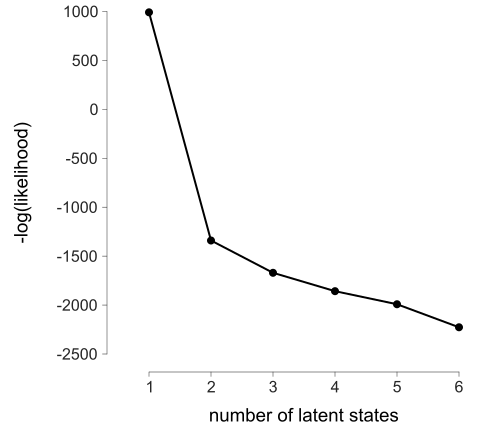

**Extended Data Fig. 17a**. **Model selection for the ARHMM based on latent state number.** The ARHMM was fit to the tongue movement phase (6–32 seconds) of the combined dataset, comprising trials from all directions (left, right, up, down) and both conditions (iNM, control). Leave-one-subject-out cross-validation was used: the model was trained on 29 of 30 subjects during the tongue movement phase and tested on the full timeseries (0–56 seconds) of the held-out subject, repeated across all 30 subjects. The average negative log-likelihood of the test data is shown across different numbers of latent states. An “elbow” at two latent states indicates this model complexity best captures the structure of the data without overfitting.

|  |  | **Network** | | | |
| --- | --- | --- | --- | --- | --- |
| **Direction** | **Flow** | **Motor** | **Sensory** | **Attention** | **Reward** |
| Combined | Input | 0.056 | 0.050 | 0.026 | –0.023 |
|  | Output | 0.049 | 0.041 | 0.031 | –0.013 |
| Left | Input | 0.013 | 0.025 | 0.011 | 0.081 |
|  | Output | 0.074 | 0.059 | –0.018 | 0.016 |
| Right | Input | 0.013 | 0.036 | –0.019 | –0.015 |
|  | Output | 0.004 | 0.035 | –0.001 | –0.023 |
| Up | Input | 0.049 | 0.047 | 0.047 | –0.010 |
|  | Output | 0.097 | 0.029 | 0.062 | –0.055 |
| Down | Input | –0.066 | –0.039 | –0.061 | –0.043 |
|  | Output | 0.014 | –0.072 | –0.124 | –0.026 |

**Extended Data Fig. 17b. Direction-specific network dynamics under individualized neuromodulation (iNM).** For each tongue motor and sensory control (TMSC) direction (left, right, up, down, combined), we quantified ROI-to-ROI connectivity within motor (M), sensory (S), attention (A), and reward (R) networks using an ARHMM. Connectivity strength was defined as the mean absolute value of autoregressive coefficients under iNM (fuchsia/red) and control (light/dark blue) conditions. Difference maps show absolute changes: orange/fuchsia indicates iNM-driven increase and light/dark blue indicates decrease. The Left TMSC direction (dominant state) showed the largest iNM-driven increase in motor and sensory network output, suggesting enhanced top-down modulation specific to the dominant movement pattern. The Down TMSC direction showed reduced input and output across all networks under iNM, indicating a potential suppression or disengagement mechanism. The Up TMSC direction showed increased output particularly in the attention network, suggesting iNM-facilitated recruitment of higher-order control during upward tongue movement. Across directions, combined TMSC input-output patterns reflected broad engagement of motor, sensory, and attention circuits under iNM, while the reward network consistently showed minimal or negative modulation, which reflects its limited role in precision-guided sensorimotor TMSC control.

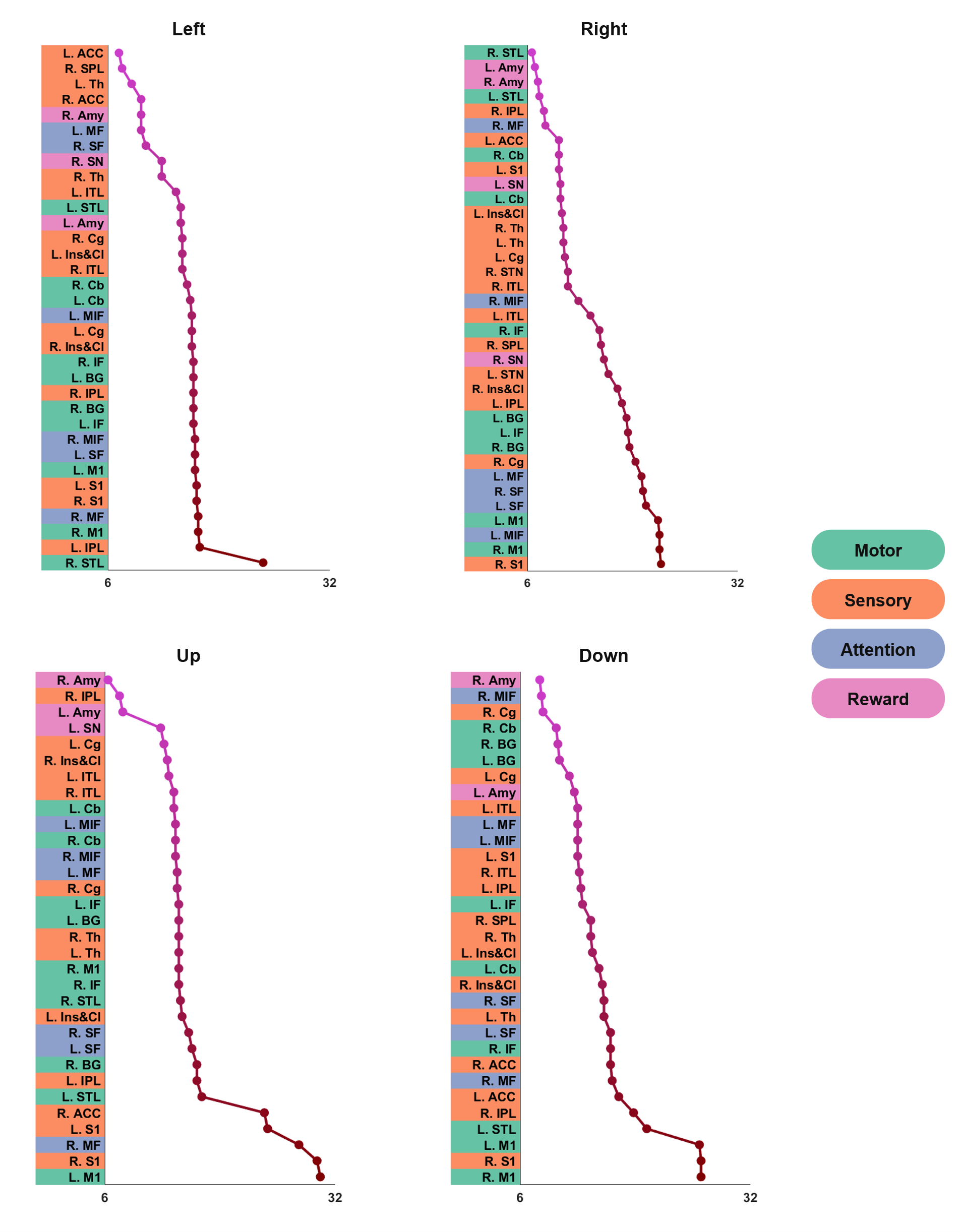

**Extended Data Fig. 18**: **Time of earliest peak ROI activity, quantified by the d’ score for individual tongue movement directions.** Time of the first peak ROI activity, quantified by the d′ sensitivity index, is shown for each tongue movement direction (left, right, up, down). ROIs are ordered by descending peak latency and color-coded by network: motor (green), sensory (orange), attention (blue), and reward (pink). Peak times were identified by first applying cubic interpolation to the d′ score time series, followed by peak detection using the MATLAB’s peakfinder function. The timing of the earliest peak per ROI was then extracted for each condition.

**Table S1a.** Activation peaks of regions showing direction selectivity for Left TMSC under the iNM condition..

| TT_Daemon Atlas Region | BA | Cluster | Cluster Size  (# of Voxels) | | Region Size (# of Voxels) | RAI Peak Coordinates (mm) | | | Peak  Z-score |
| --- | --- | --- | --- | --- | --- | --- | --- | --- | --- |
|  |  |  |  |  |  | x | y | z |  |
| L precentral  (premotor & 1ary motor cortex) | 4, 6 | 1 | 9824 | | 588 | 46.5 | 7.5 | 26.5 | 7.7 |
| R precentral  (premotor & 1ary motor cortex) | 4, 6 | 1 | 9824 | | 502 | -46.5 | 10.5 | 29.5 | 7.3 |
| L postcentral  (1ary somatosensory) | 3 | 1 | 9824 | | 212 | 58.5 | 13.5 | 26.5 | 7.0 |
| R postcentral  (1ary somatosensory) | 3 |  |  |  | 259 | -58.5 | 7.5 | 20.5 | 7.2 |
| L thalamus |  | 1 | 9824 | | 79 | 10.5 | 16.5 | -0.5 | 5.4 |
| R thalamus |  | 1 | 9824 | | 88 | -10.5 | 22.5 | -9.5 | 6.4 |
| L lentiform & putamen |  | 1 | 9824 | | 129 | 25.5 | 1.5 | -3.5 | 7.1 |
| R lentiform & putamen |  | 1 | 9824 | | 158 | -25.5 | 1.5 | -6.5 | 7.3 |
| L insula & claustrum |  | 1 | 9824 | | 246 | 31.5 | -13.5 | 11.5 | 6.3 |
| R insula & claustrum |  | 1 | 9824 | | 312 | -34.5 | 7.5 | 14.5 | 6.5 |
| L inferior frontal | 9 | 1 | 9824 | | 254 | 34.5 | -10.5 | 23.5 | 5.0 |
| R inferior frontal  (operculum) | 45 | 1 | 9824 | | 246 | -31.5 | -25.5 | 8.5 | 6.7 |
| L middle frontal | 10 | 1 | 9824 | | 217 | 31.5 | -43.5 | 11.5 | 4.3 |
| R middle frontal | 10 | 1 | 9824 | | 411 | -31.5 | -40.5 | 23.5 | 5.1 |
| L medial frontal | 24 | 1 | 9824 | | 329 | 7.5 | 4.5 | 47.5 | 5.3 |
| R medial frontal  (premotor cortex) | 6 | 1 | 9824 | | 404 | -4.5 | 7.5 | 53.5 | 5.6 |
| L superior frontal | 10 | 1 | 9824 | | 198 | 16.5 | -43.5 | 23.5 | 4.3 |
| R superior frontal  (anterior) | 9, 32 | 1 | 9824 | | 422 | -13.5 | -19.5 | 44.5 | 6.5 |
| L inferior parietal | 40 | 1 | 9824 | | 199 | 49.5 | 40.5 | 41.5 | 4.3 |
| R inferior parietal | 40 | 1 | 9824 | | 239 | -46.5 | 49.5 | 47.5 | 4.6 |
| R superior parietal |  | 1 | 9824 | | 37 | -25.5 | 70.5 | 20.5 | 3.8 |
| L nodule (lobule IX) |  | 2 | 829 | | 6 | 13.5 | 55.5 | -24.5 | 5.7 |
| L declive (lobule VI – crus 1) |  | 2 | 829 | | 190 | 31.5 | 61.5 | -21.5 | 5.5 |
| L uvula (lobule VII) |  | 2 | 829 | | 29 | 34.5 | 67.5 | -24.5 | 4.6 |
| R culmen (lobule VI) |  | 1 | 9824 | | 65 | -16.5 | 55.5 | -21.5 | 4.8 |
| L inferior temporal | 37 | 2 | 829 | | 87 | 46.5 | 61.5 | -15.5 | 6.3 |
| R inferior temporal |  | 1 | 9824 | | 80 | -43.5 | 58.5 | -12.5 | 5.7 |
| L cingulate (middle) | 24, 32 | 1 | 9824 | | 278 | 7.5 | 4.5 | 47.5 | 5.3 |
| R cingulate (middle) |  | 1 | 9824 | | 313 | -4.5 | 7.5 | 32.5 | 3.6 |
| L anterior cingulate | 24, 32 | 1 | 9824 | | 37 | 16.5 | -34.5 | 11.5 | 4.1 |
| R anterior cingulate |  | 1 | 9824 | | 113 | -13.5 | -34.5 | 14.5 | 4.8 |
| R superior temporal | 38 | 1 | 9824 | | 86 | -37.5 | -7.5 | -21.5 | 3.6 |
| R substantia nigra |  | 1 | 9824 | | 11 | -10.5 | 19.5 | -9.5 | 5.8 |
| L amygdala | 28 | 1 | 9824 | | 8 | 25.5 | 1.5 | -9.5 | 5.1 |
| R amygdala | 28 | 1 | 9824 | | 47 | -25.5 | 1.5 | -9.5 | 6.7 |

**Table S1b.** Activation peaks of regions showing direction selectivity for Left TMSC under the Control-No-iNM condition.

| TT_Daemon Atlas Region | BA | Cluster | Cluster Size  (# of Voxels) | Region Size  (# of Voxels) | RAI Peak Coordinates (mm) | | | Peak  Z-score |
| --- | --- | --- | --- | --- | --- | --- | --- | --- |
|  |  |  |  |  | x | y | z |  |
| L precentral  (premotor & 1ary motor cortex) | 6, 4 | 1 | 6211 | 504 | 49.5 | 7.5 | 26.5 | 7.9 |
| R precentral  (premotor & 1ary motor cortex) | 6, 4 | 1 | 6211 | 408 | -58.5 | 4.5 | 29.5 | 7.7 |
| L postcentral  (1ary somatosensory) | 3/1/2 | 1 | 6211 | 174 | 43.5 | 16.5 | 29.5 | 5.8 |
| R postcentral  (1ary somatosensory) | 3/1/2 | 1 | 6211 | 218 | -58.5 | 13.5 | 29.5 | 6.1 |
| L thalamus  (subthalamic nucleus) |  | 4 | 56 | 29 | 10.5 | 16.5 | -0.5 | 5.8 |
| R thalamus  (subthalamic nucleus) |  | 1 | 6211 | 36 | -13.5 | 16.5 | -0.5 | 6.1 |
| L lentiform, putamen |  | 1 | 6211 | 136 | 25.5 | 1.5 | -6.5 | 6.9 |
| R lentiform, putamen |  | 1 | 6211 | 165 | -25.5 | 1.5 | -6.5 | 7.1 |
| L insula & claustrum |  | 1 | 6211 | 230 | 34.5 | 7.5 | 14.5 | 6.5 |
| R insula & claustrum |  | 1 | 6211 | 254 | -34.5 | 7.5 | 14.5 | 6.9 |
| L middle frontal | 6, 8 | 1 | 6211 | 146 | 31.5 | -13.5 | 44.5 | 4.4 |
| R middle frontal | 10 | 1 | 6211 | 117 | -31.5 | -43.5 | 2.5 | 4.3 |
| L medial frontal  (supplementary motor area) | 8, 9 | 1 | 6211 | 296 | 7.5 | -25.5 | 47.5 | 5.3 |
| R medial frontal | 9 | 1 | 6211 | 323 | -22.5 | -34.5 | 23.5 | 5.3 |
| L superior medial frontal | 9 | 1 | 6211 | 225 | 10.5 | -46.5 | 35.5 | 4.1 |
| R superior posterior frontal | 10 | 1 | 6211 | 311 | -19.5 | -49.5 | 23.5 | 4.7 |
| L culmen (lobule VI) |  | 2 | 259 | 24 | 13.5 | 55.5 | -24.5 | 5.8 |
| R culmen (lobule VI) |  | 6 | 40 | 6 | -16.5 | 58.5 | -21.5 | 4.5 |
| L cingulate (caudal) | 24 | 1 | 6211 | 211 | 7.5 | 4.5 | 47.5 | 5.8 |
| R cingulate (caudal) | 24 | 1 | 6211 | 210 | -10.5 | -7.5 | 47.5 | 5.1 |
| L anterior cingulate |  | 1 | 6211 | 71 | 16.5 | -19.5 | 20.5 | 5.5 |
| R anterior cingulate |  | 1 | 6211 | 80 | -16.5 | -31.5 | 2.5 | 4.6 |
| L superior temporal | 22 | 3 | 162 | 156 | 37.5 | 25.5 | 5.5 | 5.4 |
| R superior temporal | 22 | 1 | 6211 | 195 | -52.5 | 10.5 | 5.5 | 5.1 |
| R parahippocampal (uncus) |  | 1 | 6211 | 48 | -31.5 | 4.5 | -24.5 | 3.6 |

**Table S2a.** Activation peaks of regions showing direction selectivity for Right TMSC under the iNM condition.

| TT_Daemon Atlas Region | BA | Cluster | Cluster Size  (# of Voxels) | Region Size (# of Voxels) | RAI Peak Coordinates (mm) | | | Peak  Z-score |
| --- | --- | --- | --- | --- | --- | --- | --- | --- |
|  |  |  |  |  | x | y | z |  |
| L precentral  (premotor & 1ary motor) | 6 | 1 | 9467 | 566 | 46.5 | 7.5 | 26.5 | 7.5 |
| R precentral  (premotor & 1ary motor) | 6 | 1 | 9467 | 484 | -46.5 | 10.5 | 26.5 | 7.2 |
| L postcentral  (1ary somatosensory) | 3 | 1 | 9467 | 196 | 55.5 | 10.5 | 44.5 | 3.8 |
| R postcentral  (1ary somatosensory) | 3 | 1 | 9467 | 261 | -58.5 | 10.5 | 20.5 | 6.6 |
| L thalamus |  | 1 | 9467 | 105 | 10.5 | 16.5 | 2.5 | 5.9 |
| R thalamus |  | 1 | 9467 | 101 | -10.5 | 16.5 | -0.5 | 6.0 |
| L lentiform & putamen |  | 1 | 9467 | 165 | 25.5 | 1.5 | -3.5 | 6.7 |
| R lentiform & putamen |  | 1 | 9467 | 126 | -25.5 | 1.5 | -6.5 | 6.4 |
| L insula & claustrum |  | 1 | 9467 | 305 | 34.5 | -1.5 | 5.5 | 5.5 |
| R insula & claustrum |  | 1 | 9467 | 239 | -34.5 | 7.5 | 14.5 | 6.1 |
| L inferior frontal (operculum) | 44, 45 | 1 | 9467 | 244 | 31.5 | -22.5 | 11.5 | 6.5 |
| R inferior frontal | 45 | 1 | 9467 | 260 | -31.5 | -22.5 | 8.5 | 6.2 |
| L middle frontal | 10, 46 | 1 | 9467 | 168 | 31.5 | -46.5 | 20.5 | 4.6 |
| R middle frontal | 9, 10, 46 | 1 | 9467 | 331 | -34.5 | -31.5 | 26.5 | 5.2 |
| L medial frontal  (supplementary motor area) | 6 | 1 | 9467 | 294 | 7.5 | 4.5 | 50.5 | 5.7 |
| R medial frontal  (supplementary motor area) | 6 | 1 | 9467 | 331 | -4.5 | 7.5 | 53.5 | 5.6 |
| L superior frontal | 10 | 1 | 9467 | 169 | 25.5 | -37.5 | 5.5 | 4.2 |
| R superior frontal | 8, 9 | 1 | 9467 | 387 | -7.5 | -31.5 | 50.5 | 5.7 |
| L nodule & dentate (lobule IX) |  | 2 | 537 | 29 | 13.5 | 55.5 | -24.5 | 4.7 |
| L culmen (lobule VI) |  | 2 | 537 | 50 | 34.5 | 55.5 | -18.5 | 4.4 |
| R culmen & dentate (lobule IV-V) |  | 1 | 9467 | 145 | -16.5 | 58.5 | -21.5 | 5.7 |
| R declive |  | 1 | 9467 | 258 | -28.5 | 82.5 | -15.5 | 5.1 |
| L inferior parietal lobule | 40 | 1 | 9467 | 56 | 43.5 | 16.5 | 29.5 | 6.0 |
| R inferior parietal lobule | 40 | 1 | 9467 | 297 | -49.5 | 43.5 | 44.5 | 4.9 |
| R superior parietal lobule | 7 | 1 | 9467 | 32 | -22.5 | 67.5 | 41.5 | 4.2 |
| L inferior temporal  (fusiform) | 37 | 2 | 537 | 76 | 43.5 | 58.5 | -15.5 | 5.6 |
| R inferior temporal  (fusiform) |  | 1 | 9467 | 107 | -49.5 | 55.5 | -0.5 | 3.9 |
| L middle cingulate | 24 | 1 | 9467 | 268 | 10.5 | -10.5 | 35.5 | 4.8 |
| R middle cingulate | 24, 32 | 1 | 9467 | 316 | -10.5 | -16.5 | 38.5 | 4.7 |
| L superior temporal | 28, 36 | 1 | 9467 | 208 | 37.5 | -13.5 | -21.5 | 4.0 |
| R middle & superior temporal | 22 | 1 | 9467 | 91 | -46.5 | 25.5 | -3.5 | 4.0 |
| L substantia nigra |  | 1 | 9467 | 11 | 13.5 | 19.5 | -3.5 | 4.6 |
| R substantia nigra |  | 1 | 9467 | 9 | -11.5 | 19.5 | -6.5 | 4.5 |
| L superior temporal lobe (uncus) | 28, 36 | 1 | 9467 | 20 | 28.5 | 4.5 | -24.5 | 4.0 |
| L amygdala |  | 1 | 9467 | 14 | 25.5 | 1.5 | -9.5 | 5.8 |
| R amygdala |  | 1 | 9467 | 9 | -25.5 | 1.5 | -9.5 | 5.4 |

**Table S2b.** Activation peaks of regions showing direction selectivity for Right TMSC under the Control-No-iNM condition.

| TT_Daemon Atlas Region | BA | Cluster | Cluster Size  (# of Voxels) | Region Size (# of Voxels) | RAI Peak Coordinates (mm) | | | Peak  Z-score |
| --- | --- | --- | --- | --- | --- | --- | --- | --- |
|  |  |  |  |  | x | y | z |  |
| L precentral  (premotor & 1ary motor) | 4, 6 | 1 | 6169 | 507 | 46.5 | 7.5 | 26.5 | 7.8 |
| R precentral  (premotor & 1ary motor) | 4, 6 | 1 | 6169 | 380 | -58.5 | 4.5 | 29.5 | 7.3 |
| L postcentral  (1ary somatosensory) | 3/1/2 | 1 | 6169 | 137 | -49.5 | 10.5 | 23.5 | 7.0 |
| R postcentral  (1ary somatosensory) | 3/1/2 | 1 | 6169 | 197 | -61.5 | 7.5 | 17.5 | 6.9 |
| L thalamus  (subthalamic nucleus) |  | 7 | 34 | 3 | 10.5 | 16.5 | -0.5 | 5.8 |
| R thalamus  (subthalamic nucleus) |  | 8 | 21 | 2 | -13.5 | 16.5 | -0.5 | 5.1 |
| L lentiform & putamen |  | 1 | 6169 | 129 | 25.5 | 1.5 | -6.5 | 7.2 |
| R lentiform & putamen |  | 1 | 6169 | 86 | -25.5 | 1.5 | -6.5 | 7.0 |
| L insula & claustrum |  | 1 | 6169 | 189 | 34.5 | 7.5 | 14.5 | 5.2 |
| R insula & claustrum |  | 1 | 6169 | 214 | -34.5 | 7.5 | 14.5 | 5.9 |
| L inferior frontal (operculum) | 9 | 1 | 6169 | 80 | 43.5 | -10.5 | 26.5 | 3.8 |
| L middle frontal | 9, 46 | 1 | 6169 | 181 | 28.5 | -19.5 | 20.5 | 5.7 |
| R middle frontal | 10, 11 | 1 | 6169 | 103 | -34.5 | -49.5 | 2.5 | 3.4 |
| L medial frontal | 9 | 1 | 6169 | 262 | 22.5 | -28.5 | 26.5 | 5.2 |
| R medial frontal | 9 | 1 | 6169 | 343 | -16.5 | -34.5 | 29.5 | 5.0 |
| L superior frontal | 10, 12 | 1 | 6169 | 262 | 28.5 | -43.5 | 8.5 | 4.7 |
| R superior frontal | 9 | 1 | 6169 | 293 | -16.5 | -46.5 | 26.5 | 5.7 |
| L nodule & dentate (lobule IX) |  | 4 | 49 | 37 | 13.5 | 55.5 | -24.5 | 5.9 |
| R culmen (lobule VI) |  | 2 | 292 | 40 | -16.5 | 58.5 | -21.5 | 5.5 |
| R tuber (lobule VIIB) |  | 2 | 292 | 3 | -49.5 | 67.5 | -24.5 | 3.9 |
| L inferior temporal (fusiform) | 37 | 3 | 129 | 37 | 43.5 | 58.5 | -15.5 | 3.4 |
| R inferior temporal (fusiform) | 20, 37 | 2 | 292 | 34 | -37.5 | 58.5 | -12.5 | 4.1 |
| L middle cingulate  (supplementary motor area) | 24, 32 | 1 | 6169 | 272 | 4.5 | 4.5 | 47.5 | 5.6 |
| R cingulate  (supplementary motor area) | 24 | 1 | 6169 | 324 | -7.5 | 4.5 | 47.5 | 5.1 |
| L anterior cingulate |  | 1 | 6169 | 66 | 10.5 | -31.5 | 11.5 | 4.3 |
| L superior temporal | 38 | 1 | 6169 | 213 | 34.5 | -4.5 | -21.5 | 4.2 |
| R superior temporal | 41, 42 | 1 | 6169 | 76 | -43.5 | 28.5 | 8.5 | 3.8 |
| L amygdala | 28 | 1 | 6169 | 29 | 25.5 | 1.5 | -9.5 | 6.1 |
| R amygdala | 28 | 1 | 6169 | 17 | -25.5 | 1.5 | -9.5 | 6.4 |

**Table S3a.** Activation peaks of regions showing direction selectivity for Up TMSC under the iNM condition.

| TT_Daemon Atlas Region | BA | Cluster | Cluster Size  (# of Voxels) | Region Size  (# of Voxels) | RAI Peak Coordinates (mm) | | | Peak  Z-score |
| --- | --- | --- | --- | --- | --- | --- | --- | --- |
|  |  |  |  |  | x | y | z |  |
| L precentral  (premotor & 1ary motor cortex) | 4 | 1 | 8400 | 538 | 58.5 | 10.5 | 26.5 | 7.8 |
| R precentral  (premotor & 1ary motor cortex) | 6 | 1 | 8400 | 514 | -49.5 | 7.5 | 29.5 | 7.2 |
| L postcentral  (caudal – 1ary somatosensory) | 3, 2, 1 | 1 | 8400 | 203 | 55.5 | 25.5 | 35.5 | 4.4 |
| R postcentral  (rostral – 1ary somatosensory) | 3a | 1 | 8400 | 272 | -58.5 | 7.5 | 20.5 | 7.2 |
| L thalamus |  | 4 | 32 | 15 | 10.5 | 16.5 | -0.5 | 4.2 |
| R thalamus |  | 6 | 17 | 12 | -10.5 | 16.5 | -0.5 | 4.8 |
| L lentiform & putamen |  | 3 | 131 | 80 | 25.5 | 1.5 | -6.5 | 6.1 |
| R lentiform & putamen |  | 1 | 8400 | 113 | -28.5 | 4.5 | -6.5 | 6.2 |
| L insula |  | 1 | 8400 | 166 | 31.5 | -25.5 | 11.5 | 5.7 |
| R insula |  | 1 | 8400 | 215 | -40.5 | 4.5 | 17.5 | 5.7 |
| L inferior frontal | 9 | 1 | 8400 | 208 | 55.5 | -7.5 | 26.5 | 7.0 |
| R inferior frontal | 10 | 1 | 8400 | 311 | -31.5 | -22.5 | 8.5 | 6.5 |
| L middle frontal | 9 | 1 | 8400 | 156 | 25.5 | -22.5 | 26.5 | 4.6 |
| R middle frontal | 10 | 1 | 8400 | 396 | -31.5 | -37.5 | 17.5 | 5.3 |
| L medial frontal | 9 | 1 | 8400 | 217 | 25.5 | -34.5 | 20.5 | 5.3 |
| R medial frontal | 6 | 1 | 8400 | 376 | -1.5 | 7.5 | 53.5 | 5.1 |
| L superior frontal | 8, 9 | 1 | 8400 | 75 | 10.5 | -28.5 | 44.5 | 4.6 |
| R superior frontal | 9 | 1 | 8400 | 290 | -10.5 | -46.5 | 32.5 | 4.6 |
| L culmen (lobule IV-V crus 1) |  | 2 | 833 | 101 | 34.5 | 49.5 | -18.5 | 5.3 |
| L declive (lobule VI) |  | 2 | 833 | 183 | 31.5 | 58.5 | -18.5 | 5.1 |
| L nodule (lobule IX) |  | 2 | 833 | 3 | 13.5 | 55.5 | -24.5 | 4.7 |
| R declive (lobule VI) |  | 1 | 8400 | 238 | -34.5 | 67.5 | -21.5 | 5.5 |
| R culmen (lobule IV-V crus 1) |  | 1 | 8400 | 146 | -34.5 | 40.5 | -21.5 | 4.9 |
| L inferior parietal | 40 | 1 | 8400 | 138 | 52.5 | 37.5 | 50.5 | 4.2 |
| R inferior parietal | 40 | 1 | 8400 | 252 | -52.5 | 37.5 | 44.5 | 4.5 |
| L inferior temporal (fusiform) | 19, 37 | 2 | 833 | 109 | 46.5 | 64.5 | -15.5 | 6.4 |
| R inferior temporal (fusiform) | 37, 20 | 1 | 8400 | 149 | -43.5 | 55.5 | -12.5 | 6.9 |
| L cingulate (middle) | 32 | 1 | 8400 | 205 | 13.5 | -4.5 | 44.5 | 4.5 |
| R cingulate | 32 | 1 | 8400 | 205 | -22.5 | -10.5 | 29.5 | 4.6 |
| R anterior cingulate |  | 1 | 8400 | 107 | -22.5 | -25.5 | 26.5 | 5.4 |
| L superior temporal | 22 | 1 | 8400 | 50 | 61.5 | 1.5 | 8.5 | 4.7 |
| R superior temporal | 22 | 1 | 8400 | 33 | -61.5 | 4.5 | 8.5 | 5.1 |
| L substantia nigra |  | 4 | 32 | 6 | 10.5 | 19.5 | -9.5 | 3.8 |

**Table S3b.** Activation peaks of regions showing direction selectivity for Up TMSC under the Control-No-iNM condition.

| TT_Daemon Atlas Region | BA | Cluster | Cluster Size  (# of Voxels) | Region Size  (# of Voxels) | RAI Peak Coordinates (mm) | | | Peak  Z-score |
| --- | --- | --- | --- | --- | --- | --- | --- | --- |
|  |  |  |  |  | x | y | z |  |
| L precentral  (premotor & 1ary motor cortex) | 4, 6 | 3 | 791 | 406 | 55.5 | 4.5 | 20.5 | 8.0 |
| R precentral  (premotor & 1ary motor cortex) | 4, 6 | 4 | 776 | 342 | -61.5 | 4.5 | 20.5 | 7.3 |
| L postcentral  (1ary somatosensory) | 3, 1, 2 | 3 | 791 | 153 | 64.5 | 13.5 | 29.5 | 6.0 |
| R postcentral  (1ary somatosensory) | 3, 1, 2 | 4 | 776 | 191 | -49.5 | 13.5 | 53.5 | 3.8 |
| L thalamus |  | 12 | 11 | 5 | 10.5 | 16.5 | -0.5 | 4.5 |
| R thalamus |  | 11 | 12 | 9 | -13.5 | 16.5 | -0.5 | 5.0 |
| L lentiform & putamen |  | 8 | 127 | 61 | 25.5 | 1.5 | -6.5 | 7.2 |
| R lentiform & putamen |  | 7 | 140 | 49 | -25.5 | 1.5 | -6.5 | 6.7 |
| L insula & claustrum |  | 2 | 821 | 78 | 31.5 | -22.5 | 17.5 | 4.5 |
| R insula & claustrum |  | 4 | 776 | 99 | -34.5 | 7.5 | 14.5 | 6.0 |
| L inferior frontal | 10 | 2 | 821 | 70 | 43.5 | -40.5 | 2.5 | 3.2 |
| R inferior frontal | 47 | 1 | 1383 | 74 | -31.5 | -13.5 | -12.5 | 4.2 |
| L middle frontal | 8 | 2 | 821 | 118 | 25.5 | -10.5 | 35.5 | 4.3 |
| R middle frontal | 46 | 1 | 1383 | 104 | -28.5 | -43.5 | 11.5 | 5.2 |
| L medial frontal | 10, 46 | 2 | 821 | 167 | 19.5 | -46.5 | 11.5 | 5.1 |
| R medial frontal | 10 | 1 | 1383 | 232 | -22.5 | -37.5 | 17.5 | 4.7 |
| L superior frontal | 6, 8 | 2 | 821 | 142 | 16.5 | -16.5 | 50.5 | 4.6 |
| R superior frontal | 8 | 1 | 1383 | 156 | -10.5 | -25.5 | 44.5 | 4.6 |
| L declive  (superior vermian lobule VI) |  | 6 | 356 | 81 | 43.5 | 61.5 | -18.5 | 5.4 |
| L culmen (lobule IV-V) |  | 6 | 356 | 37 | 31.5 | 52.5 | -15.5 | 4.5 |
| R declive (lobule VI) |  | 5 | 447 | 120 | -34.5 | 58.5 | -15.5 | 5.4 |
| R culmen (lobule IV-V) |  | 5 | 447 | 64 | -31.5 | 40.5 | -18.5 | 3.7 |
| L inferior temporal (fusiform) |  | 2 | 833 | 63 | 43.5 | 58.5 | -15.5 | 5.2 |
| R inferior temporal (fusiform) |  | 1 | 8400 | 49 | -40.5 | 49.5 | -15.5 | 4.4 |
| L cingulate (middle) | 32, 6 | 1 | 1383 | 120 | 7.5 | 4.5 | 47.5 | 5.4 |
| R cingulate (middle) | 32 | 1 | 1383 | 170 | -16.5 | -16.5 | 32.5 | 5.1 |
| L superior temporal |  | 3 | 791 | 27 | 58.5 | 1.5 | 8.5 | 4.4 |
| R superior temporal |  | 4 | 776 | 68 | -61.5 | 7.5 | 8.5 | 4.6 |
| L amygdala |  | 8 | 127 | 14 | 25.5 | 1.5 | -9.5 | 6.3 |
| R amygdala |  | 7 | 140 | 23 | -25.5 | 1.5 | -9.5 | 6.2 |

**Table S4a.** Activation peaks of regions showing direction selectivity for Down TMSC under the iNM condition.

| TT_Daemon Atlas Region | BA | Cluster | Cluster Size  (# of Voxels) | Region Size (# of Voxels) | RAI Peak Coordinates (mm) | | | Peak  Z-score |
| --- | --- | --- | --- | --- | --- | --- | --- | --- |
|  |  |  |  |  | x | y | z |  |
| L precentral  (premotor & 1ary motor cortex) | 4, 6 | 1 | 5582 | 508 | 58.5 | 10.5 | 26.5 | 8.0 |
| R precentral  (premotor & 1ary motor cortex) | 4, 6 | 1 | 5582 | 427 | -49.5 | -4.5 | 8.5 | 4.9 |
| L postcentral  (1ary somatosensory) | 40 | 1 | 5582 | 198 | 58.5 | 28.5 | 20.5 | 3.5 |
| R postcentral  (1ary somatosensory) | 3/1/2 | 1 | 5582 | 263 | -58.5 | 13.5 | 29.5 | 7.4 |
| L thalamus |  | 4 | 190 | 42 | 10.5 | 16.5 | -0.5 | 5.5 |
| R thalamus |  | 4 | 190 | 56 | -10.5 | 16.5 | -0.5 | 6.2 |
| L lentiform |  | 1 | 5582 | 127 | 25.5 | 4.5 | -3.5 | 6.8 |
| R lentiform |  | 1 | 5582 | 109 | -25.5 | 1.5 | -6.5 | 6.2 |
| L insula & claustrum |  | 1 | 5582 | 218 | 31.5 | -16.5 | 5.5 | 6.8 |
| R insula & claustrum |  | 1 | 5582 | 194 | -31.5 | -19.5 | 8.5 | 5.7 |
| L inferior frontal (operculum) | 44 | 1 | 5582 | 218 | 55.5 | -4.5 | 23.5 | 6.6 |
| R inferior frontal (orbital) | 47 | 1 | 5582 | 260 | -52.5 | -16.5 | -0.5 | 4.1 |
| L middle frontal | 10 | 1 | 5582 | 83 | 28.5 | -37.5 | 11.5 | 4.7 |
| R middle frontal | 10 | 1 | 5582 | 180 | -43.5 | -49.5 | 11.5 | 3.0 |
| L medial frontal | 6 | 1 | 5582 | 168 | 7.5 | 4.5 | 50.5 | 6.0 |
| R medial frontal | 6 | 1 | 5582 | 268 | -4.5 | 7.5 | 53.5 | 5.7 |
| L superior frontal | 10 | 1 | 5582 | 84 | 31.5 | -43.5 | 14.5 | 4.5 |
| R superior frontal | 9 | 1 | 5582 | 180 | -16.5 | -46.5 | 23.5 | 4.6 |
| L uvula (crus 1 - lobule VII) |  | 3 | 845 | 33 | 28.5 | 64.5 | -24.5 | 4.7 |
| L nodule & ventral dentate  (lobule IX) |  | 3 | 845 | 27 | 13.5 | 55.5 | -24.5 | 4.2 |
| L culmen (crus 1 - lobule VI) |  | 3 | 845 | 99 | 43.5 | 43.5 | -27.5 | 3.7 |
| R culmen & ventral dentate  (crus 1 - lobule VI) |  | 2 | 1076 | 119 | -19.5 | 58.5 | -21.5 | 4.6 |
| L inferior parietal | 40 | 1 | 5582 | 71 | 49.5 | 52.5 | 47.5 | 3.7 |
| R inferior parietal | 40 | 1 | 5582 | 253 | -37.5 | 49.5 | 41.5 | 4.7 |
| R superior parietal | 7 | 1 | 5582 | 23 | -28.5 | 52.5 | 38.5 | 4.6 |
| L cingulate (middle) | 24, 32 | 1 | 5582 | 144 | 7.5 | -10.5 | 38.5 | 4.3 |
| R cingulate (middle) | 32 | 1 | 5582 | 178 | -7.5 | -16.5 | 38.5 | 5.0 |
| L anterior cingulate |  | 1 | 5582 | 34 | 13.5 | -31.5 | 23.5 | 3.6 |
| R anterior cingulate | 33 | 1 | 5582 | 46 | -4.5 | -10.5 | 23.5 | 4.3 |
| L superior temporal |  | 1 | 5582 | 100 | 37.5 | 1.5 | -21.5 | 3.5 |

**Table S4b.** Activation peaks of regions showing direction selectivity for Down TMSC under the Control-No-iNM condition.

| TT_Daemon Atlas Region | BA | Cluster | Cluster Size  (# of Voxels) | Region Size (# of Voxels) | RAI Peak Coordinates (mm) | | | Peak  Z-score |
| --- | --- | --- | --- | --- | --- | --- | --- | --- |
|  |  |  |  |  | x | y | z |  |
| L precentral  (premotor & 1ary motor cortex) | 3a | 1 | 909 | 420 | 52.5 | 4.5 | 23.5 | 7.7 |
| R precentral  (premotor & 1ary motor cortex) | 3a | 2 | 791 | 314 | -61.5 | 4.5 | 26.5 | 7.3 |
| L postcentral  (1ary somatosensory) | 3/1/2 | 1 | 909 | 149 | 58.5 | 13.5 | 26.5 | 6.7 |
| R postcentral  (1ary somatosensory) | 3/1/2 | 2 | 791 | 186 | -58.5 | 13.5 | 29.5 | 7.0 |
| L thalamus  (subthalamic nucleus) |  | 13 | 12 | 8 | 10.5 | 16.5 | -0.5 | 4.7 |
| R thalamus  (subthalamic nucleus) |  | 14 | 10 | 8 | -13.5 | 16.5 | -0.5 | 4.5 |
| L lentiform & putamen |  | 8 | 114 | 58 | 25.5 | 1.5 | -6.5 | 6.8 |
| R lentiform & putamen |  | 7 | 139 | 69 | -25.5 | 1.5 | -6.5 | 7.2 |
| L insula & claustrum |  | 1 | 909 | 146 | 34.5 | 7.5 | 14.5 | 5.2 |
| R insula & claustrum |  | 2 | 791 | 157 | -37.5 | 4.5 | 14.5 | 5.4 |
| L inferior frontal (operculum) | 47 | 12 | 17 | 100 | 37.5 | -31.5 | -6.5 | 4.1 |
| L middle frontal | 6, 8 | 11 | 42 | 47 | 19.5 | -19.5 | 53.5 | 4.3 |
| R middle frontal | 9 | 10 | 51 | 1 | -25.5 | -28.5 | 23.5 | 3.7 |
| L medial frontal | 9, 10 | 6 | 276 | 98 | 22.5 | -31.5 | 23.5 | 4.5 |
| R medial frontal | 9, 10 | 10 | 51 | 161 | -22.5 | -37.5 | 14.5 | 4.0 |
| L superior frontal | 10 | 6 | 276 | 91 | 31.5 | -43.5 | 14.5 | 4.6 |
| R superior frontal (posterior) | 9 | 3 | 461 | 67 | -16.5 | -49.5 | 23.5 | 4.4 |
| L uvula (lobule VII – crus 2) |  | 5 | 387 | 10 | 31.5 | 79.5 | -24.5 | 3.4 |
| R declive (lobule VI – crus 1) |  | 4 | 431 | 136 | -22.5 | 61.5 | -18.5 | 5.2 |
| R declive (lobule VI – crus 1) |  | 4 | 431 | 52 | -22.5 | 55.5 | -15.5 | 5.0 |
| L middle cingulate | 24 | 3 | 461 | 65 | 10.5 | 4.5 | 47.5 | 4.7 |
| R middle cingulate | 24 | 3 | 461 | 93 | -10.5 | 4.5 | 47.5 | 5.0 |
